# A rapid and universal liquid chromatograph-mass spectrometry-based platform, refmAb-Q nSMOL, for monitoring monoclonal antibody therapeutics

**DOI:** 10.1101/2022.04.22.489238

**Authors:** Noriko Iwamoto, Yoshinobu Koguchi, Kotoko Yokoyama, Akinobu Hamada, Atsushi Yonezawa, Brian D. Piening, Eric Tran, Bernard A. Fox, William L. Redmond, Takashi Shimada

**Author notes:** **Corresponding Author** Takashi Shimada, Ph.D., Shimadzu Scientific Instruments, Inc. Keihanna Research Laboratory, Shimadzu Corporation., 3-9-4 Seikacho-Hikaridai, Sorakugun, Kyoto 619-0237, Japan.

## Abstract

Accurate quantitation of antibody is critical for development of monoclonal antibody therapeutics (mAbs). Therapeutic drug monitoring has been applied to measure levels of mAbs in clinics for dose adjustment for autoimmune disease. Trough levels of mAbs can be a biomarker for cancer immunotherapy. Thus, the deployment of a rapid and universal platform for mAb monitoring may benefit processes ranging from drug development to clinical practice for a wide spectrum of diseases. However, mAb monitoring often requires development and conduct of an individual ligand binding assay such as ELISA, which is impractical to scale. We streamlined quantitation of antibody therapeutics by a nano-surface and molecular-orientation limited (nSMOL) proteolysis assay using LC-MS with a universal reference antibody (refmAb-Q), for accurate multiplexed quantitation of unique signature peptides derived from mAbs. This innovative refmAb-Q nSMOL platform may provide a practical solution for quantitating an ever-increasing number of mAbs from developmental to clinical use settings.

## Introduction

The US FDA approved the 100th therapeutic monoclonal antibody (mAb) in April 2021 and nearly 900 mAbs are currently in the pipeline^1,2^. These mAbs have been used for treating patients with inflammatory diseases, infectious diseases, and cancer. Drs. Allison and Honjo’s receipt of the Nobel Prize in Physiology or Medicine in 2018 for their work that led to the development of immune checkpoint inhibitor (ICI) therapy was one of many symbolic testimonies for the success of mAb-based therapy. Despite such promising prospects, mAb development is costly and time consuming, creating an additional financial burden for patients. Therefore, any solutions to streamline essential processes in drug development and facilitate proper use of mAbs are in great need.

Monitoring of mAb levels is critical for drug development to ensure safety and efficacy of mAbs^3-5^. Therapeutic drug monitoring (TDM), used for small molecule drugs, has been applied to adjust the doses of mAbs for patients with chronic inflammatory immune disease^6,7^ In addition, we and others have demonstrated that the trough levels of ICIs may serve as a potential biomarker for the effectiveness of cancer immunotherapy^8-13^. This is of particular note as current biomarkers for prediction of ICI response such as PD-L1 immunohistochemistry (IHC) and tumor mutational burden (TMB) profiling are imperfect, and a minority of treated individuals exhibit a significant tumor response to the therapy^14^. Thus, the deployment of a rapid and universal platform for mAb monitoring may provide benefits across various diseases and processes ranging from drug development to clinical practice.

A ligand-binding assay (LBA), namely enzyme-linked immunosorbent assay (ELISA), is currently the first choice for measuring mAbs. However, ELISA requires unique sets of reagents and assay optimization for each therapeutic mAb, which makes the adaptation and management impractical to cover an ever-increasing number of mAbs^15^. ELISA tends to suffer from poor sensitivity/selectivity at lower concentrations as well as non-specific binding as it is an indirect technique that relies on either a target antigen or an anti-idiotype antibody to capture mAbs. The presence of anti-drug Abs may also interfere with the binding of mAbs to the target antigen. Moreover, ELISAs are not suitable for multiplexing since it is impractical to prepared paired idiotypic antibodies for each mAb without causing any interference^16^. Even though planer and solution-based multiplexing applications are available, they suffer from lack of adequate standardization and quality control. In contrast, liquid chromatograph-mass spectrometry (LC-MS)-based mAb assays directly detect a structure unique to each mAb and therefore are compatible with multiplex quantitation^17^. However, typical LC-MS-based mAb assays have been plagued with analytical instability due to excess tryptic peptides and the trypsin enzymes since the whole analyte is digested^18,19^. We hypothesized that a technology capable of enriching the signature peptides should address such issues.

Previously, we developed nano-surface and molecular-orientation limited (nSMOL) proteolysis to overcome such challenges^20^. Briefly, IgGs are captured in a Protein A resin with a 100 nm pore. As a result, IgGs orient their Fab to the reaction solution. IgGs are proteolyzed by trypsin immobilized on the surface of nanoparticles with a 200 nm diameter. Immobilized trypsin has physicochemically limited access to the Fab of IgGs because of the difference of the two resins, therefore decreasing the peptide number while maintaining the structural specificity of complementarity-determining region (CDR) peptides for downstream multiple reaction monitoring (MRM) with triple-quadrupole LC-MS. Indeed, we have previously shown that nSMOL can be used to identify signature peptides of more than 30 unique mAbs in patient serum, confirming that the assay meets the criteria of the FDA Bioanalytical Method Validation Guidance for Industry (Table S1)^21^. However, the quantification of each mAb required the use of an authentic reference protein. This is impractical, for example, for a clinical laboratory to maintain unique assays for many different mAbs as each mAb assay requires at least 16 standards (duplicate of 8 standards). Thus, our goal was to develop an innovative assay based on nSMOL technology that could accurately measure concentrations of different mAbs using one universal reference antibody, named refmAb-Q.

## Materials and methods

### Experimental Design

The objective of the study was to develop a universal monitoring technology for mAbs using LC-MS-based assay. LC-MS is inherently suitable for multiplex and sequential analysis, but the matrix difference from each biological sample was a major hurdle to establish a reliable multiplexed assay. Since our nSMOL assay can overcome such challenge, we sought to examine a possibility of using universal reference control for multiplexed quantitation of mAbs. To this end, we conducted 18-plexed measurement of mAbs, and examined reproducibility of the CPS between trastuzumab, a universal reference candidate, and other mAbs. We also tried to understand the requirement for a good reference mAb by a hierarchical clustering analysis. Finally, we tried to validate the precision of the refmAb-Q nSMOL platform with clinical samples.

### Analysis workflow of measuring antibody concentration using the refmAb-Q nSMOL assay

For assessing the quantitative analysis of antibodies using a reference mAb, nSMOL assay (n=3) was first performed using the reference and target analyte mAbs with low QC and high QC concentrations within the linear quantitative range. The IS peptide P14R was used to correct the cps values of MS signals in each measurement. The averaged cps ratio of the reference antibody to the target analyte antibody was calculated using the IS area ratio of each. Next, a calibration curve for the reference antibody was prepared for the determination of its approximate linear regression. The concentration was calculated by substituting the averaged ratio of the reference to target analyte into the linear fitting curve. In the case of using the reference antibodies with different linear quantitative ranges, the concentration ratio was additively used as a coefficient based on the difference in concentration of each antibody (Equation below).

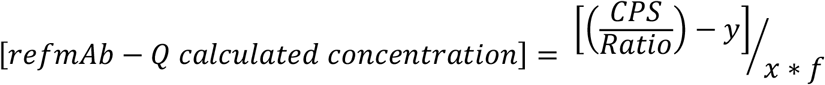

CPS: Peak intensity of signature peptide from analyte mAb

Ratio: CPS ratio of analyte mAb to reference mAb

y: intercept of linear regression curve created by reference mAb

x: slope of linear regression curve created by reference mAb

f: coefficient of quantitative range

(1: the quantifiable range from 1-200 μg/ml, 0.5: from 2-400 μg/ml, 0.4: from 2.5-500 μg/ml)

As a comparison group, the same mAbs of clinical samples (authentic mAbs) were used as a reference, and quantitative values were obtained by conventional methods using a calibration curve from the same reference (the conventional nSMOL assay). To evaluate the performance of refmAb-Q nSMOL assay, we first set the concentration at 50 μg/ml of mAb mixture in human serum for the identification of signature peptide structure by quadrupole time-of-flight (Q-TOF)-MS and the quantitative reproducibility by triple-quadrupole MS. The ratio of the reference mAb to target analyte mAb was set from 1 to 500 μg/ml samples in human serum and selected four concentrations from low to high QC samples.

### Physicochemical classification of signature peptides

To classify candidate mAbs based on their potential of serving as a reference mAb for the refmAb-Q, we consider physicochemical properties of signature peptides. MS detects ionized molecular signals as a mass/charge value. The mass number is invariant in a molecule-specific property, while the charge is defined by the probability of its electron presence. Molecules ionized by ESI have multiple charge states, among which the most abundant ions are preferentially selected and detected. Although the ionization reaction itself depends on the energy level of the molecule, it is difficult to clearly define the energy level especially in high MW biopolymers. In the case of peptides, a stable localization of the electron cloud is formed at the site of amide bonding. Thus, electron density and distribution of the amino acid residue likely defines the ionization reaction in a molecule-specific manner. The primary amines of peptides are the most likely proton acceptors. In addition, indole rings, some of aromatic rings, and sites that form globular structures can be also the proton acceptor. On the other hands, in the case of multivalent ion formation, intramolecular electrostatic repulsion is formed depending on the distance between the electron localizations, and the probability of the presence of the local electron site affects the most stable structure, size, and flattening. As a result, the molecular structure is expected to be affected by MW and multivalency. Therefore, we evaluated the physicochemical properties of the mAb signature peptides such as not only common factors in MS (the S/N ratio and hydrophobicity) but also the most stable energy level, the molecular diameter calculated with the most abundant electron valency, and the flattening ratio when the molecular surface area is approximated as an ellipse. Signature peptides have a wide range of MW, and peptide molecules become closer to a spherical shape dependent on decreasing of the size. Therefore, the factor per MW was also considered in the calculation of flattening. Hierarchical clustering was performed using BioVinci version 3.0.9 (San Diego, CA), and physicochemical properties were calculated by ChemOffice version 17.1 (PerkinElmer, Waltham, MA).

### Clinical sample information

Serum samples were obtained from patients with advanced melanoma who received ipilimumab^9,22^, pembrolizumab^8^, or combination of ipilimumab and nivolumab. All patients provided written informed consent and all studies were carried out in accordance with the Declaration of Helsinki under good clinical practice and Institutional Review Board approval.

### Statistical Analysis

Correlation analysis of nSMOL assay with the authentic reference and refmAB-Q nSMOL assay was performed using Prism version 7.05 (GraphPad Software, San Diego, CA).

## Results

### Developmental concept of refmAb-Q nSMOL assay

Monitoring of mAbs is not only critical for drug development but also holds a great promise in clinical settings through TDM or as a biomarker. In contrast to intensive investment in pharmacodynamic studies, monitoring of mAbs is rarely conducted outside pharmacokinetic (PK) study even though proper distribution and persistence of mAbs to the targeted organ(s) is likely prerequisite to clinical response. While ELISAs are the first choice for mAb assays, they require the development of an idiotypic antibody to capture mAbs and extensive assay-specific optimization (Figure S1A, top, and Table S2). ELISAs also suffer from a relatively narrow dynamic range, interference from anti-drug antibody, and cross-reactivity. A MS-based assay has been developed to directly detect the signature peptide of mAbs (Figure S1A, bottom), however it suffers from matrix effects related to trypsin digestion of whole antibody and the trypsin enzyme itself, which produces excess peptide fragments that interfere with reliable detection of signature peptides.

To leverage the strength of LC-MS-based approach but eliminate associated matrix effect issues, we have developed the nSMOL platform. The process is based upon selective proteolysis of antibody variable regions by maintaining structural specificity while decreasing undesired tryptic peptides through facilitating interaction between outward-oriented mAbs on protein A capture beads and catalytic trypsin-immobilized beads (Figure S1B and Table S2). Since our original report^20^, we have demonstrated successful analytical validation for over 40 mAbs (3 chimeric, 10 humanized, 11 human, 4 Fc-fusion, 3 antibody-drug conjugate, 1 scFv, 5 biosimilar, 6 mouse/rat mAbs) using the same standardized reagents and protocol regardless of analyte mAbs. Comparing with the “gold-standard” ELISA, the nSMOL assay achieved a wider dynamic range as well as a linear standard curve and was unperturbed by the presence of anti-drug antibody^23^. On top of these favorable analytical attributes, we have standardized assay development processes using Q-TOF-MS and other informatics tools (e.g., Skyline of MacCoss Lab^24^) to determine a target signature peptide within a week even if mAb peptide sequence is not available (Table S2).

Our success with the nSMOL assay inspired us to scale the assay to measure multiple mAbs in one assay run (Figure S1C and Table S2). This feature is critical for ensuring practical implementation of the nSMOL assay in a PK study, TDM assay, or biomarker study for multiple mAbs. We have already confirmed that the nSMOL assay protocol is compatible with automation. In this report, we sought to determine whether we could use one universal mAb reference to generate a standard curve to avoid excessive sets of authentic antibodies as standards for each antibody when analyzing a few samples for each mAb of interest in one assay run.

To establish refmAb-Q nSMOL, we redesigned the nSMOL assay with substantially enhanced features for multiplex quantification of mAbs with one universal reference mAb (Scheme). We harnessed the nSMOL chemistry, which uses 5 μL of unfractionated serum/plasma as a starting material for intact IgGs to detect the discrete mAb-specific signature peptide derived from the CDR. This feature enabled universal adaptation of the nSMOL protocol regardless of analyte mAbs if they bind to Protein A. We also introduced an antibody sensitizing (structural relaxing on H-chain) step with a reductive acidic pretreatment (pH 2) in addition to conventional non-acidic conditions to improve trypsin reaction of locally trypsin-resistant mAbs (∼25% of mAbs) for optimal recovery of signature peptides^25^. Based on these conditions, we aimed to establish a simplified standard curve-based extrapolation of mAb concentrations by using one universal reference mAb since the peak intensity ratio of each signature peptide can be established during MRM.

### Trastuzumab and other antibodies can be used as a reference antibody for the refmAb-Q nSMOL assay

To select a universal reference antibody for the refmAb-Q nSMOL assay, we picked 18 FDA-approved mAbs (4 chimeric, 7 humanized, and 7 human mAbs) and compared detection reproducibility of each mAb from 18 mAb spiked-in serum samples between singleplex and multiplex nSMOL assays. Signature peptides were previously identified and were derived from various location within the variable region of mAb (2 H-CDR1, 5 H-CDR2, 4 H-CDR3, 1 L-CDR1, 3 L-CDR2, and 3 L-CDR3: Figure S2 and Table S1). MS/MS spectra of mAb-derived signature peptides (Figure S3) were also adequate to assign the sequence determination with a good database search score using Q-TOF-MS (Table S3). Next, we confirmed that the separation was sufficient for multiplex quantitation with acceptable chromatogram overlaps within 3 min gradients with adding two more Fc-fusion proteins in the assay (total 20 analytes: 18 mAbs and 2 Fc-fusion proteins), and that each component can be switched by the MRM transition (Figure 1A). The ratio of multiplex assay to each singleplex nSMOL assay was equivalent, both in absolute and relative (P14R internal standard) values. All inter-assay errors for all antibodies were almost within 10%, demonstrating that the count per second (CPS) of each mAb is intrinsic, and that all mAbs are candidates for a potential universal reference in refmAb-Q nSMOL assay (Figure 1B). Next, we asked whether we could quantify mAbs using the CPS value ratio between the signature peptide of each mAb and a certain universal reference mAb (a reference in refmAb-Q assay: Scheme) to simplify the multiplex assay. To test this, the CPS ratios between each signature peptide of mAb from lower limit of quantitation (LLOQ) to high-quality control samples (QC) concentrations were compared (Figure 2, Figure S4). The results showed that the CPS ratio to trastuzumab was unique to each mAb and consistent. There seems be a trend for the CPS of signature peptides of certain mAbs to be decreased at higher concentrations in 20-plex assay. This is probably due to the interference effect of the composition of the solution and the stabilizer in the drug substance, which may have resulted in a change of the total IgG amount compared with a single reaction. In clinical samples of peripheral circulating blood, such effect is considered to be negligible (Figure S4). We further identified the other 3 mAbs as candidate reference mAbs in addition to trastuzumab (preferential choice antibodies in Figure 3) based on the physicochemical size and flattening properties that mass spectra are dependent on (e.g., the energy level of peptides attributed from intramolecular electron density and distribution)^26^. On the other hand, the use of antibodies with low CPS (e.g., Ave or Rit) was inadequate as a universal reference mAb in low concentrations (Figure S5). The expanded choices of reference mAb gives more flexibility for mAb monitoring when using the refmAb-Q nSMOL platform.

**Figure 1.**
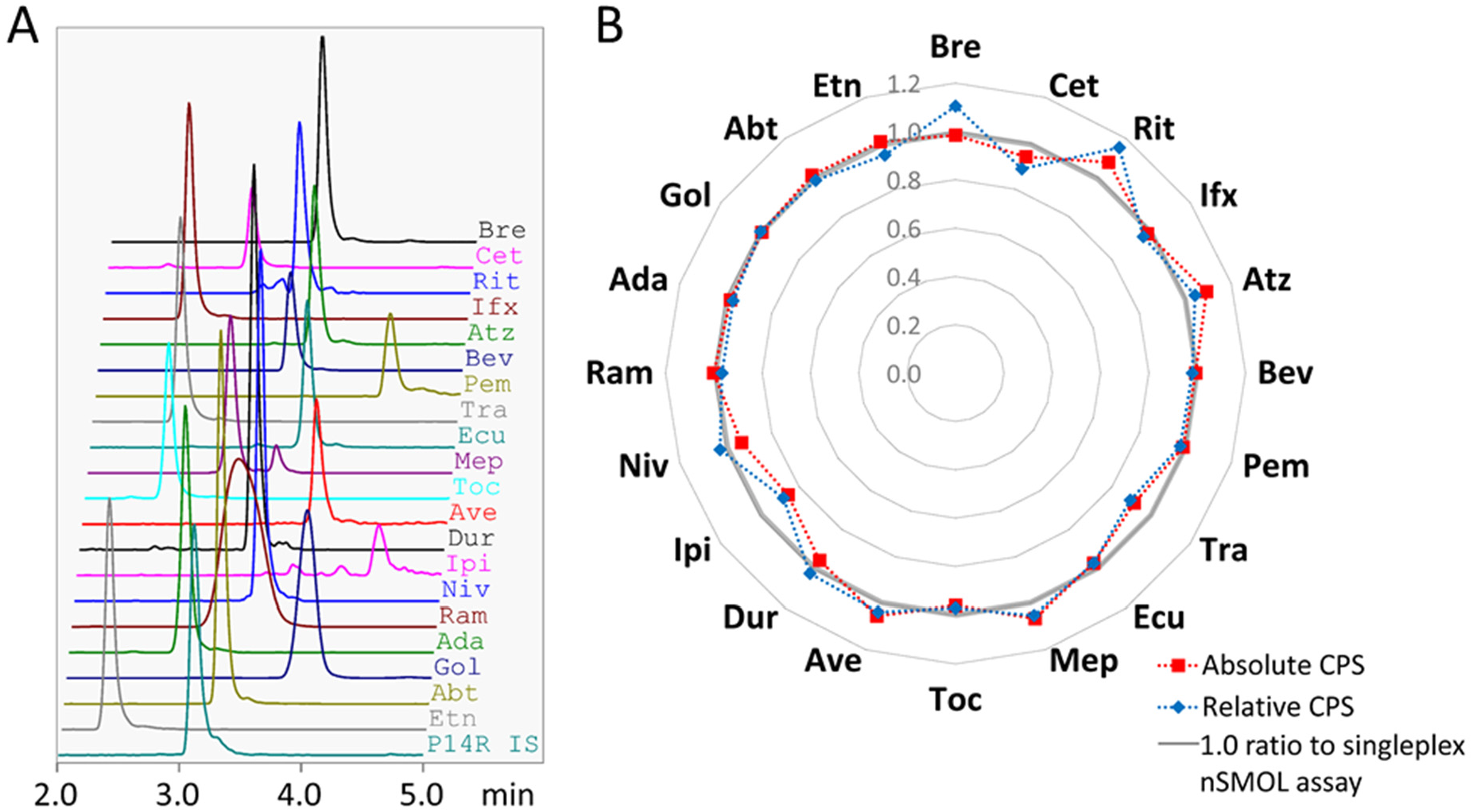
Performance and reproducibility of the multiplexed nSMOL assay. A) A representative 20-plex MRM chromatogram and chromatogram of P14R synthetic peptide for an internal correction. B) Averaged CPS ratio of the 20-plex nSMOL assay to the singleplex nSMOL assay for absolute and relative CPS value from each antibody signature peptide in human serum (n = 3).

**Figure 2.**
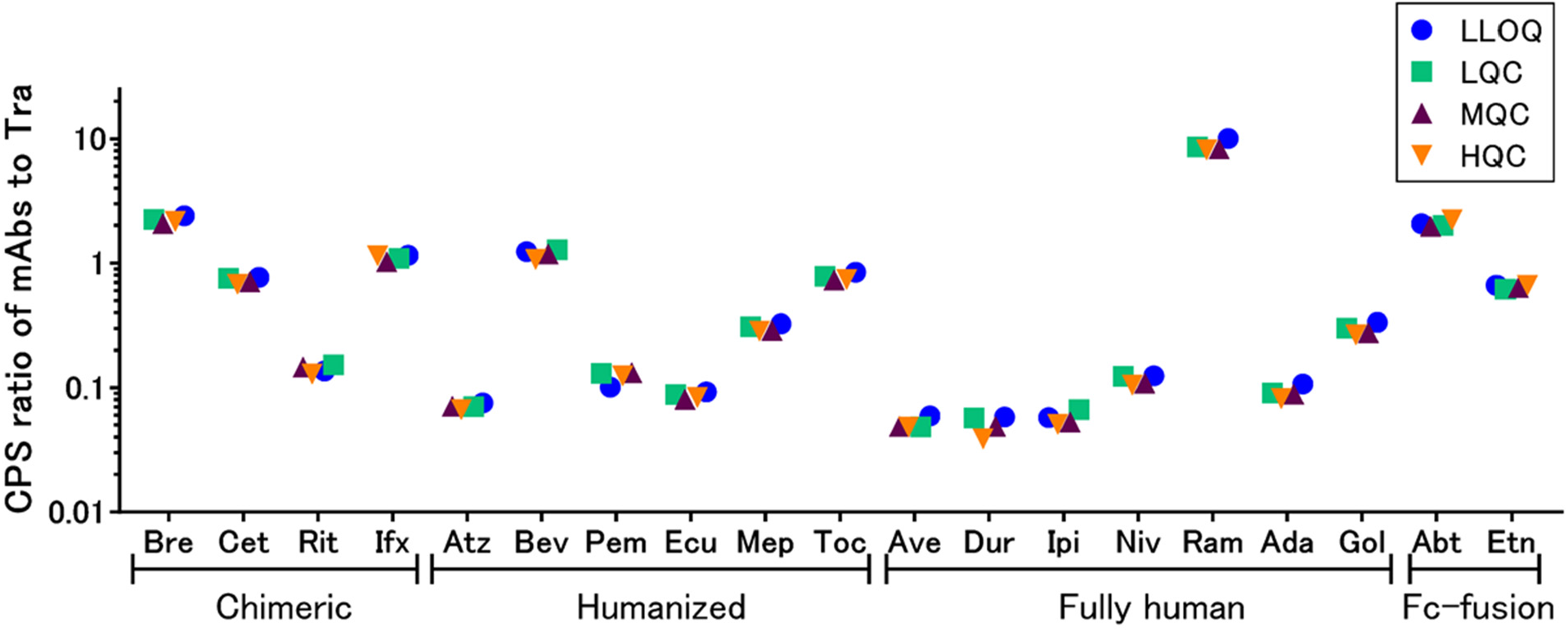
The consistent average CPS ratio of each signature peptide to reference. The average CPS ratio of each signature peptide of mAb to that of trastuzumab (Tra) reference across four different concentrations in human serum are shown (n=3). LLOQ: lower limit of quantification, LQC: low quality control samples, MQC: middle QC, HQC: high QC.

**Figure 3.**
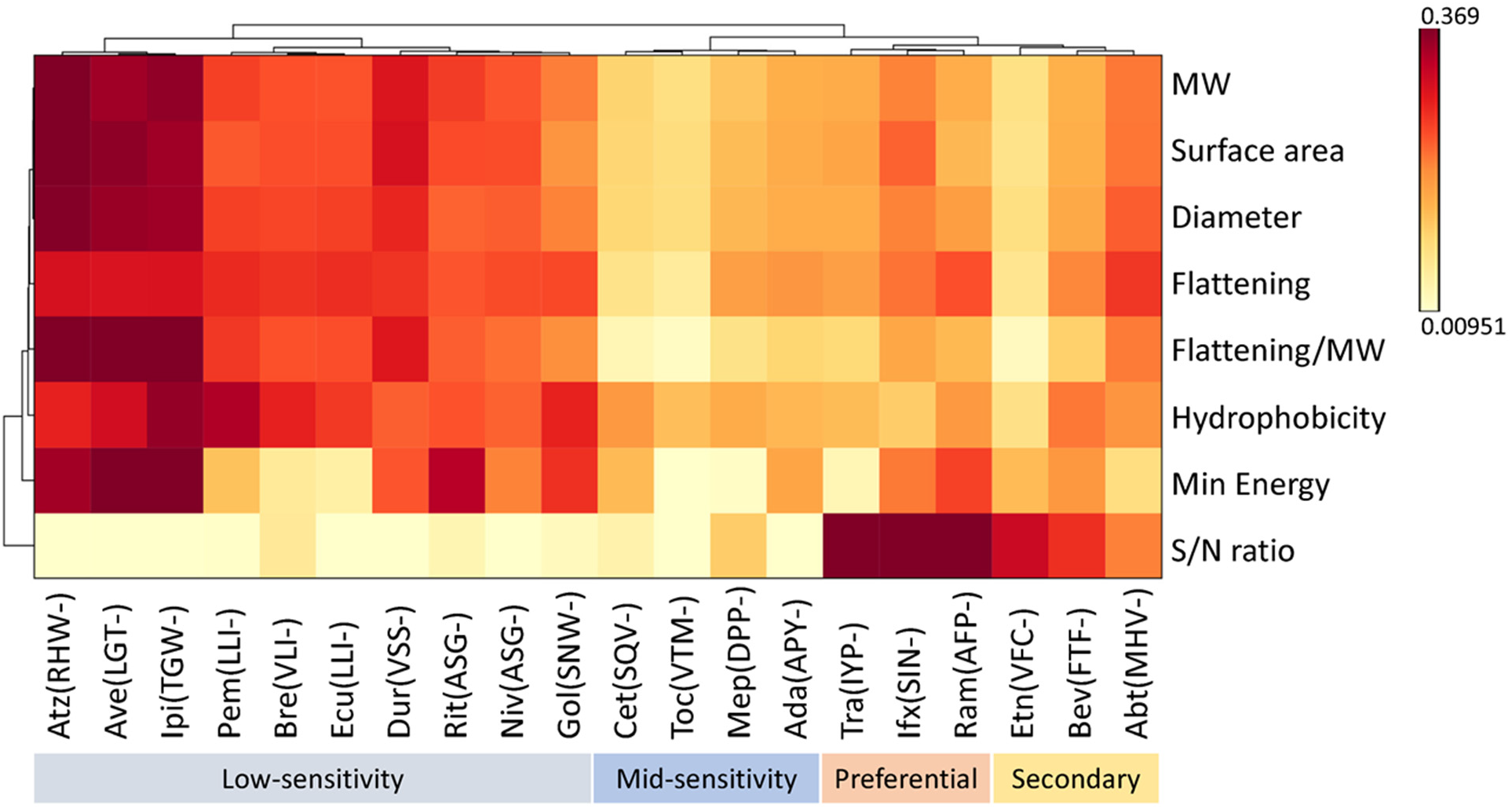
Classification of signature peptides suitable for reference. Hierarchical clustering distinguished preferential choice reference antibodies from other mAbs based on physicochemical properties, physicochemical size, and flattening properties of signature peptides. The preferential choice mAbs exhibited high sensitivity and S/N ratio. MW: molecular weight, S/N: signal to noise. The three N-terminal amino acids of each signature peptide are shown in parentheses.

The refmAb-Q nSMOL assay showed almost perfect concordance with the conventional nSMOL assay for detecting mAbs in clinical samples. To validate the refmAb-Q nSMOL platform for multiplexed quantification of mAbs with a reference antibody, we selected trastuzumab as a reference mAb and conducted refmAb-Q nSMOL assay for serum samples from ipilimumab-treated patients (n=71) and those from pembrolizumab-treated patients (n=74). We also used ipilimumab and pembrolizumab, respectively, as an authentic reference. We performed a Pearson correlation analysis between the result obtained by nSMOL assay with individual authentic reference and that with refmAb-Q nSMOL assay for the verification (Figure 4A, 4B, Table S4A, S4B). Samples close to the detection limit also showed almost perfect concordance (inserts in Figure 4). Finally, we confirmed that the refmAb-Q nSMOL assay was highly concordant with the conventional nSMOL assay with the authentic mAb reference when analyzing samples from patients receiving combination immunotherapy of ipilimumab and nivolumab in a multiplex setting (Figure 4C, 4D Table S4C, S4D). These results demonstrated that refmAb-Q nSMOL can be applied for antibody monitoring in clinical samples with highly sufficient reproducibility, including samples from patients receiving combination therapy.

**Figure 4.**
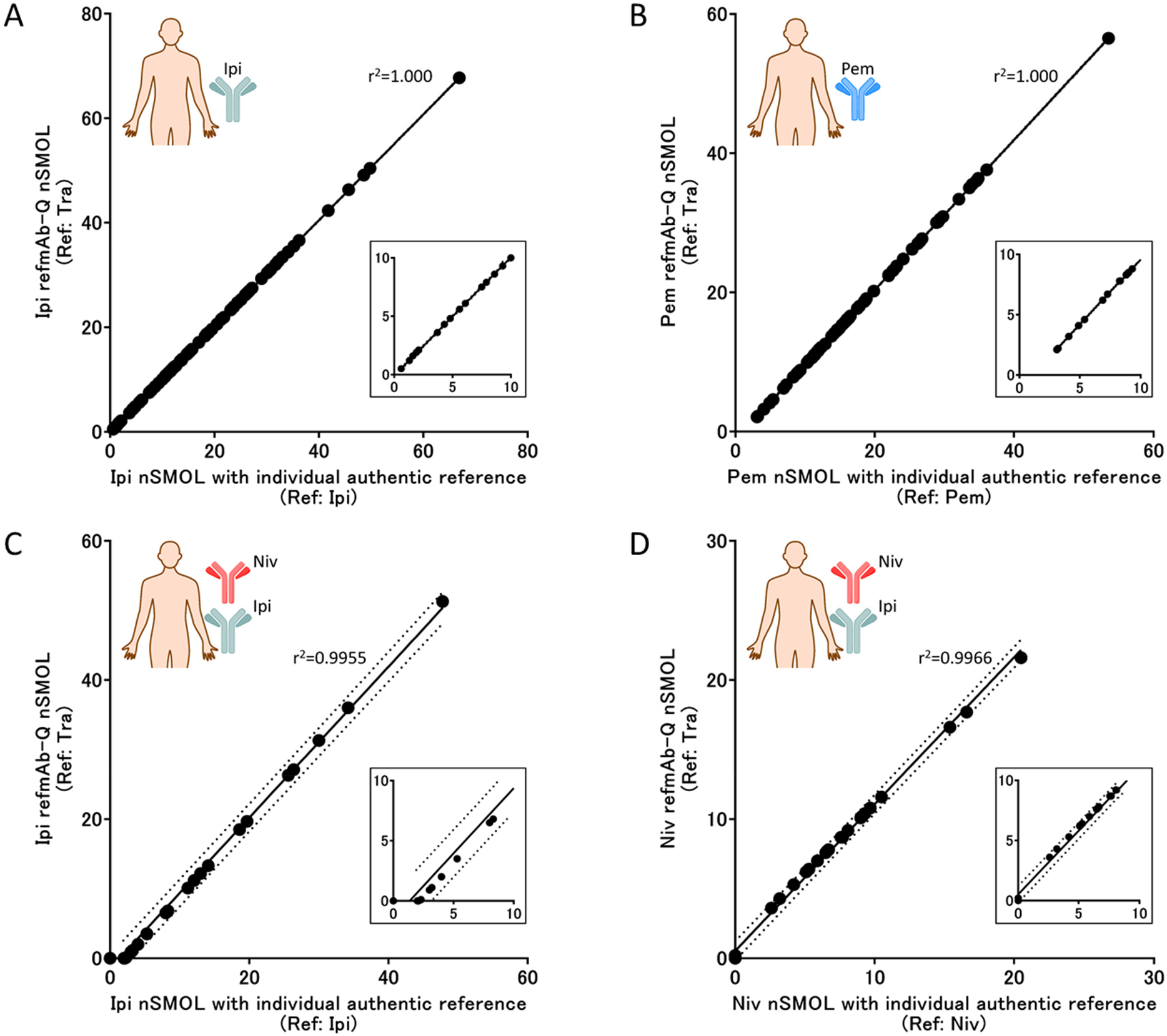
Performance of the refmAb-Q nSMOL assay for quantitating mAb levels of clinical samples. Assay concordance was examined between authentic mAb (x-axis) and refmAb-Q (Tra: y-axis) using sera from patients with advanced melanoma treated with A) ipilimumab (n = 71), B) pembrolizumab (n = 74), or C-D) ipilimumab plus nivolumab (n = 27). C) and D) depict detection of C) ipilimumab and D) nivolumab, respectively. The concordance in lower concentration is shown in the inserted box.

## Discussion

In this report, we demonstrated that trastuzumab can be used as a universal reference for quantitation of mAbs by the nSMOL assay, which is termed as the refmAb-Q nSMOL platform. We found excellent concordance between conventional nSMOL with an authentic antibody reference and refmAb-Q nSMOL with the trastuzumab universal reference. Finally, we demonstrated stellar performance of the refmAb-Q nSMOL assay for measuring clinical samples from patients receiving mAb monotherapy and combination therapy. Currently, more than 100 mAbs are clinically available and nearly 900 mAbs are in the pipeline^1^. TDM-based therapeutic guidance is becoming widely utilized in clinical practice for patients losing response to anti-TNF agents for inflammatory bowel diseases^27^. We and others showed that the trough levels of ICI mAb can be a biomarker for cancer immunotherapy^3,4,8-10^, this is of particular importance given the relatively low predictive capability of current biomarkers such as anti-PD-L1 IHC and TMB^14^. Therefore, development of a universal platform for quantitative analysis of mAbs is essential to streamline the PK study, TDM for effective use of mAbs, and biomarker discovery. The refmAb-Q nSMOL assay may serve as a prototype for the greater effort of developing a standardized platform of quantitation of mAbs to meet current challenges and needs described above.

An innovation of the refmAb-Q nSMOL assay is that it is a uniquely antibody-agnostic platform by establishing a universal reference standard with an analogous mAb instead of using corresponding authentic antibodies for the practical simultaneous quantitation of multiple mAbs. This was made possible because we found reproducible CPS ratios between the specific signature peptide of each target antibody and that of trastuzumab obtained by the nSMOL assay. In our previous studies, we showed that the proteolysis efficiency of antibodies on nSMOL reaction is almost 95% or higher^28^, and that the chemical equivalent of signature peptides produced from each biological sample (e.g., endogenous IgG and mAb) can be constant. Since the peak intensity of LC-MS depends on the peptide sequence and composition, the peak intensity obtained from each constant amount of peptide matrix has a constant physical quantity^29^. In other words, the peptides in the same amount can be determined using a constant ratio of peak intensity, which is the basic concept for refmAb-Q nSMOL technology. The refmAb-Q nSMOL inherits unique and beneficial features of nSMOL: 1) the nSMOL yields high sensitivity by recovering signature peptide(s) from the Fab region of mAbs for downstream LC-MS analysis without going through extensive sample preparation steps; 2) the nSMOL assay offers a mAb-agnostic solution by capturing mAbs and Fc fusion proteins with Protein A/G; 3) the assay reproducibility is not influenced by heterogeneity of subjects, samples (e.g., plasma or serum), or mAb concentrations; and 4) a biological matrix is constant and limited to the Fab peptides from total endogenous IgGs, which prevents physical adsorption of mAb-derived signature peptides to the sample vial, thus ensuring 48-hour analytical stability (manuscript in preparation). High-MW protein analysis with LC-MS is an advanced and complex technique that involves several multi-step physicochemical reactions (e.g., molecular fractionation, denaturation, proteolysis, separation, resin adsorption, elution, electrostatic interaction, electrospray ionization, vacuum control, coulomb repulsion, ion amplitude control, collision-induced dissociation, ion valency determination, and ion selection). The ability of LC-MS to directly assign the accurate peptide structures through molecular separation and MS/MS analysis is quite attractive. However, accurate quantitation, cost effectiveness, general usability, and reproducibility of all fragment ion intensities are still insufficient due to the protocol and technological complexity. Based on the stable and structure-indicated chemistry of signature peptide collection using Fab-selective proteolysis, our refmAb-Q nSMOL platform provides a breakthrough in quantitative reproducibility in measuring mAbs and general usability for various biological samples, which have long been considered as a weakness of MS.

Currently, a bottom-up approach is typically used for the quantitation of mAb by LC-MS, which relies on detection of a signature peptide^30^. The signature peptide was determined so that it provides excellent sensitivity and high signal to noise ratio when analyzed by LC-MS. In the analysis of mAbs in biological samples using LC-MS, there has been an active debate on how to correct analytical data using internal standards^31,32^. A stable isotope-labeled (SIL)-peptide of the signature peptide is most commonly used as an internal standard, which has been widely used in quantitative proteomics. However, there is no clear consensus regarding when the SIL-peptide should be added; the SIL-peptide was reported to be added before sample preparation^33,34^, before reduction^35,36^, before digestion^37,38^, or after digestion^39,40^. Moreover, the use of SIL-peptide may not correct for variations in LC-MS detection especially during steps before mAb digestion as the physical and chemical behavior of the signature peptide within the intact mAb and the SIL-peptide can be different. To overcome this shortcoming, SIL-mAbs have been introduced as an internal standard^41-43^. In fact, where more pre-digestion steps are required, an SIL-mAb yielded better results^44^. However, there are several issues in the use of SIL-mAbs: lot-to-lot errors, high cost, differences in recovery rate likely due to the structural/dynamic difference, main detectable peptides on Fc-loop, and lower stability than a pharmaceutical grade (e.g., presence of aggregates, denaturation). It is impractical to synthesize the SIL-mAb for ever increasing number of approved mAbs. In contrast, the nSMOL assay utilizes an authentic antibody (native and not SIL form) as a quantitation standard in separate reaction tubes just as the ELISA assay does. The P14R is used for the internal standard to correct measurement. Regardless of which type of quantitation standard is used, several dilutions of the quantitation standard are required for each mAb. To address this, we examined the utility of an analogous mAb in the replacement of authentic mAbs for the quantitation standard. Although the usefulness of analogous mAb as the internal standard has been examined by using alemtuzumab as an internal standard for quantitation of infliximab and bevacizumab, its use was limited to mAbs that have structurally similar signature peptides^45^, which falls short of a universal utility. In contrast, our refmAb-Q approach does not rely on structural similarity in signature peptides between reference and target mAbs. Instead, refmAb-Q relies on CPS ratio between trastuzumab and target mAbs, ensuring unrestricted utility. Trastuzumab and other preferential choice mAbs are readily and reliably available in a pharmaceutical grade. Together, the refmAb-Q, an approach of utilizing one analogous mAb as a universal reference for the nSMOL assay will address challenges in the bottom-up approach to quantitate mAbs.

Major advantages of the refmAb-Q nSMOL assay stem from its backbone, the nSMOL assay. We have applied and validated the nSMOL assay for more than 40 mAbs including 2 Fc fusion proteins^8,9,21,46-48^. The exquisite specificity of mAbs and availability of 6 CDR legions (3 from each heavy and light chains) enables robust selection of the signature peptide as we and others have demonstrated^30^. Therefore, we do not anticipate encountering difficulties in identifying a signature peptide as we expand our portfolio for the nSMOL method into the rest of FDA-and EMA-approved mAbs. The nSMOL assays showed excellent accuracy/recovery and precision (%CV < 10%). We showed great concordance between the nSMOL assay and ELISA or microfluidic-based LBA^49^. The lower LLOQ is one of the important hallmarks of the nSMOL assay. For example, in our direct comparison with bevacizumab, we found that the LLOQ for the nSMOL was 0.146 μg/ml whereas the LLOQ for conventional pellet digestion was 18.8 μg/ml^28^. Further, the best LLOQ was 0.06 μg/ml in the trastuzumab assay^50^. The nSMOL assay has an excellent linear quantitative range: the LLOQ of the nSMOL assay is typically lower than 1 μg/mL and it can measure mAbs at up to 100-300 μg/ml^51^. This feature curtails errors introduced through sample dilution and during the process of extrapolation of mAb concentration from the standard curve, while such errors are difficult to mitigate in ELISA (e.g., a linear standard curve is typically accompanied by narrower dynamic range whereas wider dynamic range ends up with a sigmoidal standard curve). Although others successfully measured mAbs using the nSMOL assay^47,52,53^ and we demonstrated the minimal inter-operator variabilities in measuring bevacizumab with the nSMOL assay^49^, we have built and validated a prototype for an automated nSMOL platform using an automated liquid handler in our effort to further address inter-operator variabilities (manuscript in preparation). We also showed reasonable inter-instrument/platform variabilities from the LC-MS when conducting the nSMOL assays. We compared two different triple quadrupole LC-MS instruments and found high concordance between them. As LC-MS was widely adapted in the clinical laboratory and pharmaceutical analysis over the last decade^54,55^, we are confident that a streamlined assay such as the refmAb-Q nSMOL with automated sample preparation will enable reproducible quantitation of mAbs from any stage of drug development through to the clinical care setting.

It is going to be increasingly important for quantitative assays of mAbs to be agnostic and to offer a multiplex capability as we and other showed that the antibody levels in circulation can be a biomarker for therapeutic efficacy and more antibody therapeutic are in clinical trials and in the pipeline. Among 100 mAbs approved by the US FDA, 42 of them are targeting 10 molecular interactions^1^. For instance, anti-TNF mAbs account for 4 antibodies (infliximab, adalimumab, certolizumab-pegol, golimumab). Etanercept also targets to neutralize TNF-α as a Fc fusion protein of TNFRII. These anti-TNF agents were approved for various inflammatory or autoimmune diseases. These patients can be treated with other mAb targeting IL-12/23p40, C5, or IL-6R at a medical center for autoimmunity and inflammation. Therefore, we developed multiplexed nSMOL assay to simultaneously detect 9 mAbs such as infliximab, adalimumab, ustekinumab, golimumab, eculizumab, etanercept, abatacept, tocilizumab, and mepolizumab in a single run and successfully detected mAbs from patient samples^41,51^. Others have also reported simultaneous detection of 2-7 mAbs^19,41,55-58^, further proving the feasibility of multiplexed detection of mAbs by LC-MS. Recent approvals from cancer immunotherapy to autoimmune disease treatment have involved several antibodies in the same target, and we expect to see this practice continue or perhaps increase in the number of unique mAbs. Specifically, there are an increasing number of immunotherapy combinations for treating advanced cancer patients. Some of these are combinations of two mAbs and are clinically approved (anti-PD-1/anti-CTLA-4), ready for regulatory review (anti-PD-1/anti-LAG-3), or actively pursued (e.g., anti-PD-1/anti-VEGF)^59^, in which cases the multiplex assay will enable simultaneous detection of mAbs. In addition, many bispecific antibodies are actively being pursued in the immune-oncology field^60,61^. The detection of intact bispecific antibodies requires a multiplex assay. Although further study is required to confirm the nSMOL assay can be used to quantitate bispecific antibodies, since the nSMOL assay is compatible even with Fc-fusion protein such as abatacept and etanercept, or with 3 CDR-peptide quantitation from bevacizumab^28^, we envision that the nSMOL assay can support the measurement of bispecific antibodies. Together, refmAb-Q nSMOL will provide a practical solution for quantitation of mAbs in the ever-changing landscape of mAb treatment.

Another important hallmark of the refmAb-Q nSMOL platform is its capability for rapid method development. Development of the nSMOL method for a new mAb only takes 1-2 weeks, while that of the LBA can take up to several months. This is because the nSMOL assay directly detects the signature peptide of the mAb whereas the LBA depends on successful development of an anti-idiotype antibody for mAb quantitation through indirect detection. Development of the nSMOL method for a new mAb is well streamlined as it only requires identification of the signature peptide, verification of the signature peptide in biological matrix, and establishment of LC and MS parameter^21^. Even though computational analysis does not completely replace the need for LC-MS analysis to identify the signature peptide, it aids in narrowing down the candidate signature peptides to be examined. Once the signature peptide is identified, it is necessary to confirm that there is no significant interference with detecting the signature peptide of the mAb of interest from endogenous antibodies contained in human sera. As mAbs are sufficiently different from endogenous antibodies, we and others have successfully verified many mAbs for LC-MS-based quantitation with minimum to no interference from endogenous antibodies. Establishment of LC and MS parameters can be achieved by applying a standard gradient setting for detecting tryptic peptides. Regarding the implementation of refmAb-Q nSMOL assay, one may need to establish the CPS ratio at least once with the instrument in use as ionization and detection efficiency of an analyte may differ among instruments. As refmAb-Q relies on the ratio between the universal reference and the mAb of interest, it is not influenced by day-to-day instrument performance variabilities. Together, the implementation of the refmAb-Q nSMOL may enable a time-effective and cost-effective operation for measuring mAbs.

## Conclusion

We envision that wide adaptation of refmAb-Q nSMOL is possible if adequate support is provided. The protocol detail for the nSMOL assay from identifying the signature peptide to conducting assays has been published^21^. Pharmaceutical companies can substantially simplify the development of PK studies for a new mAb. If they chose to disclose the signature peptide and LC-MS condition after approval for the clinical use, it would make it easier for end users to include a new mAb in their analysis portfolio. A community website storing and updating LC-MS condition for approved mAbs would serve as a great resource for PK and TDM assays for mAbs. Together, our refmAb-Q nSMOL assay has the potential to transform how we conduct PK and TDM assays for mAbs.

## Author information

## Author Contributions

The manuscript was written through contributions of all authors. All authors have given approval to the final version of the manuscript. ‡These authors contributed equally.

Conceptualization: NI, YK, TS, AH, AY, ET, BAF, BDP, WLR; Methodology: NI, KY, TS; Investigation: NI, YK, KY, TS; Supervision: TS; Writing original draft: NI, YK, TS; Writing review & editing: ET, BAF, BDP, WLR

## Scheme

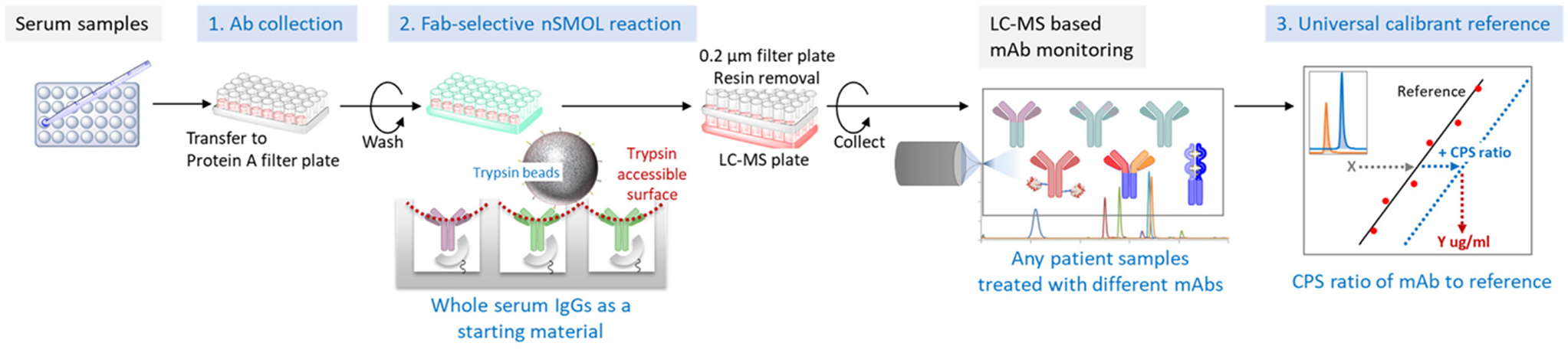
Workflow of the refmAb-Q nSMOL. A procedural workflow of the refmAb-Q nSMOL assay. The CPS ratio of an analyte mAb to a trastuzumab reference is used for the determination of concentration.

