## Supplementary information for "A rapid and universal liquid chromatograph-mass spectrometry-based platform, refmAb-Q nSMOL, for monitoring monoclonal antibody therapeutics"

### Supporting experimental section

### Reagents

The nSMOL Antibody BA kit (Shimadzu, Kyoto, Japan) was used for antibody analysis. Modified trypsin-immobilized glycidyl methacrylate (GMA)-coated nanoferrite particle FG beads with surface activation by N-hydroxysuccinimide were purchased from Tamagawa Seiki (Nagano, Japan). Toyopearl AF-rProtein A HC-650F resin was purchased from Tosoh Bioscience (Tokyo, Japan). Brentuximab vedotin was purchased from Takeda Pharmaceuticals (Osaka, Japan). Cetuximab was purchased from Merck Biopharma (Tokyo, Japan). Rituximab was purchased from Zenyaku Kogyo (Tokyo, Japan). Infliximab and golimumab were purchased from Mitsubishi Tanabe Pharma (Osaka, Japan). Atezolizumab, bevacizumab, trastuzumab, and tocilizumab were purchased from Chugai Pharmaceutical (Tokyo, Japan). Pembrolizumab and avelumab were purchased from Merck (Kenilworth, NJ, USA). Eculizumab was purchased from Alexion Pharmaceuticals (Boston, MA, USA). Mepolizumab was purchased from GlaxoSmithKline (Brentford, UK). Durvalumab was purchased from AstraZeneca (Cambridge, UK). Ipilimumab, nivolumab, and abatacept were purchased from Bristol Myers Squibb (New York, NY, USA). Ramucirumab was purchased from Eli Lilly (Indianapolis, IN, USA). Adalimumab was purchased from AbbVie (North Chicago, IL, USA). Etanercept was purchased from Pfizer (New York, NY, USA). Control human serum was obtained from Innovative Research (Novi, MI). The synthetic peptide P14R (14 Pro and Arg), octyl-β-D-glucopyranoside (OTG), organic solvents, and other reagents were obtained from Sigma-Aldrich (St. Louis, MO).

**Structural confirmation of signature peptides**

Multiple sequence alignment analysis was performed using the ClustalW version 2.1 algorithm on GENETYX software version 13 (GENETYX, Tokyo, Japan) for the prediction of the signature peptides of the antibodies in the complementarity-determining regions (CDR1, 2, and 3) and specific peptides for Fc-fusion proteins in the fused domain. In this analysis, theoretical tryptic peptides with no overlaps in the sequence of immunoglobulin Fc, hinge, and the conserved positions of cysteine residues and S-S bonds were aligned and predicted as good signature peptides.

Two micrograms of each antibody and Fc-fusion proteins were denatured and reduced in 8 M urea and 2 mM neutralized Tris(2-carboxyethyl)phosphine (TCEP) at room temperature for 30 min. Next, the sample solution was diluted 1:10 in a 25 mM Tris-HCl solution (pH

8.0) and digested using modified trypsin at 37 °C for 16 h. The proteolytic reaction was quenched by the addition of a trifluoroacetic acid (TFA) solution at a final concentration of 0.5%. The peptide solution was purified using a MonoSpin C18 spin filter (GL Science, Tokyo, Japan). The eluting solution was evaporated in a centrifugal evaporator and reconstituted in 0.1% formic acid (FA) solution. The structures of the tryptic peptides from each protein were analyzed using microflow high-resolution liquid chromatography-quadrupole ion trap time-of-flight (Q-TOF) MS (Nexera Mikros high-performance microflow liquid chromatograph and LCMS-9030, Shimadzu). The LC-MS conditions were as follows: solvent A, 0.1% aqueous formic acid; solvent B, 80% acetonitrile with 0.1% formic acid; trap column, Triart Capillary trap column, 0.3 × 5 mm, 5 µm, 12 nm pore (YMC, Kyoto, Japan); separation column, L-column2 ODS, 0.3 × 150 mm, 2 µm, 12 nm pore (Chemicals Evaluation and Research Institute, Tokyo, Japan); column temperature, 40 °C; flow rate, 5 µL/min; gradient program, 0-10 min: %B=0, 10-95 min: %B=0-40 gradient, 95-105 min: %B=70-100 gradient, 100-115 min: %B=100, 115-130 min: %B=0. MS and MS/MS spectra were obtained using a desolvation line, interface, and heat block at 200, 100, and 250 °C, respectively. Nebulizer nitrogen gas flow was set at 1 L/min. The heating gas flow rate was 3 L/min. The electrode of the electrospray ionization (ESI) interface was set to

3 kV. The pulse times for MS and MS/MS were 194 and 154 μs, respectively. The ion accumulation time was set at 100 ms for MS and 80 ms for MS/MS. MS/MS analysis was performed using the intensity-dependent top-8 MS/MS per scan on data-dependent acquisition (DDA). The precursor MS was set from m/z = 300 to 1,500, and the fragments were set from m/z = 200 to 1,500. The ion valency was set from +2 to +6. The electrode of the collision-induced dissociation (CID) cell was set at -25 ± 5 V, and the argon gas pressure was set at 250 kPa. The precursor and fragment ions were assigned using Mascot Proteome Server version 2.6.2, Distiller peak processing software (Matrix Science, London, UK) and PEAKS Studio version X/X+ software (Bioinformatics Solutions, Waterloo, Canada) on SwissProt sequence database version 2020_04 and the in-house FASTA database for monoclonal antibody and Fc-fusion protein sequence information obtained from KEGG (Kyoto Encyclopedia of Genes and Genomes of Kyoto University Bioinformatics Center, Kyoto, Japan) DRUG Database in GenomeNet and DrugBank (The Metabolomics Innovation Center, University of Alberta, Canada). The allowance of peptide m/z tolerance on the database search was set to within 10 ppm for precursor ions and 20 mDa for fragment ions.

**Setting the conditions for the multiple reaction monitoring (MRM) of each signature peptide**

The peptide quantitation was conducted using an LC-MS with triple quadrupole (Nexera X2 and LCMS-8050, Shimadzu). The LC-MS conditions were as follows: solvent A, 0.1% aqueous formic acid; solvent B, acetonitrile with 0.1% formic acid; column, Shim-pack GISS C18, 2.1 × 50 mm, 1.9 µm, 20 nm pore (Shimadzu); column temperature, 50 °C; flow rate, 0.4 mL/min; gradient program, 0-1 min: %B=1, 1-6 min: %B=1-50 gradient, 6-7.5 min: %B=95, 7.5-9 min: %B=1. MS spectra were obtained using an ESI probe temperature, desolvation line, and heat block at 300 °C, 250 °C, and 400 °C, respectively. Nebulizer, heating, and drying gas flows were set to 3 L/min, 10 L/min, and 10 L/min, respectively. The dwell time was set to 10 ms for each MRM transition. The MRM monitor ions of the peptide fragments were determined from the measured values of the structure-assigned fragments from the Q-TOF-MS analysis. The CID argon partial pressure in the Q2 cell was set to 270 kPa. The candidate MRM transition m/z was computationally set, and the electrode voltage of the Q1 pre bias, collision cell Q2, Q3 pre bias, and the most abundant m/z of the precursor and fragment ions were performed using the optimization support software (LabSolutions, Shimadzu). The most abundant MRM transition with good linearity and no interference in human serum was selected for quantitation, and the second and third transitions were used for the structural confirmation of each peptide.

**Preparation of antibody analysis using nSMOL proteolysis**

In previous reports, we performed a bioanalytical validation for many monoclonal antibodies and Fc-fusion proteins in human serum/plasma using the nSMOL assay (Table S1). In this study, all assays were performed using our previously described methods, with minor modifications. Briefly, all sample sets in human serum were prepared and stored at -80 °C for at least 24 h before each nSMOL assay. Antibody-spiked human serum was prepared as an individual or 20-mixed spike. A 5 μL aliquot of antibody-spiked human serum was diluted 1:10 in PBS (pH 7.4) containing 0.1% OTG to avoid non-specific binding to the resin and plastic materials. The IgG fraction from the plasma sample was collected using 25 μL of PBS-substituted immunoglobulin collection Protein A resin slurry (25%) in 100 μL of PBS containing OTG with gentle vortexing at 25 °C for 10 min on a Unifilter filtration plate (Whatman, Maidstone, UK). Serum samples were prepared in duplicate; one was directly subjected to a resin washing step and the other was subjected to the washing process after pretreatment with acidic 250 mM TCEP-HCl (<pH 2) for 30 min at room temperature. The

collection resin was mixed, washed twice with 200 μL of PBS containing 0.1% OTG to remove other serum proteins except for IgGs, and washed again with 200 μL of PBS to remove the detergents that can inhibit column separation and ionization of peptides in the ESI interface and cause carryover. Each washing substitution was performed by centrifugation (100 × g for 2 min) on filter plates. After these washing steps, the collection resin was suspended with 80 μL of the nSMOL-enhanced reaction solution containing the P14R internal standard (IS), and the suspension was immediately transferred onto a Protein LoBind plate (Eppendorf, Hamburg, Germany), repeated twice for completely collecting the suspended resin. nSMOL proteolysis was carried out using 10 μg trypsin on FG-beads with gentle vortexing at 50 °C for 5 h in a saturated vapor atmosphere for uniform contact between the collection resin and the FG bead nanoparticles. After nSMOL proteolysis, the reaction was quenched by adding 10% formic acid at a final concentration of 0.5%. The peptide solution was collected by centrifugation (1,500 × g for 10 min) on an AcroPrep Advance PTFE 0.2-μm filter plate (Pall, New York, NY) to remove the collection resin and the trypsin FG-beads. These filtered analytes were collected in an overlapped low-protein binding Microresico plate (Richell, Toyama, Japan).

### Supporting Figures and Tables

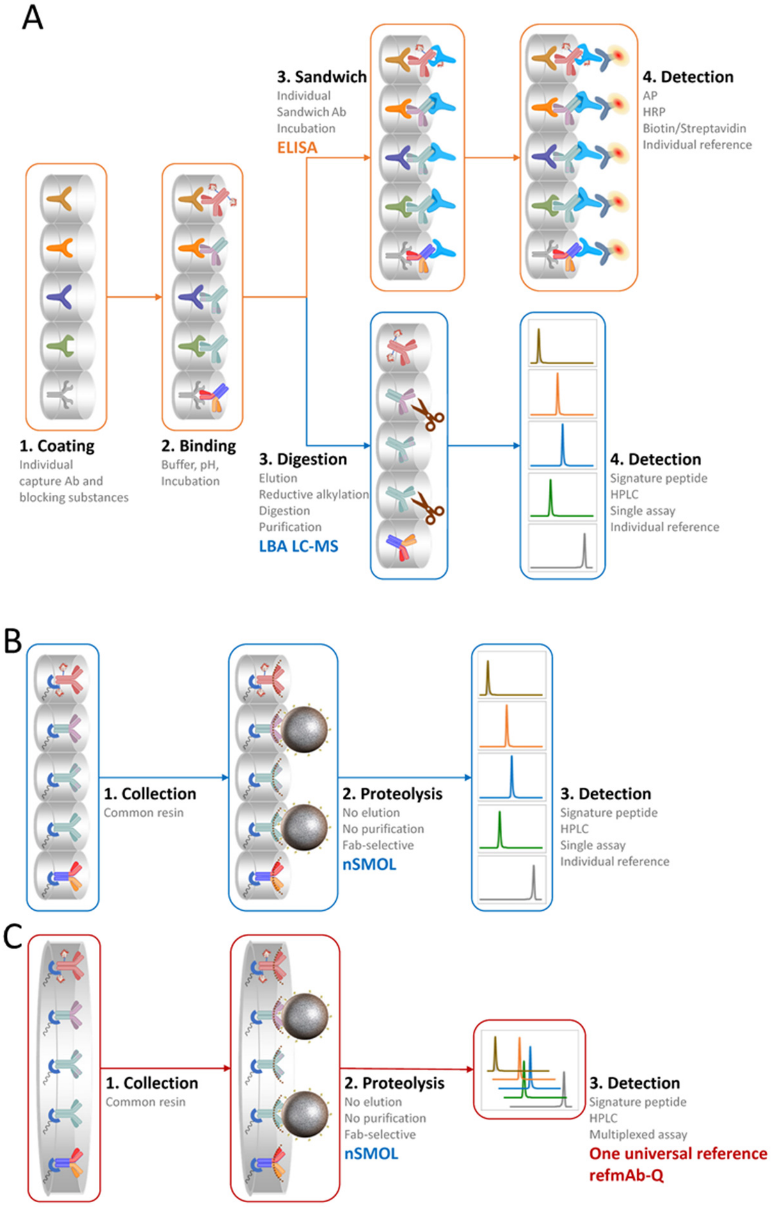


**Figure S1. Establishment of the refmAb-Q nSMOL.** A) A diagram of the LBA-based mAb assay. B) A diagram of the conventional nSMOL mAb assay method. C) A diagram of the refmAb-Q nSMOL mAb assay by redesigning the conventional nSMOL assay for MS-directed technique.

**A**

```
Bre H   1:QIQLQQSGPEVVKPGASVKISCKASGYTFTD-YYITWVKQKPGQGLEWIGWIYPG--SGN 57
Cet H   1:QVQLKQSGPGLVQPSQSLSITCTVSGFSLTN-YGVHWVRQSPGKGLEWLGVIWS---GGN 56
Rit H   1:QVQLQQPGAELVKPGASVKMSCKASGYTFTS-YNMHWVKQTPGRGLEWIGAIYPG--NGD 57
Ifx H   1:EVKLEESGGGLVQPGGSMKLSCVASGFIFSN-HWMNWVRQSPEKGLEWVAEIRSKSINSA 59
Atz H   1:EVQLVESGGGLVQPGGSLRLSCAASGFTFSD-SWIHWVRQAPGKGLEWVAWISPY--GGS 57
Bev H   1:EVQLVESGGGLVQPGGSLRLSCAASGYTFTN-YGMNWVRQAPGKGLEWVGWINTY--TGE 57
Pem H   1:QVQLVQSGVEVKKPGASVKVSCKASGYTFTN-YYMYWVRQAPGQGLEWMGGINPS--NGG 57
Tra H   1:EVQLVESGGGLVQPGGSLRLSCAASGFNIKD-TYIHWVRQAPGKGLEWVARIYPT--NGY 57
Ecu H   1:QVQLVQSGAEVKKPGASVKVSCKASGYIFSN-YWIQWVRQAPGQGLEWMGEILPG--SGS 57
Mep H   1:QVTLRESGPALVKPTQTLTLTCTVSGFSLTS-YSVHWVRQPPGKGLEWLGVIWA---SGG 56
Toc H   1:EVQLQESGPGLVRPSQTLSLTCTVSGYSITSDHAWSWVRQPPGRGLEWIGYISY---SGI 57
Ave H   1:EVQLLESGGGLVQPGGSLRLSCAASGFTFSS-YIMMWVRQAPGKGLEWVSSIYPS--GGI 57
Dur H   1:EVQLVESGGGLVQPGGSLRLSCAASGFTFSR-YWMSWVRQAPGKGLEWVANIKQD--GSE 57
Ipi H   1:QVQLVESGGGVVQPGRSLRLSCAASGFTFSS-YTMHWVRQAPGKGLEWVTFISYD--GNN 57
Niv H   1:QVQLVESGGGVVQPGRSLRLDCKASGITFSN-SGMHWVRQAPGKGLEWVAVIWYD--GSK 57
Ram H   1:EVQLVQSGGGLVKPGGSLRLSCAASGFTFSS-YSMNWVRQAPGKGLEWVSSISSS--SSY 57
Ada H   1:EVQLVESGGGLVQPGRSLRLSCAASGFTFDD-YAMHWVRQAPGKGLEWVSAITWN--SGH 57
Gol H   1:QVQLVESGGGVVQPGRSLRLSCAASGFIFSS-YAMHWVRQAPGNGLEWVAFMSYD--GSN 57

Bre H  58:TKYNEKFKGKATLTVDTSSSTAFMQLSSLTSEDTAVYFCAN---YG-NYWFA-----YWG 108
Cet H  57:TDYNTPFTSRLSINKDNSKSQVFFKMNSLQSNDTAIYYCARALTYY-DYEFA-----YWG 110
Rit H  58:TSYNQKFKGKATLTADKSSSTAYMQLSSLTSEDSAVYYCARSTYYGGDWYFN-----VWG 112
Ifx H  60:THYAESVKGRFTISRDDSKSAVYLQMTDLRTEDTGVYYCSRNY---YGSTY-----DYWG 111
Atz H  58:TYYADSVKGRFTISADTSKNTAYLQMNSLRAEDTAVYYCARRH---WPG-----GFDYWG 109
Bev H  58:PTYAADFKRRFTFSLDTSKSTAYLQMNSLRAEDTAVYYCAKYPHYYGSSHWY---FDVWG 114
Pem H  58:TNFNEKFKNRVTLTTDSSTTTAYMELKSLQFDDTAVYYCARRDYRFDMG------FDYWG 111
Tra H  58:TRYADSVKGRFTISADTSKNTAYLQMNSLRAEDTAVYYCSRWG---GDGFY---AMDYWG 111
Ecu H  58:TEYTENFKDRVTMTRDTSTSTVYMELSSLRSEDTAVYYCAR--YFFGSSPNW--YFDVWG 113
Mep H  57:TDYNSALMSRLSISKDTSRNQVVLTMTNMDPVDTATYYCARDPPSS-LLRLD-----YWG 110
Toc H  58:TTYNPSLKSRVTMLRDTSKNQFSLRLSSVTAADTAVYYCARSLAR--TTAMD-----YWG 110
Ave H  58:TFYADTVKGRFTISRDNSKNTLYLQMNSLRAEDTAVYYCARIK-LGTVTTV-----DYWG 111
Dur H  58:KYYVDSVKGRFTISRDNAKNSLYLQMNSLRAEDTAVYYCAREGGWFGELAF-----DYWG 112
Ipi H  58:KYYADSVKGRFTISRDNSKNTLYLQMNSLRAEDTAIYYCARTGWLGP--------FDYWG 109
Niv H  58:RYYADSVKGRFTISRDNSKNTLFLQMNSLRAEDTAVYYCATN-------------DDWYG 104
Ram H  58:IYYADSVKGRFTISRDNAKNSLYLQMNSLRAEDTAVYYCAR-----VTDAF-----DIWG 107
Ada H  58:IDYADSVEGRFTISRDNAKNSLYLQMNSLRAEDTAVYYCAKVSYLSTASSL-----DYWG 112
Gol H  58:KKYADSVKGRFTISRDNSKNTLYLQMNSLRAEDTAVYYCARDRGIAAGGNYYYYGMDVWG 117

Bre H 109:QGTQVTVSAASTK                                                121
Cet H 111:QGTLVTVSAASTK                                                123
Rit H 113:AGTTVTVSAASTK                                                125
Ifx H 112:QGTTLTVSSASTK                                                124
Atz H 110:QGTLVTVSSASTK                                                122
Bev H 115:QGTLVTVSSASTK                                                127
Pem H 112:QGTTVTVSSASTK                                                124
Tra H 112:QGTLVTVSSASTK                                                124
Ecu H 114:QGTLVTVSSASTK                                                126
Mep H 111:RGTPVTVSSASTK                                                123
Toc H 111:QGSLVTVSSASTK                                                123
Ave H 112:QGTLVTVSSASTK                                                124
Dur H 113:QGTLVTVSSASTK                                                125
Ipi H 110:QGTLVTVSSASTK                                                122
Niv H 105:QGTLVTVSSASTK                                                117
Ram H 108:QGTMVTVSSASTK                                                120
Ada H 113:QGTLVTVSSASTK                                                125
Gol H 118:QGTTVTVSSASTK                                                130
```

**B**

**Bre** L 1:DIVLTQSPASLAVSLGQRATISCK-ASQSVDFDGD--SYMNWYQQKPGQPPK**VLIYAASN** 57
Cet L 1:DILLTQSPVILSVSPGERVSFSCR-ASQSIG------TNIHWYQQRTNGSPRLLIKYASE 53
Rit L 1:QIVLSQSPAILSASPGEKVTMTCR-ASSSVS-------YIHWFQQKPGSSPKPWIYATSN 52
Ifx L 1:DILLTQSPAILSVSPGERVSFSCR-ASQFVG------SSIHWYQQRTNGSPRLLIKYASE 53
Atz L 1:DIQMTQSPSSLSASVGDRVTITCR-ASQDVST------AVAWYQQKPGKAPKLLIYSASF 53
Bev L 1:DIQMTQSPSSLSASVGDRVTITCS-ASQDISN------YLNWYQQKPGKAPKVLIYFTSS 53
**Pem** L 1:EIVLTQSPATLSLSPGERATLSCR-ASKGVSTSGY--SYLHWYQQKPGQAPR**LLIYLASY** 57
Tra L 1:DIQMTQSPSSLSASVGDRVTITCR-ASQDVNT------AVAWYQQKPGKAPKLLIYSASF 53
**Ecu** L 1:DIQMTQSPSSLSASVGDRVTITCG-ASENIYG------ALNWYQQKPGKAPK**LLIYGATN** 53
Mep L 1:DIVMTQSPDSLAVSLGERATINCK-SSQSLLNSGNQKNYLAWYQQKPGQPPKLLIYGAST 59
Toc L 1:DIQMTQSPSSLSASVGDRVTITCR-ASQDISS------YLNWYQQKPGKAPKLLIYYTSR 53
**Dur** L 1:EIVLTQSPGTLSLSPGERATLSCR-ASQR**VSSS-----YLAWYQQKPGQAPR**LLIYDASS 54
Ipi L 1:EIVLTQSPGTLSLSPGERATLSCR-ASQSVGSS-----YLAWYQQKPGQAPRLLIYGAFS 54
Niv L 1:EIVLTQSPATLSLSPGEPATLSCR-ASQSVSS------YLAWYQQKPGQAPRLLIYDASN 53
**Ram** L 1:DIQMTQSPSSVSASIGDRVTITCR-ASQGIDN------WLGWYQQKPGKAPKLLIYDASN 53
**Ada** L 1:DIQMTQSPSSLSASVGDRVTITCR-ASQGIRN------YLAWYQQKPGKAPKLLIYAAST 53
**Gol** L 1:EIVLTQSPATLSLSPGERATLSCR-ASQSVYS------YLAWYQQKPGQAPRLLIYDASN 53

**Bre** L 58:**LESGIPAR**FSGSGSGTDFTLNIHPVEEEDAATYYCQQSNEDP-WTFGGGTKLEIK 112
Cet L 54:SISGIPSRFSGSGSGTDFTLSINSVESEDIADYYCQQNNNWP-TTFGAGTKLELK 108
Rit L 53:LASGVPVRFSGSGSGTSYSLTISRVEAEDAATYYCQQWTSNP-PTFGGGTKLEIK 107
Ifx L 54:SMSGIPSRFSGSGSGTDFTLSINTVESEDIADYYCQQSHSWP-FTFGSGTNLEVK 108
Atz L 54:LYSGVPSRFSGSGSGTDFTLTISSLQPEDFATYYCQQYLYHP-ATFGQGTKVEIK 108
Bev L 54:LHSGVPSRFSGSGSGTDFTLTISSLQPEDFATYYCQQYSTVP-WTFGQGTKVEIK 108
**Pem** L 58:**LESGVPAR**FSGSGSGTDFTLTISSLEPEDFAVYYCQHSRDLP-LTFGGGTKVEIK 112
Tra L 54:LYSGVPSRFSGSRSGTDFTLTISSLQPEDFATYYCQQHYTTP-PTFGQGTKVEIK 108
**Ecu** L 54:**LADGVPSR**FSGSGSGTDFTLTISSLQPEDFATYYCQNVLNTP-LTFGQGTKVEIK 108
Mep L 60:RESGVPDRFSGSGSGTDFTLTISSLQAEDVAVYYCQNVHSFP-FTFGGGTKLEIK 114
Toc L 54:LHSGVPSRFSGSGSGTDFTFTISSLQPEDIATYYCQQGNTLP-YTFGQGTKVEIK 108
**Dur** L 55:RATGIPDRFSGSGSGTDFTLTISRLEPEDFAVYYCQQYGSLP-WTFGQGTKVEIK 109
Ipi L 55:RATGIPDRFSGSGSGTDFTLTISRLEPEDFAVYYCQQYGSSP-WTFGQGTKVEIK 109
Niv L 54:RATGIPARFSGSGSGTDFTLTISSLEPEDFAVYYCQQSSNWP-RTFGQGTKVEIK 108
**Ram** L 54:LDTGVPSRFSGSGSGTYFTLTISSLQAEDFAVYFCQQAK**AFP-PTFGGGTK**VDIK 108
**Ada** L 54:LQSGVPSRFSGSGSGTDFTLTISSLQPEDVATYYCQRYNR**AP-YTFGQGTK**VEIK 108
**Gol** L 54:RATGIPARFSGSGSGTDFTLTISSLEPEDFAVYYCQQR**SNWPPFTFGPGTK**VDIK 109

**C**

Ave L 1:QSALTQPASVSGSPGQSITISCTGTSSDVGGYNYVSWYQQHPGKAPKLMIYDVSNRPSGV 60
Ave L 61:SNRFSGSKSGNTASLTISGLQAEDEADYYCSSYTSSSTRVFGTGTKVTVL 110

**D**

Abt 1:**MHVAQPAVVLASSR**GIASFVCEYASPGKATEVRVTVLRQADSQVTEVCAATYMMGNELTF 60
Abt 61:LDDSICTGTSSGNQVNLTIQGLRAMDTGLYICKVELMYPPPYYLGIGNGTQIYVIDPEPC 120
Abt 121:PDSDQEPK 128

**E**

Etn 1:LPAQVAFTPYAPEPGSTCRLREYYDQTAQMCCSKCSPGQHAK**VFCTK**TSDTVCDSCEDST 60
Etn 61:YTQLWNWVPECLSCGSRCSSDQVETQACTREQNRICTCRPGWYCALSKQEGCRLCAPLRK 120
Etn 121:CRPGFGVARPGTETSDVVCKPCAPGTFSNTTSSTDICRPHQICNVVAIPGNASMDAVCTS 180
Etn 181:TSPTRSMAPGAVHLPQPVSTRSQHTQPTPEPSTAPSTSFLLPMGPSPPAEGSTGDEPK 238

**Figure S2. Location of signature peptides.** Sequence alignment of each IgG-formed mAbs Fv region on A) heavy chain, B) light kappa chain, C) light lambda chain, and fused domain of Fc-fusion proteins D) abatacept, and E) etanercept. The sequence of signature peptide is highlighted in gray. mAbs that have signature peptides on heavy chain are shown in red, and those on light chain are shown in blue. Bre: brentuximab vedotin, Cet: cetuximab, Rit: rituximab, Ifx: infliximab, Atz: atezolizumab, Bev: bevacizumab, Pem: pembrolizumab, Tra: trastuzumab, Ecu: eculizumab, Mep: mepolizumab, Toc: tocilizumab, Ave: avelumab, Dur: durvalumab, Ipi: ipilimumab, Niv: nivolumab, Ram: ramucirumab, Ada: adalimumab, Gol: golimumab, Abt: abatacept, Etn: etanercept.

**A)** Bre, VLIYAASNLESGIPAR (precursor m/z 837.5009)

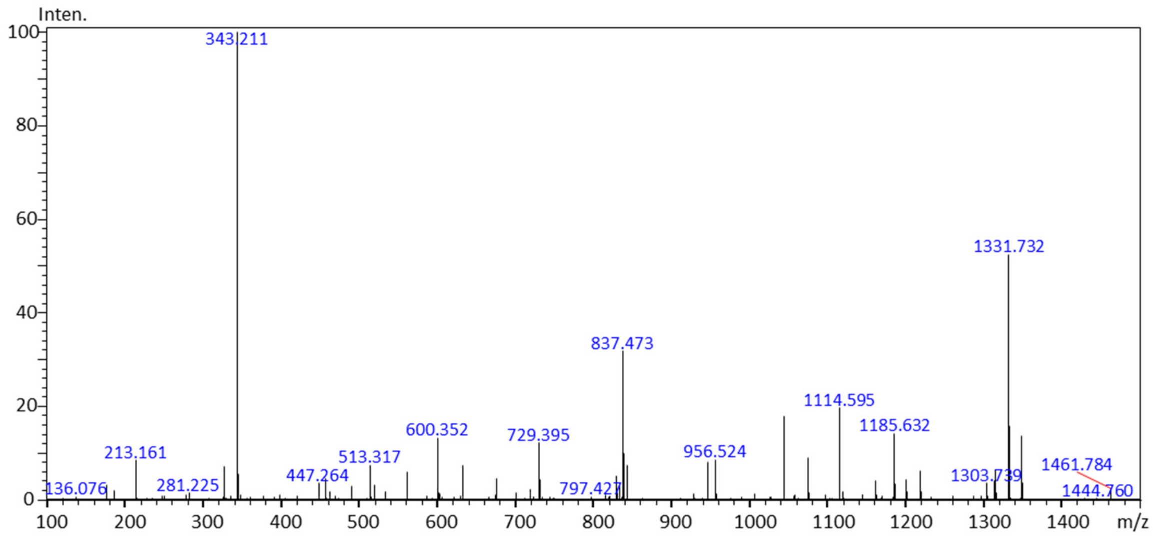


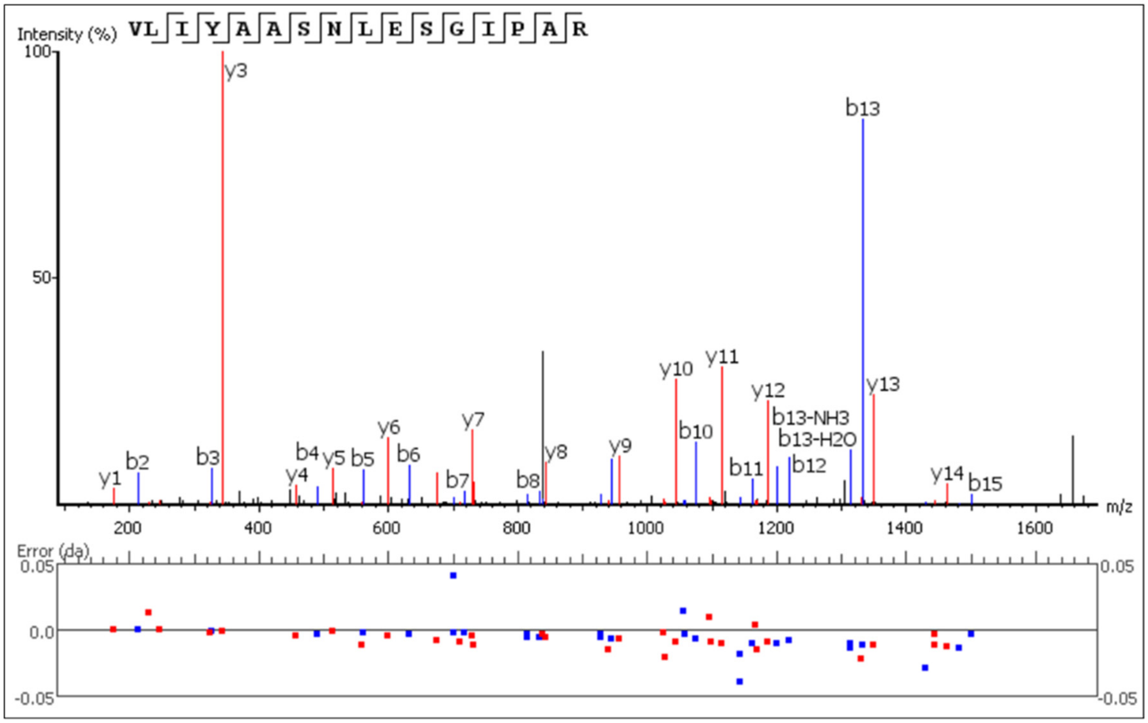

**B)** Cet, SQVFFK (prec. m/z 378.2106)

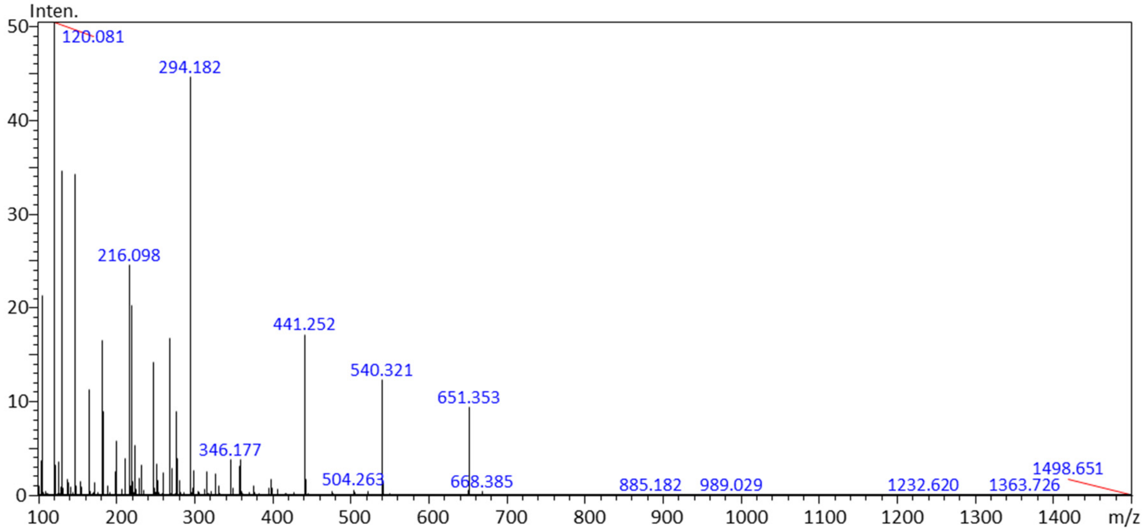


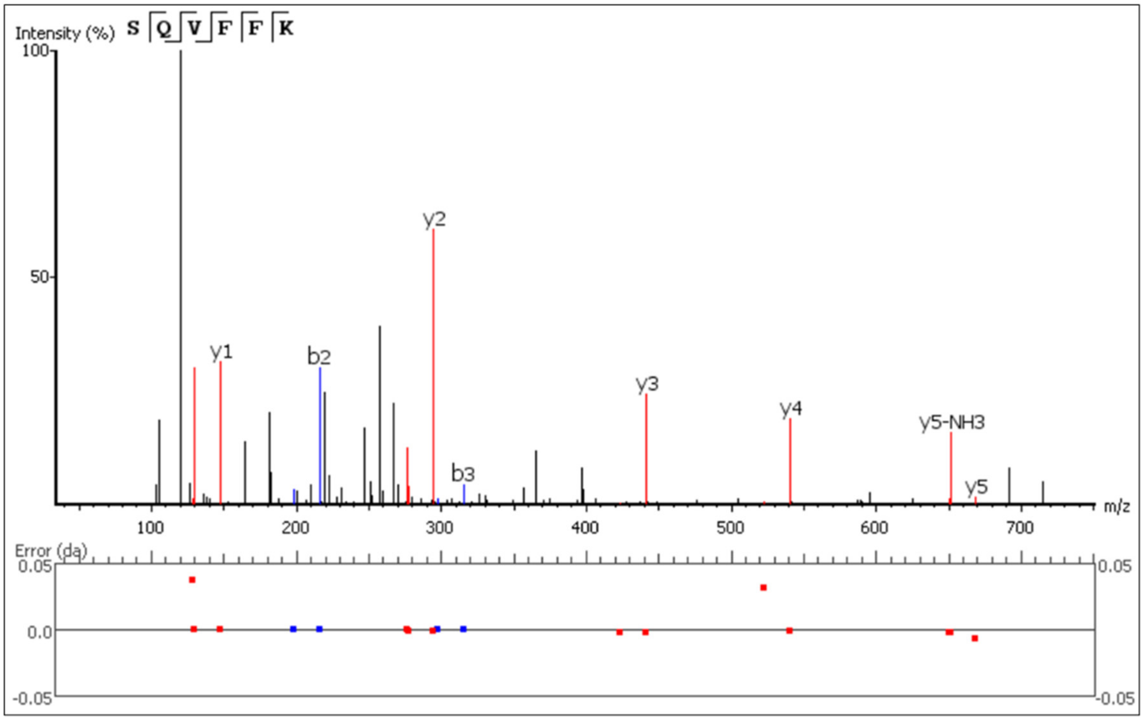

**C)** Rit, ASGYTFTSYNMHWVK (prec. m/z 597.9327)

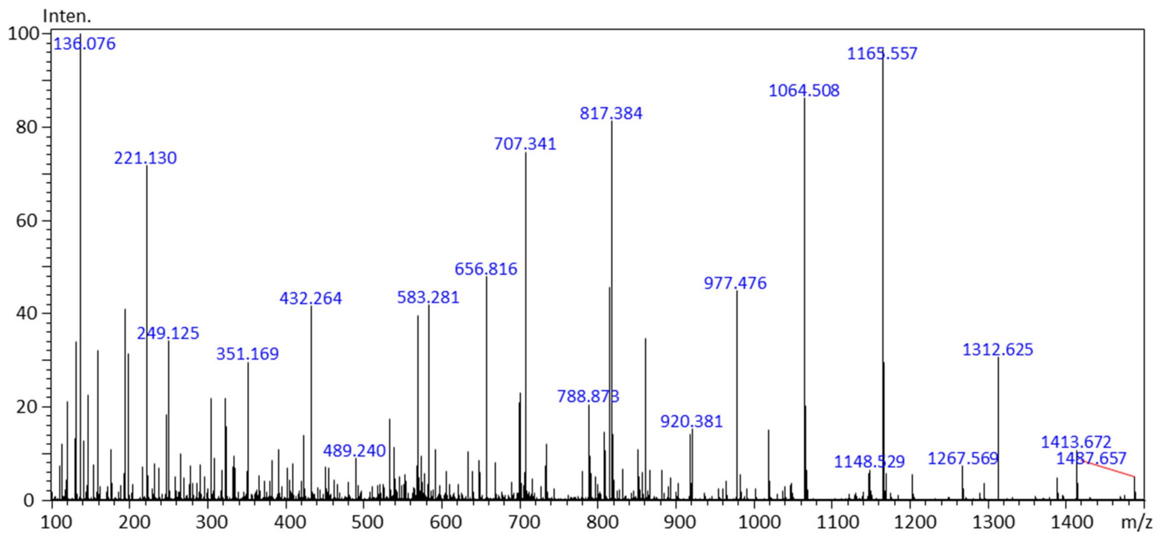


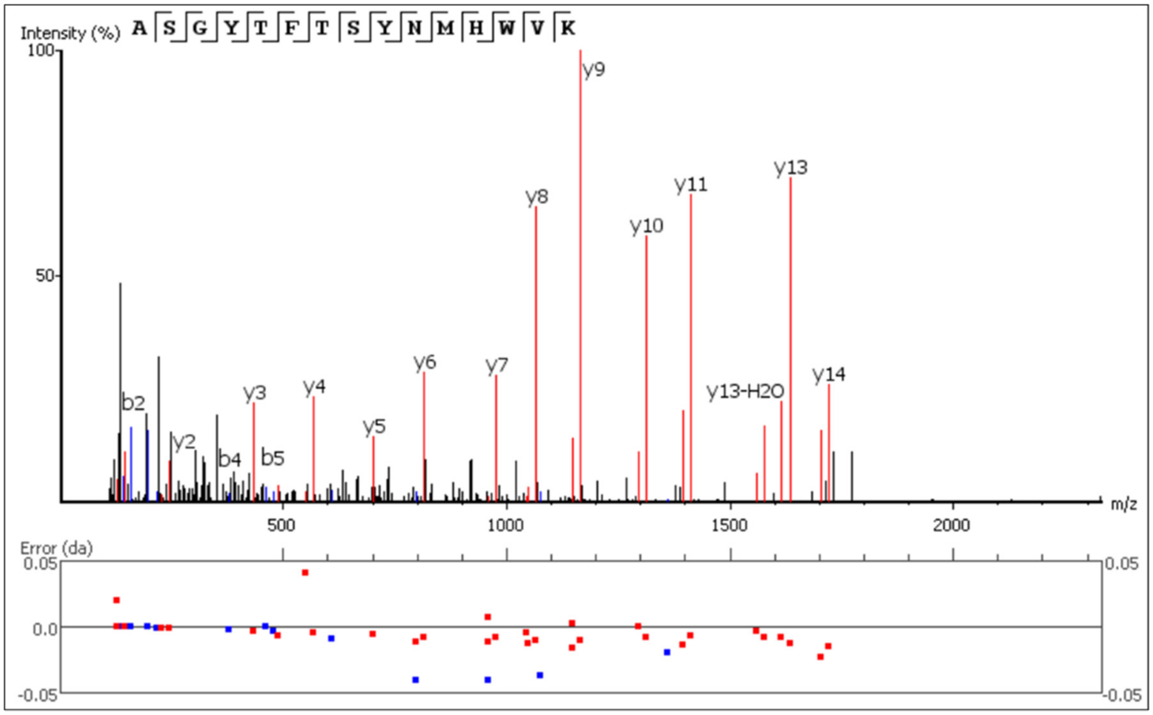

**D)** Ifx, SINSATHYAESVK (prec. m/z 703.8737)

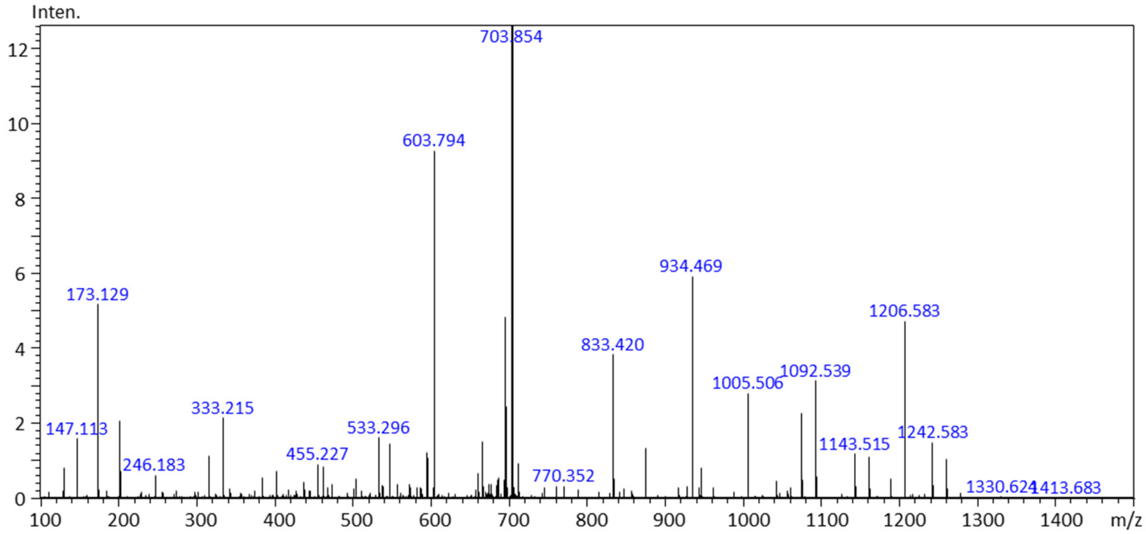


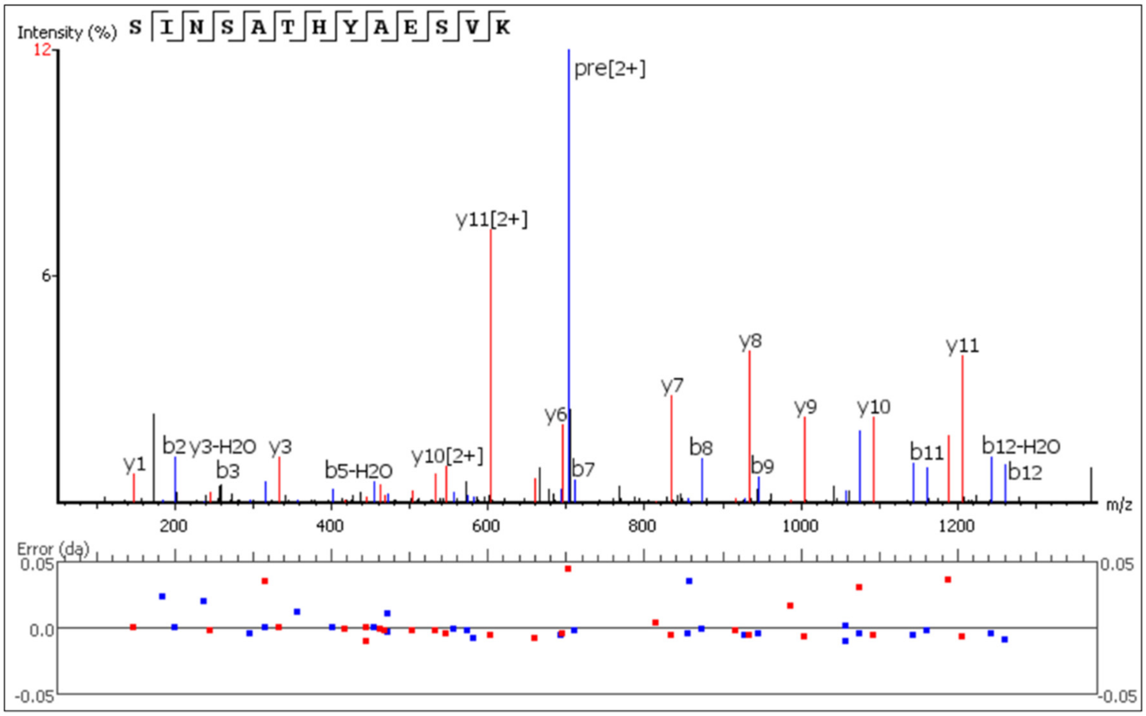

**E)** Atz, RHWPGGFDYWGQGTLVTVSSASTK (prec. m/z 660.1810)

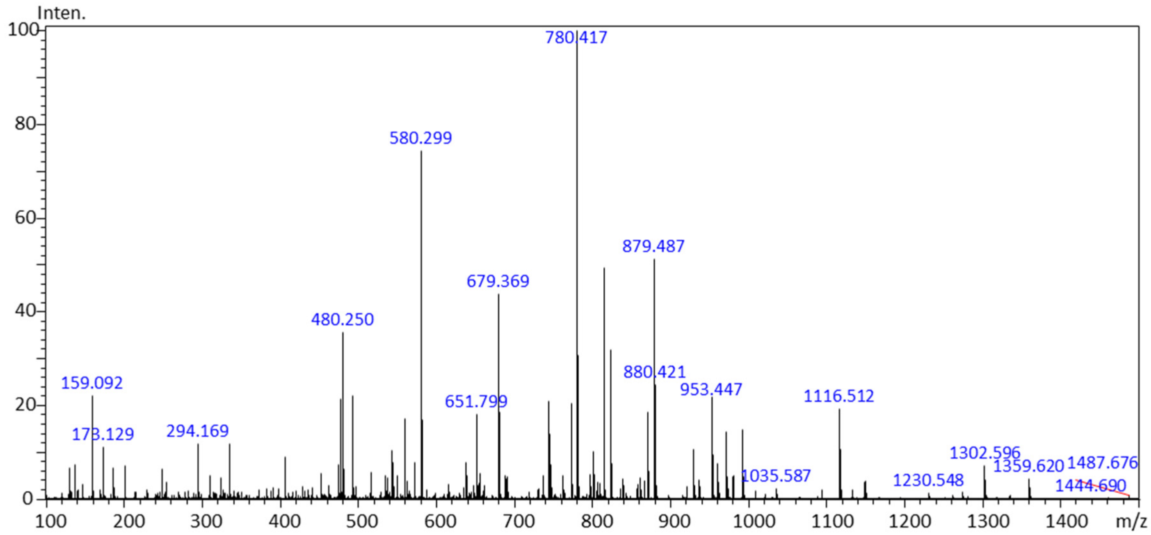


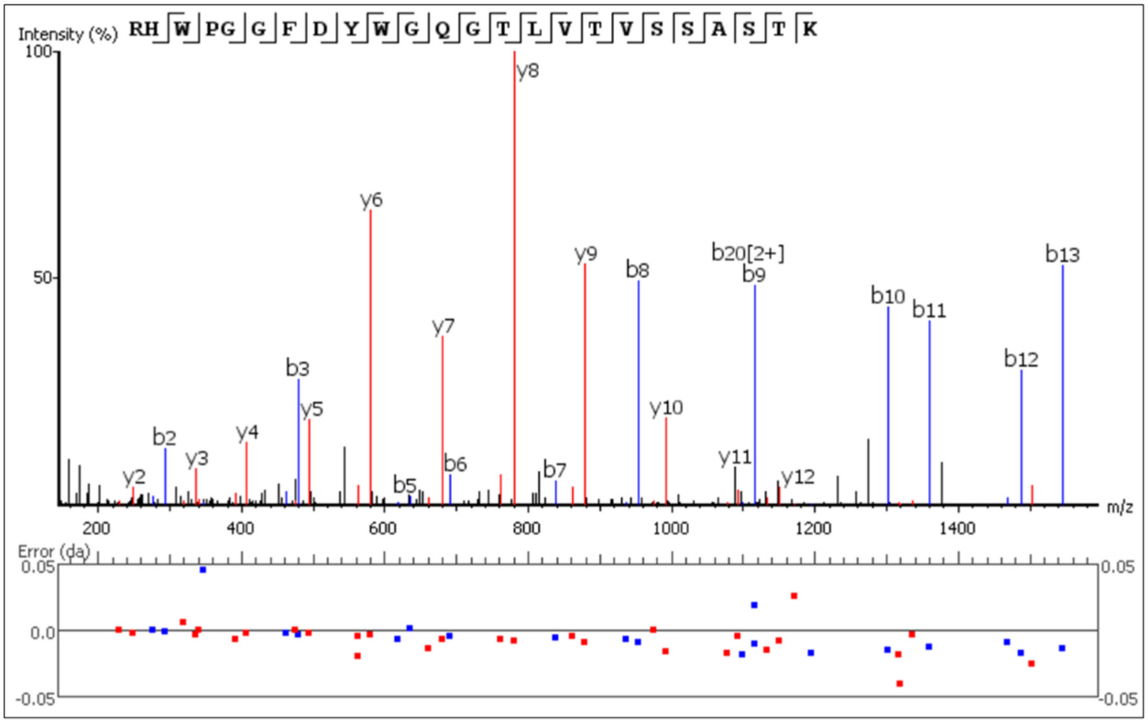

**F)** Bev, FTFSLDTSK (prec. m/z 523.2654)

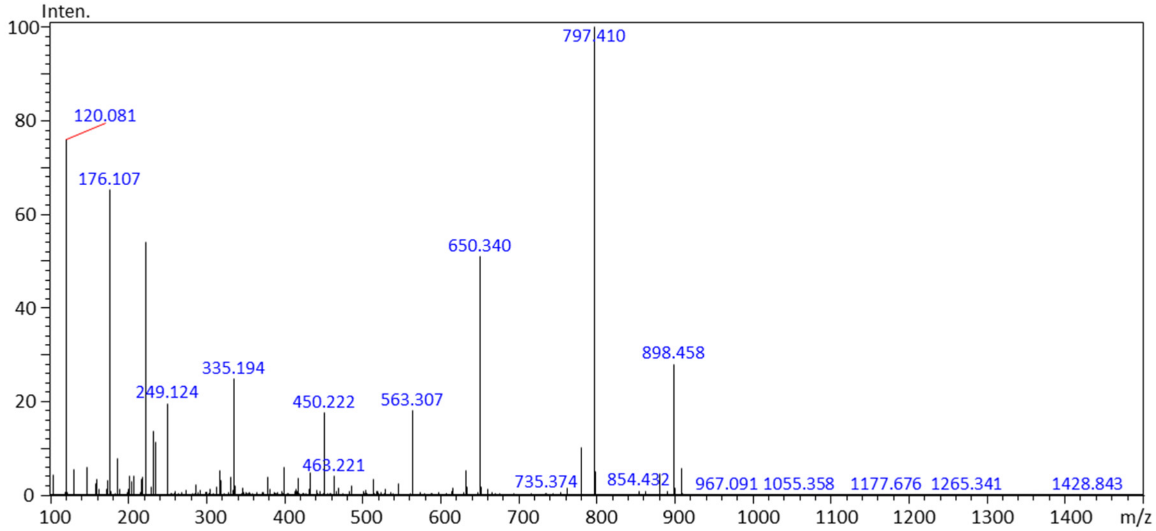


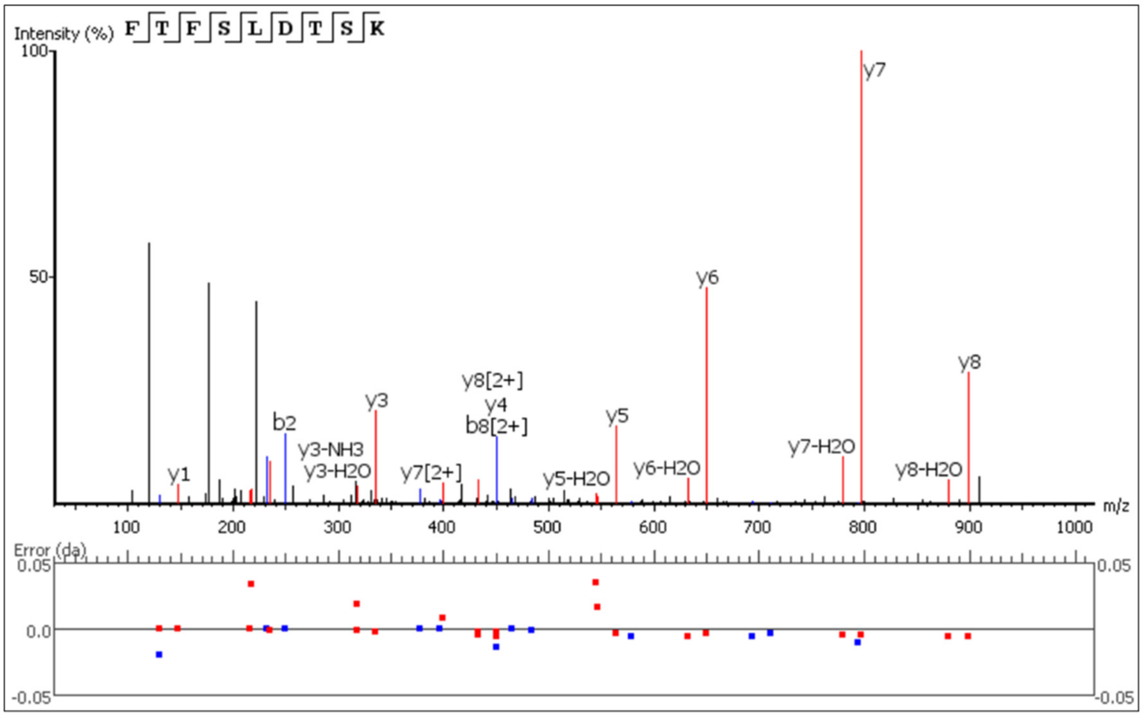

**G)** Pem, LLIYLASYLESGVPAR (prec. m/z 883.0929)

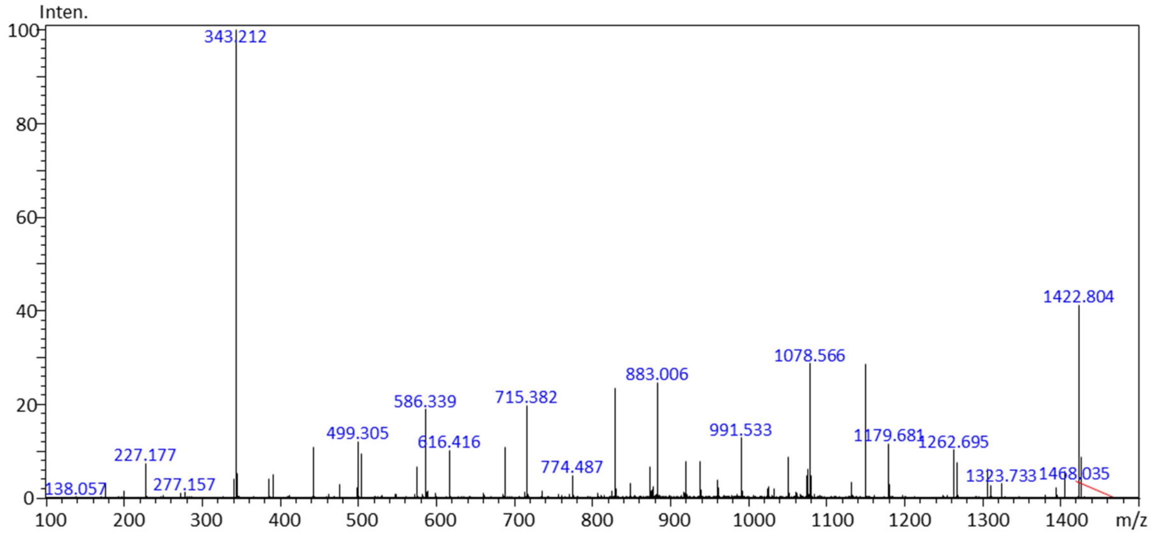


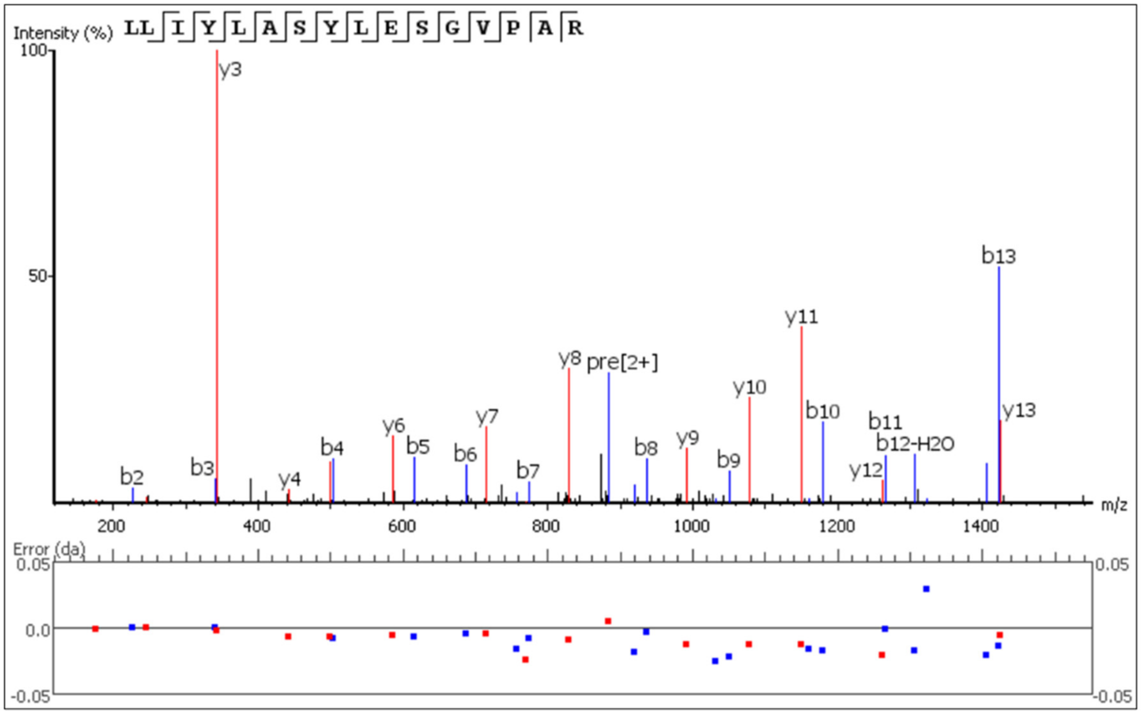

**H)** Tra, IYPTNGYTR (prec. m/z 542.7740)

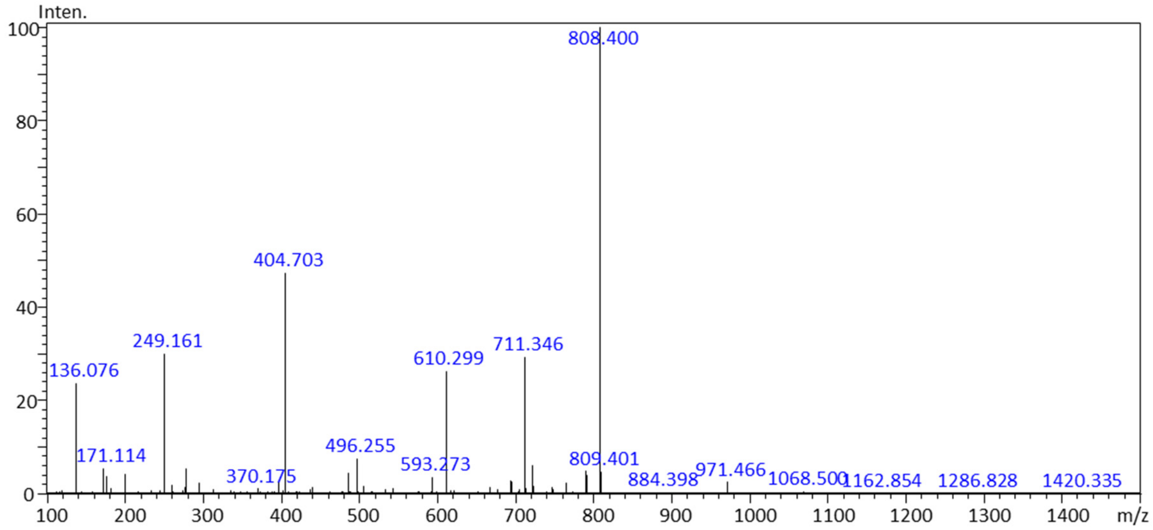


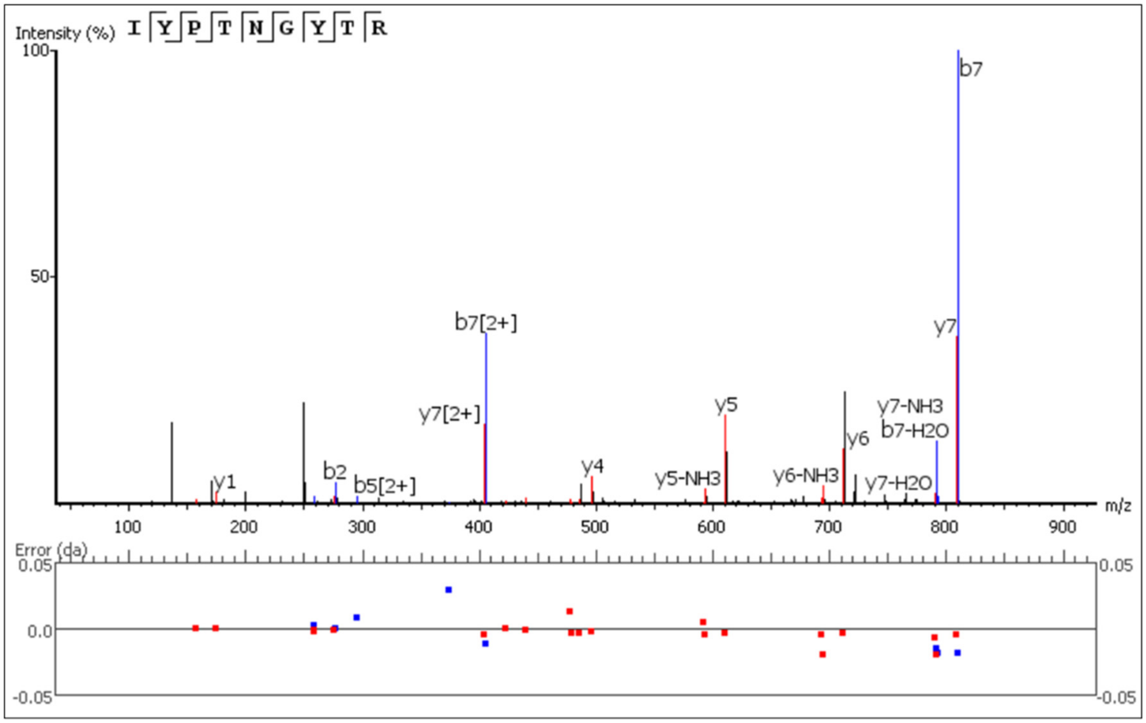

**I)** Ecu, LLIYGATNLADGVPSR (prec. m/z 830.4581)

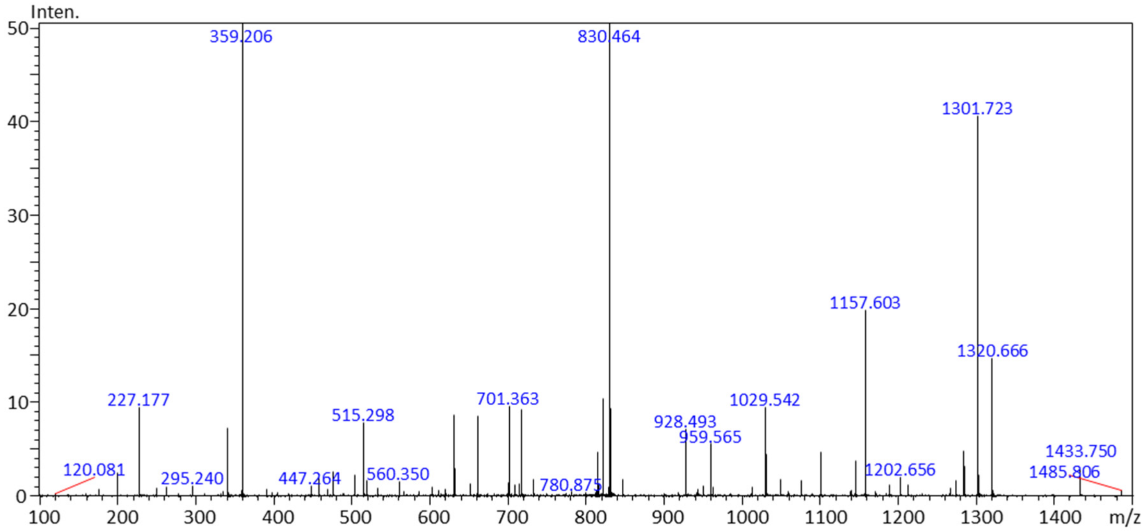


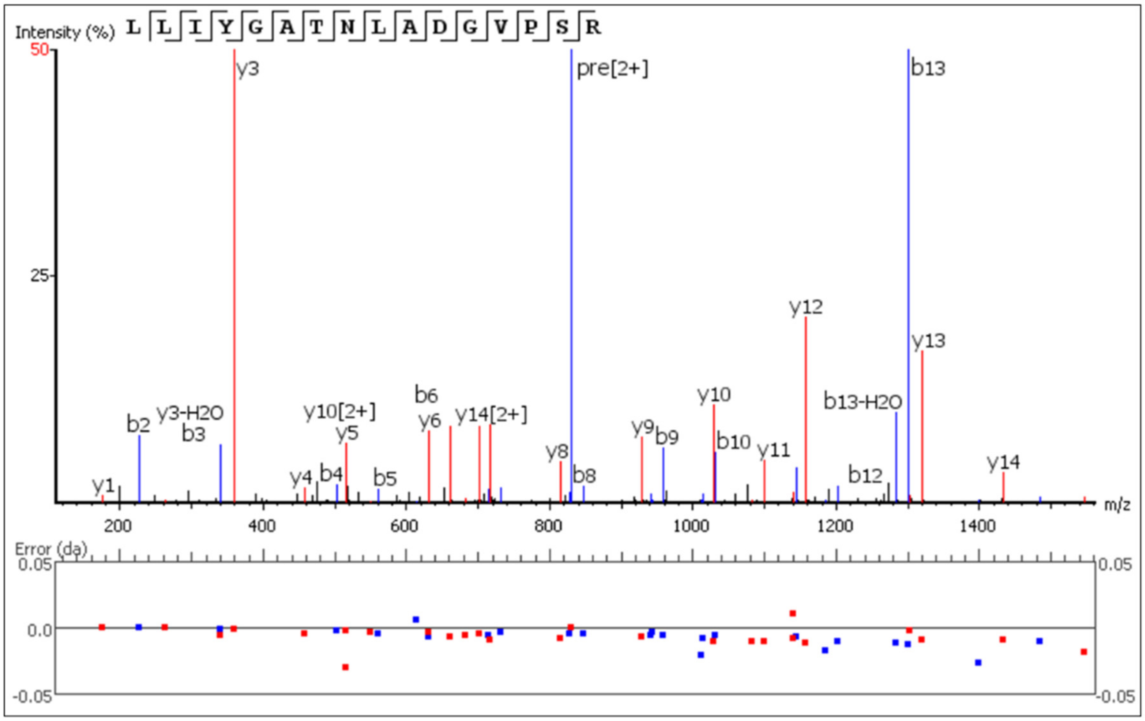

**J)** Mep, DPPSSLLR (prec. m/z 442.7524)

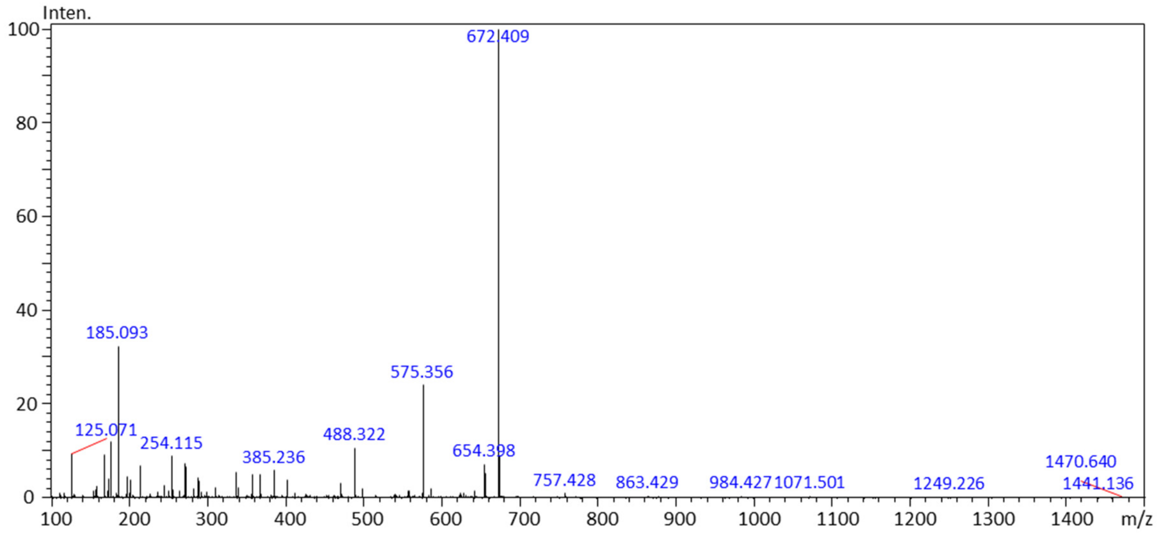


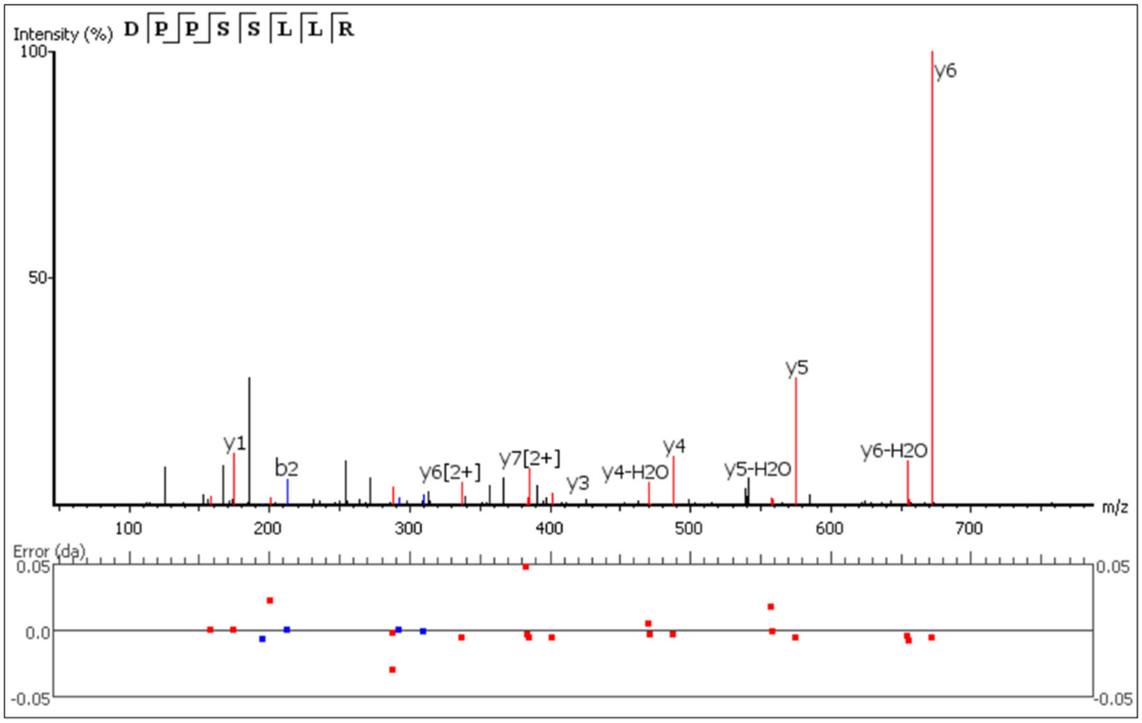

**K)** Toc, VTMLR (prec. m/z 310.1837)

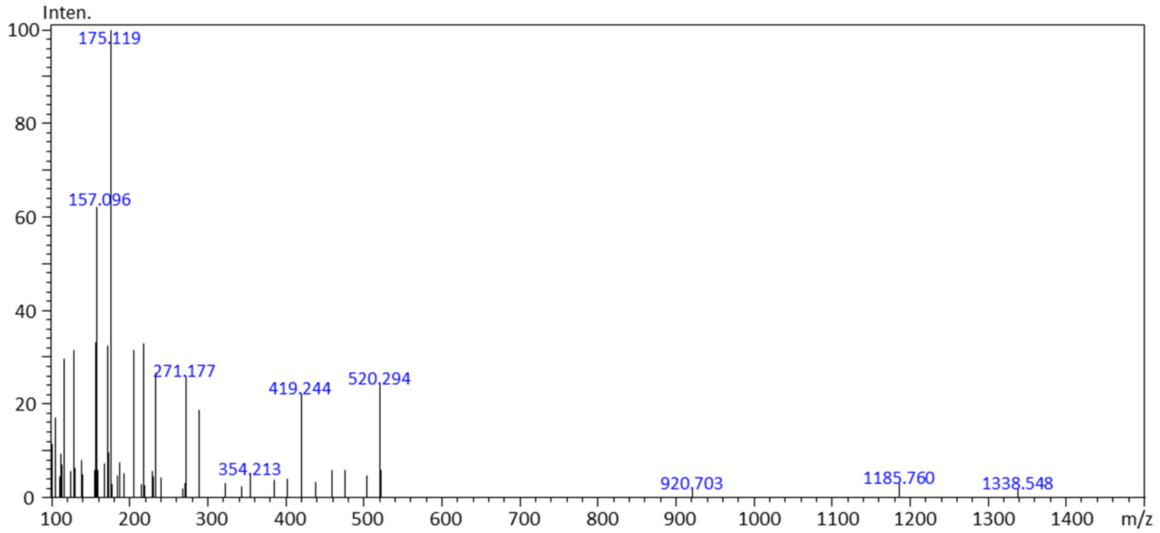


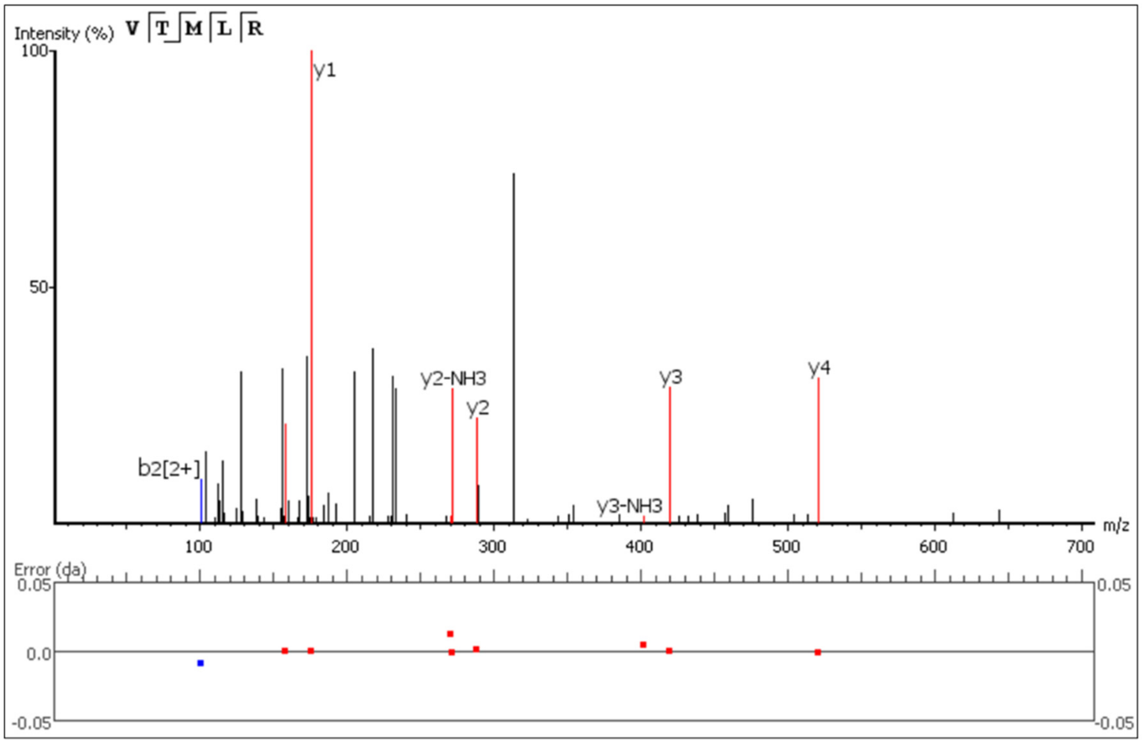

**L)** Ave, LGTVTTVDYWGQGTLVTVSSASTK (prec. m/z 824.7394)

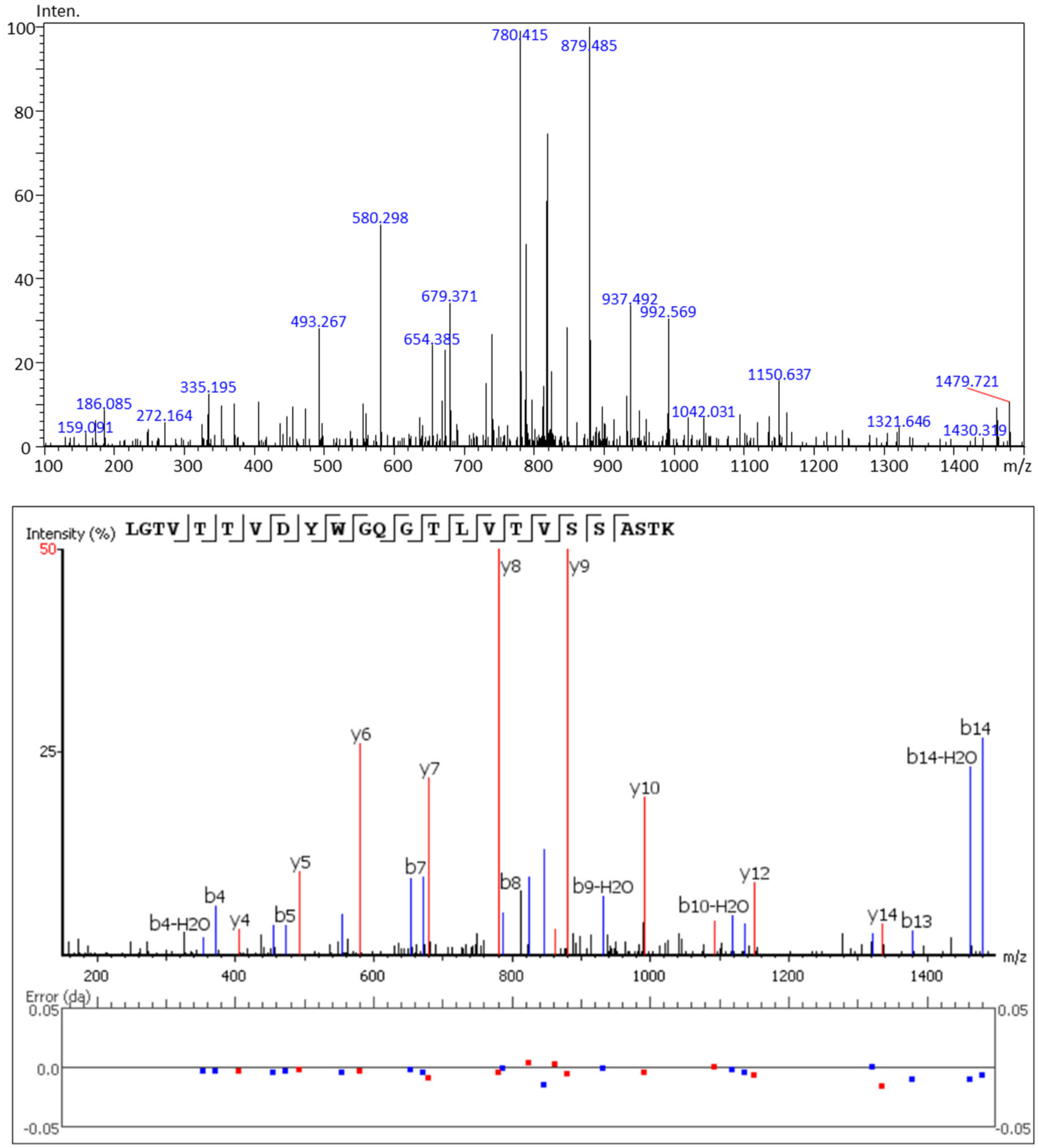

**M)** Dur, VSSSYLAWYQQKPGQAPR (prec. m/z 689.3567)

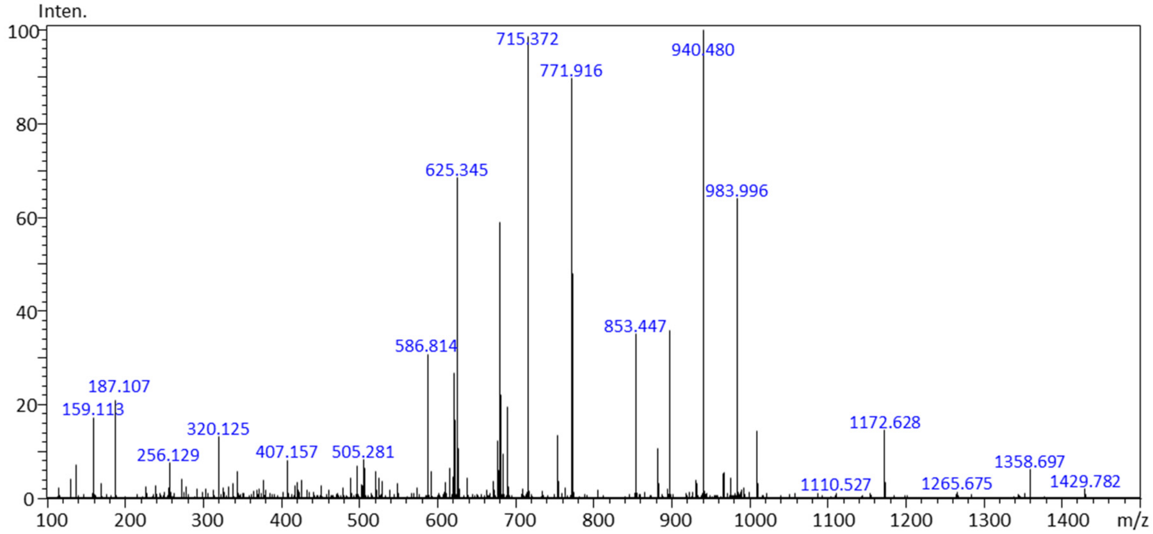


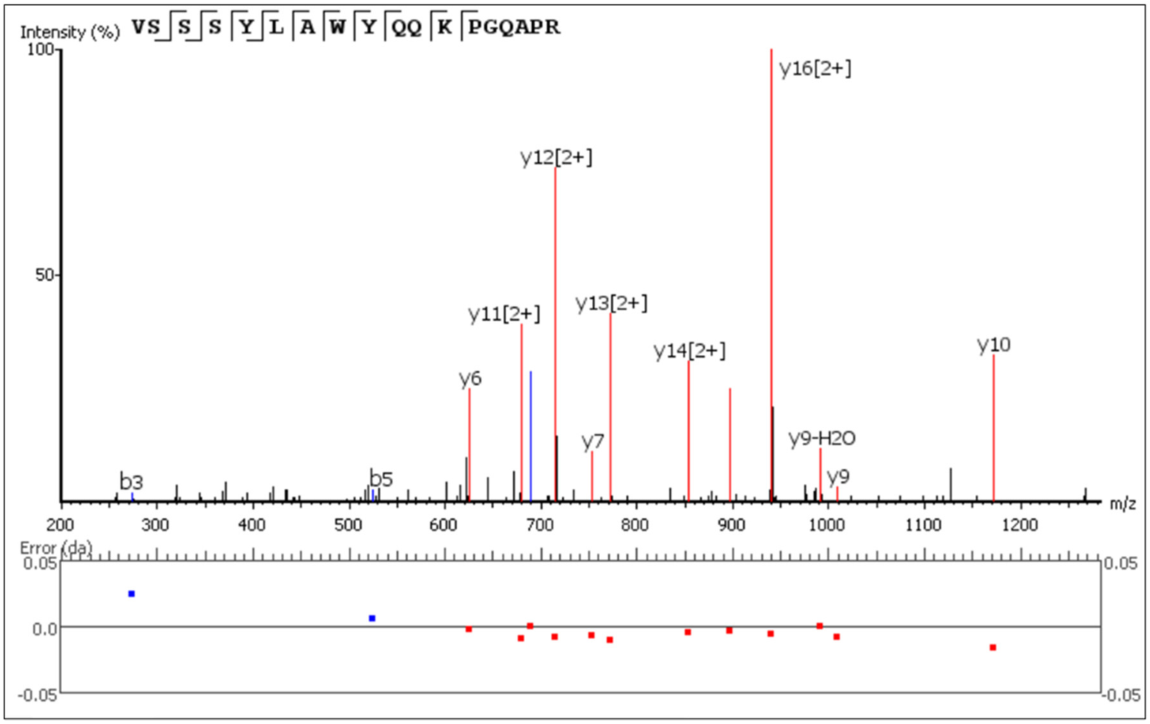

**N)** Ipi, TGWLGPFDYWGQGTLVTVSSASTK (prec. m/z 853.5245)

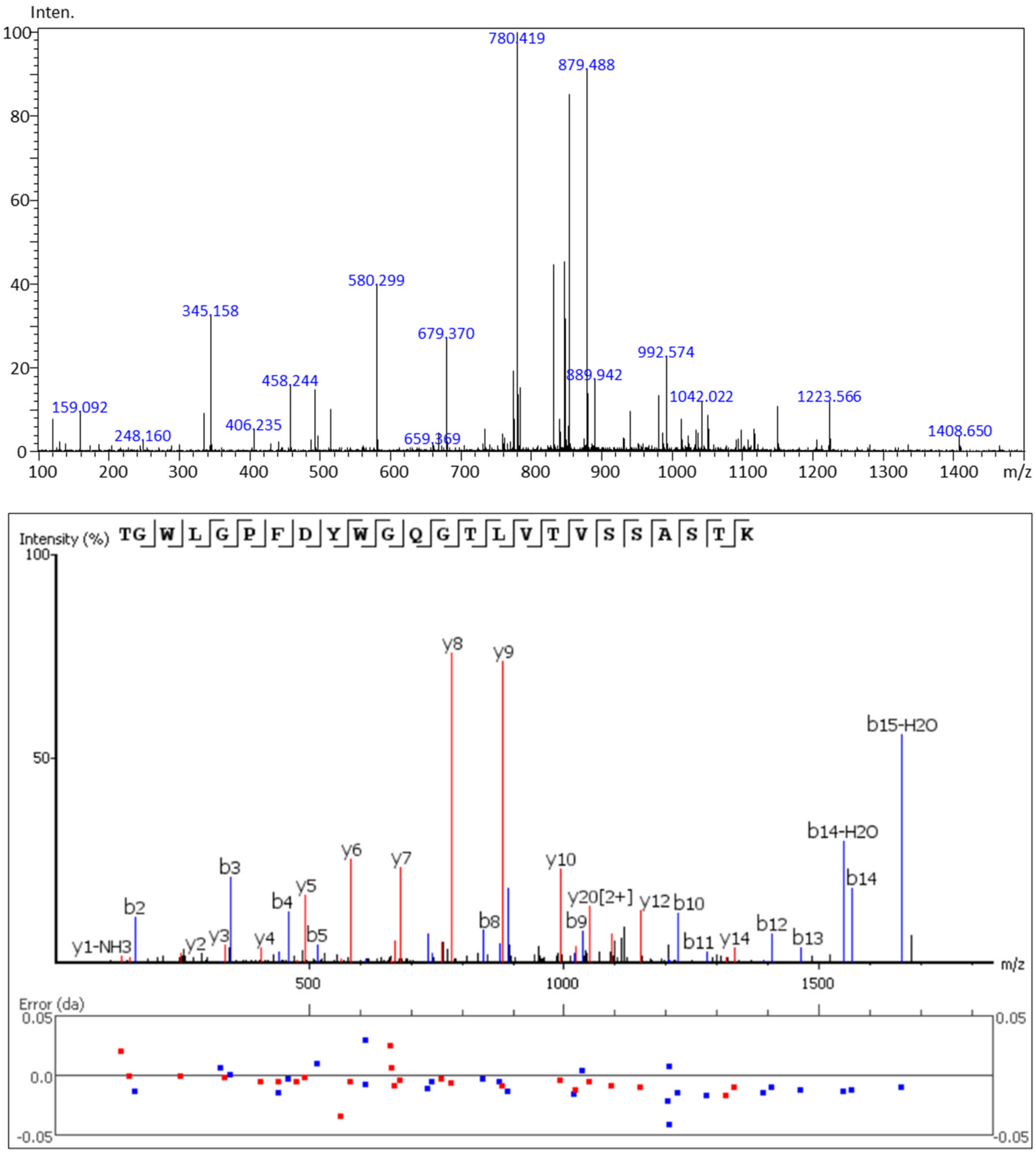

**O)** Niv, ASGITFSNSGMHWVR (prec. m/z 550.5323)

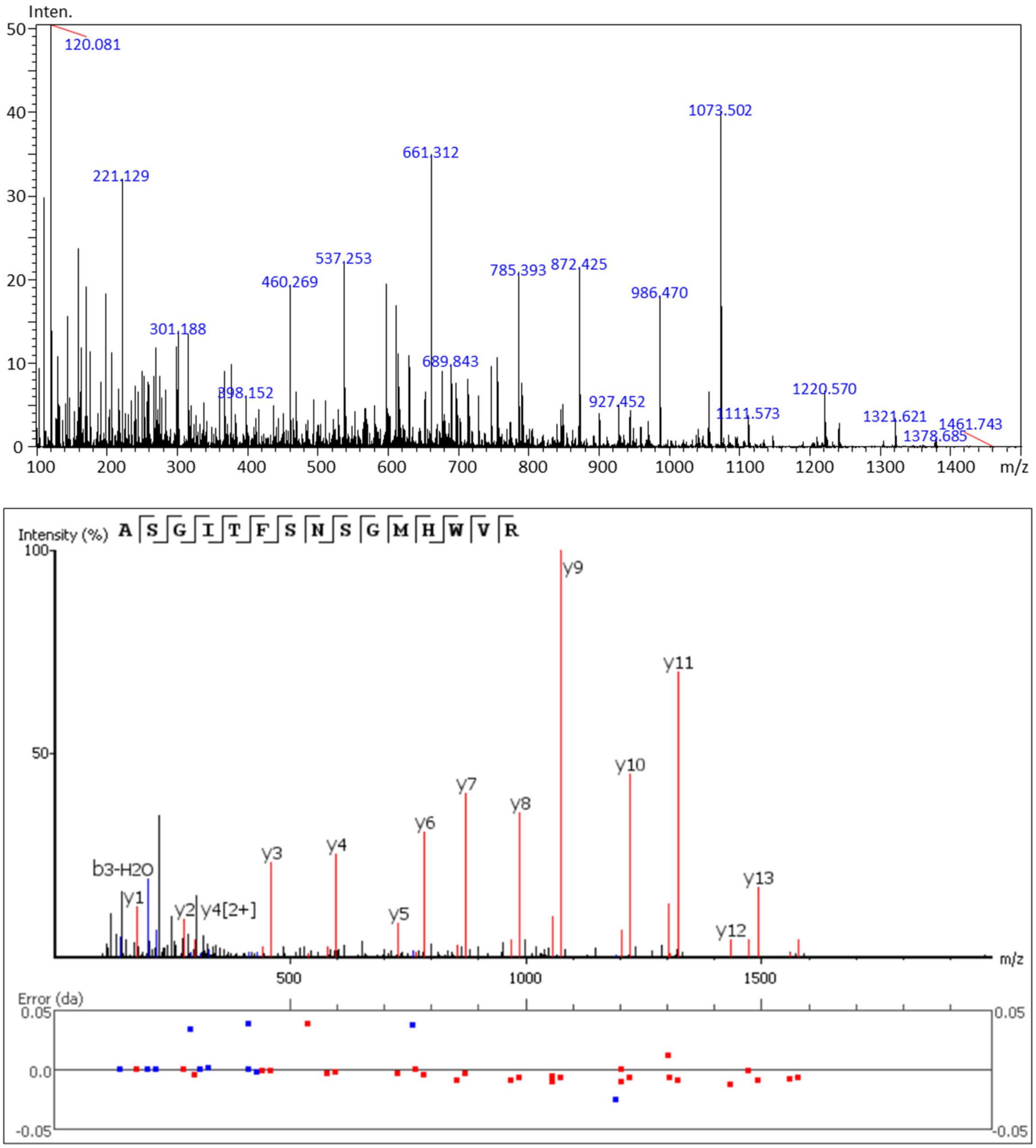

**P)** Ram, AFPPTFGGGTK (prec. m/z 540.2824)

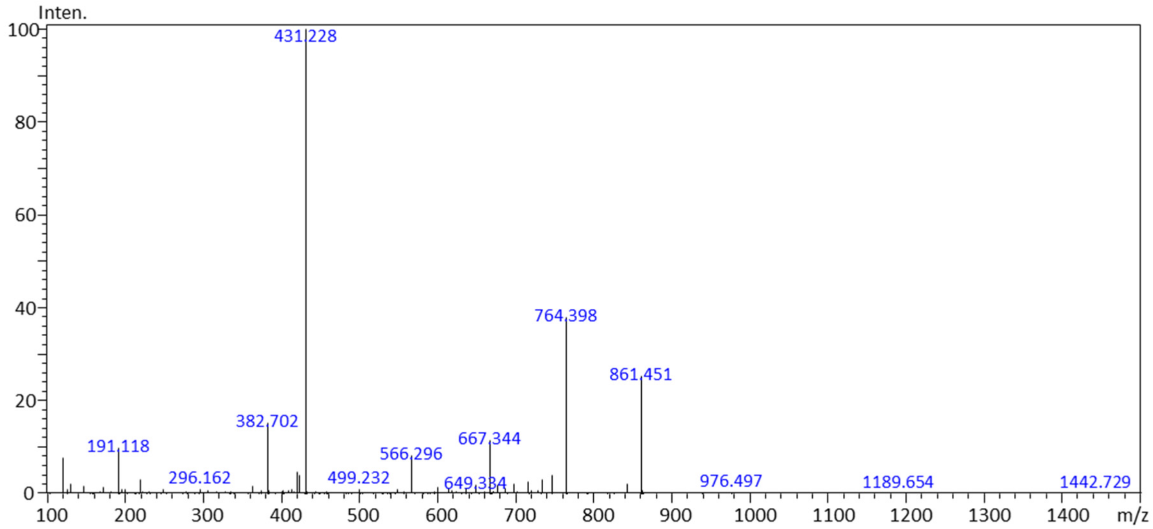


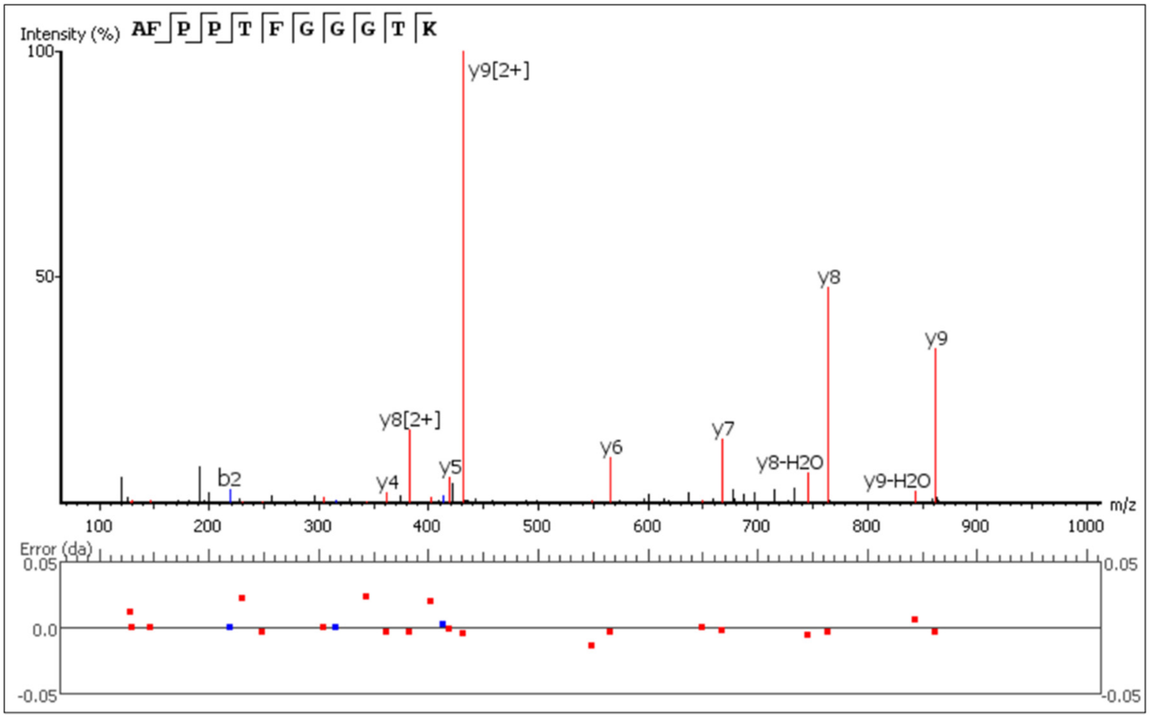

**Q)** Ada, APYTFGQGTK (prec. m/z 535.2791)

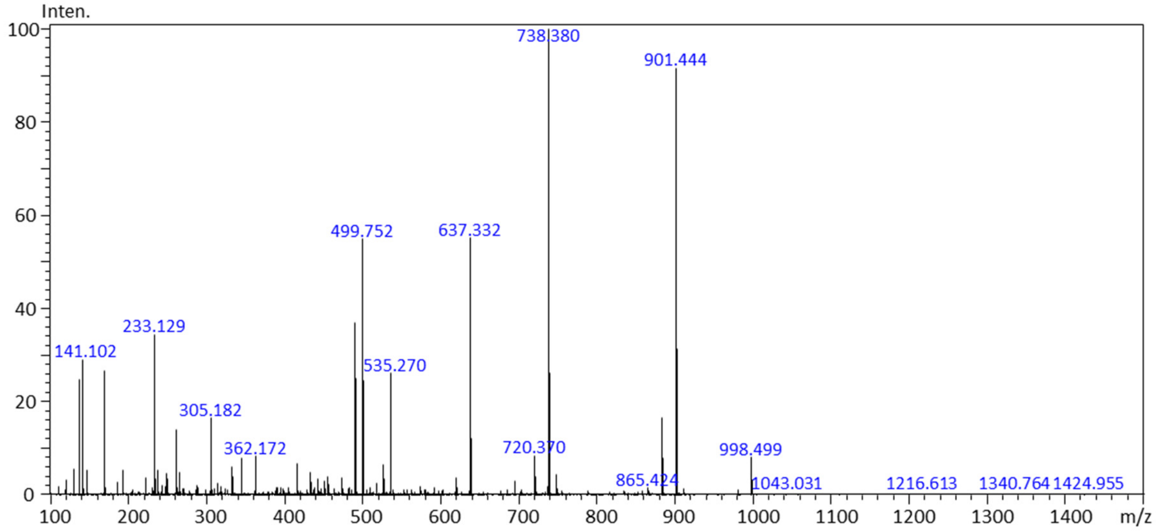


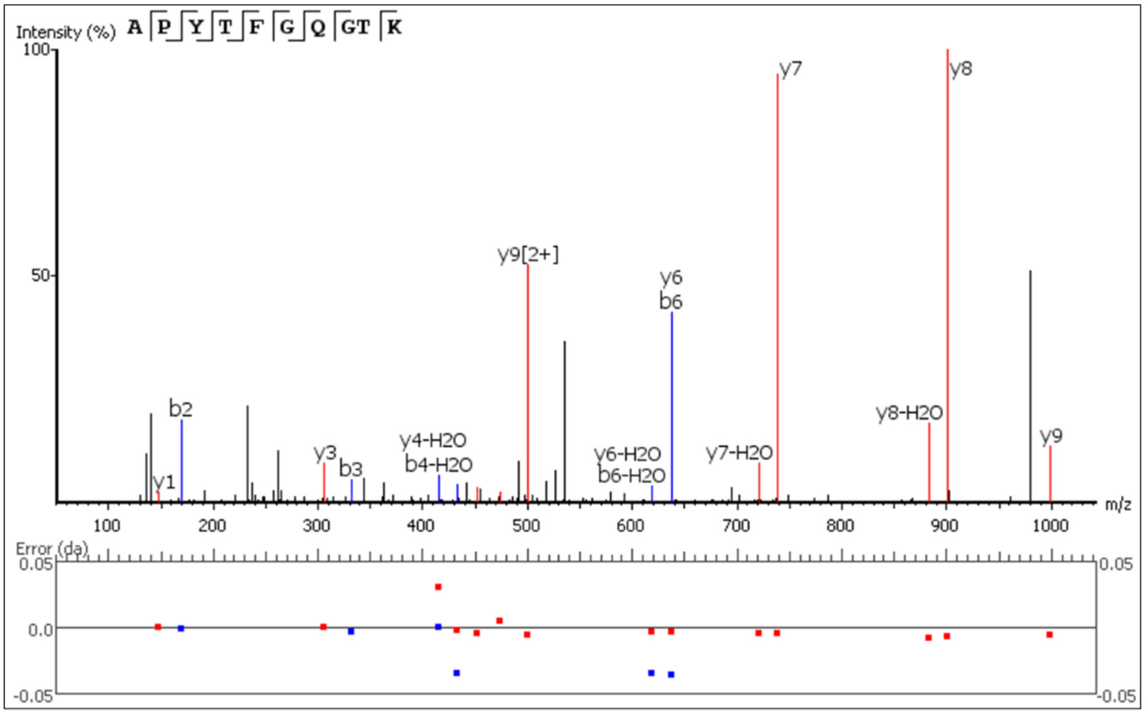

**R)** Gol, SNWPPFTFGPGTK (prec. m/z 718.3620)

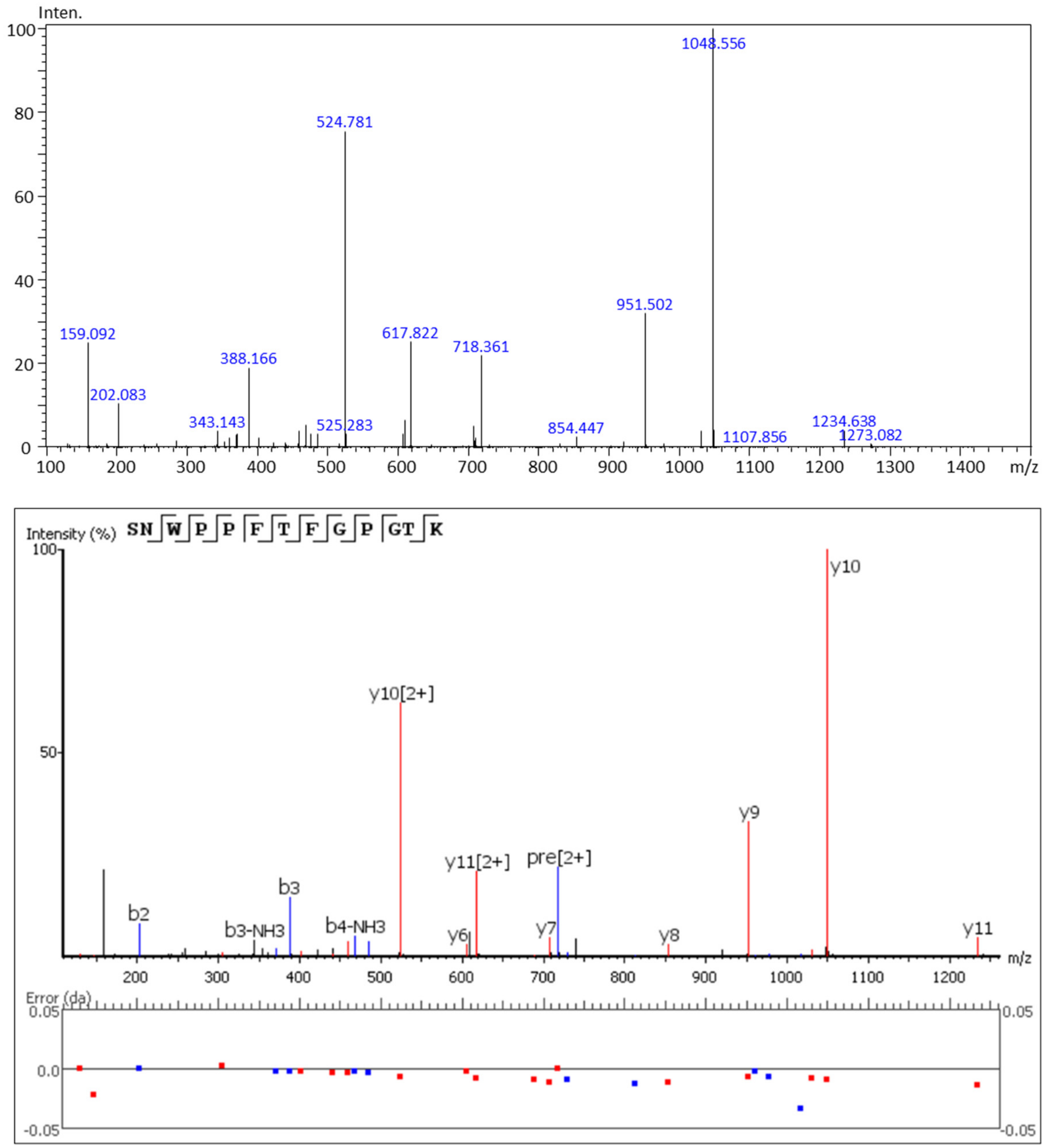

**Figure S3. MS/MS spectra and sequence assignment of each signature peptide on 18-plex refmAb-Q nSMOL analysis using Q-TOF-MS.** Top box shows the MS/MS spectra, and middle box is the assignment of ion series of y- (red) and b- (blue) ions. Bottom box shows the error distribution of fragment ions (Da). A) Bre, VLIYAASNLESGIPAR (precursor m/z 837.5009), B) Cet, SQVFFK (prec. m/z 378.2106), C) Rit, ASGYTFTSYNMHWVK (prec. m/z 597.9327), D) Ifx, SINSATHYAESVK (prec. m/z 703.8737), E) Atz, RHWPGGFDYWGQGTLVTVSSASTK (prec. m/z 660.1810), F) Bev, FTFSLDTSK (prec. m/z 523.2654), G) Pem, LLIYLASYLESGVPAR (prec. m/z 883.0929), H) Tra, IYPTNGYTR (prec. m/z 542.7740), I) Ecu, LLIYGATNLADGVPSR (prec. m/z 830.4581), J) Mep, DPPSSLLR (prec. m/z 442.7524), K) Toc, VTMLR (prec. m/z 310.1837), L) Ave, LGTVTTVDYWGQGTLVTVSSASTK (prec. m/z 824.7394), M) Dur, VSSSYLAWYQQKPGQAPR (prec. m/z 689.3567), N) Ipi, TGWLGPFDYWGQGTLVTVSSASTK (prec. m/z 853.5245), O) Niv, ASGITFSNSGMHWVR (prec. m/z 550.5323), P) Ram, AFPPTFGGGTK (prec. m/z 540.2824), Q) Ada, APYTFGQGTK (prec. m/z 535.2791), R) Gol, SNWPPFTFGPGTK (prec. m/z 718.3620).

**A)** The ratio to brenzuximab

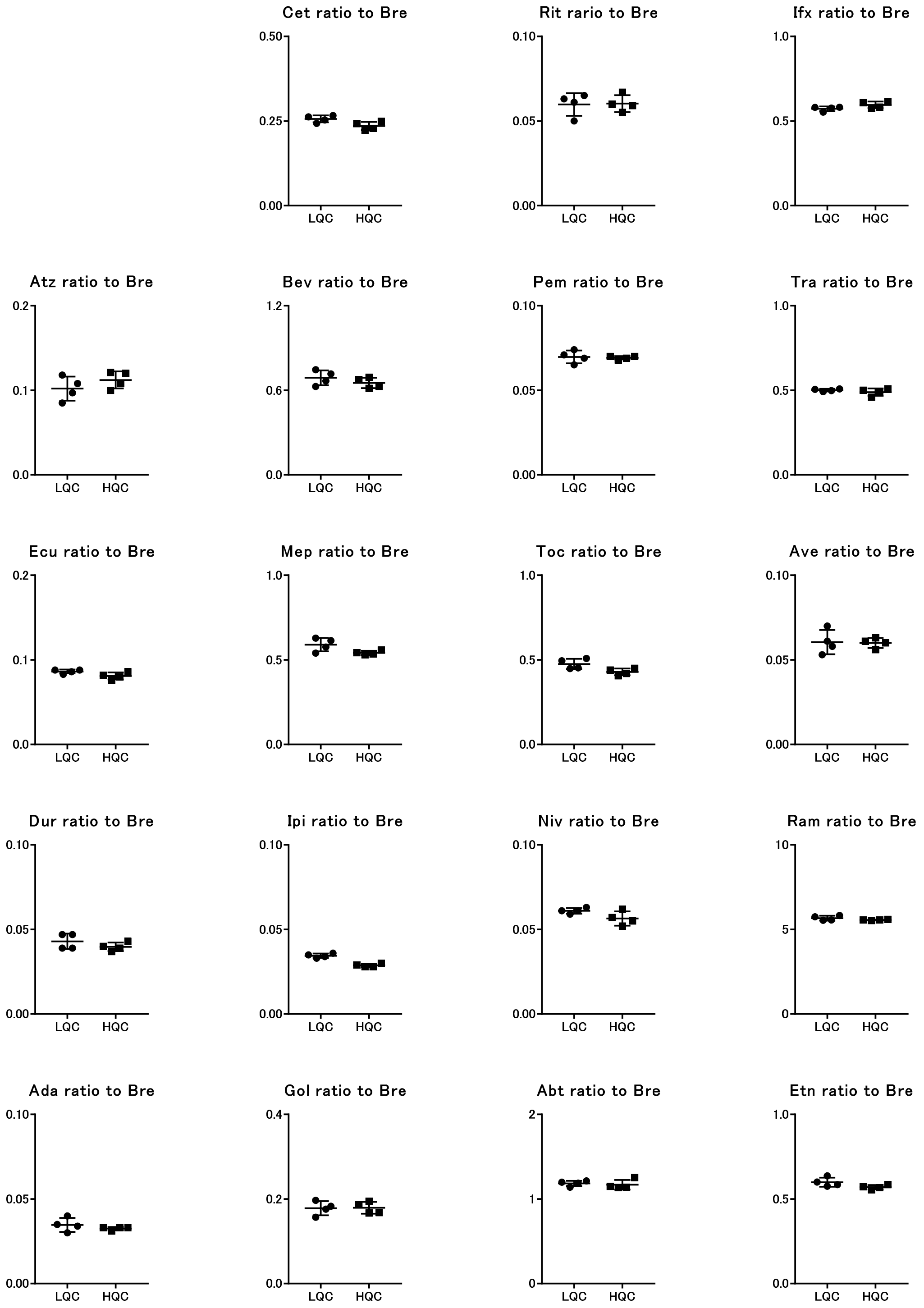

**B)** The ratio to cetuximab

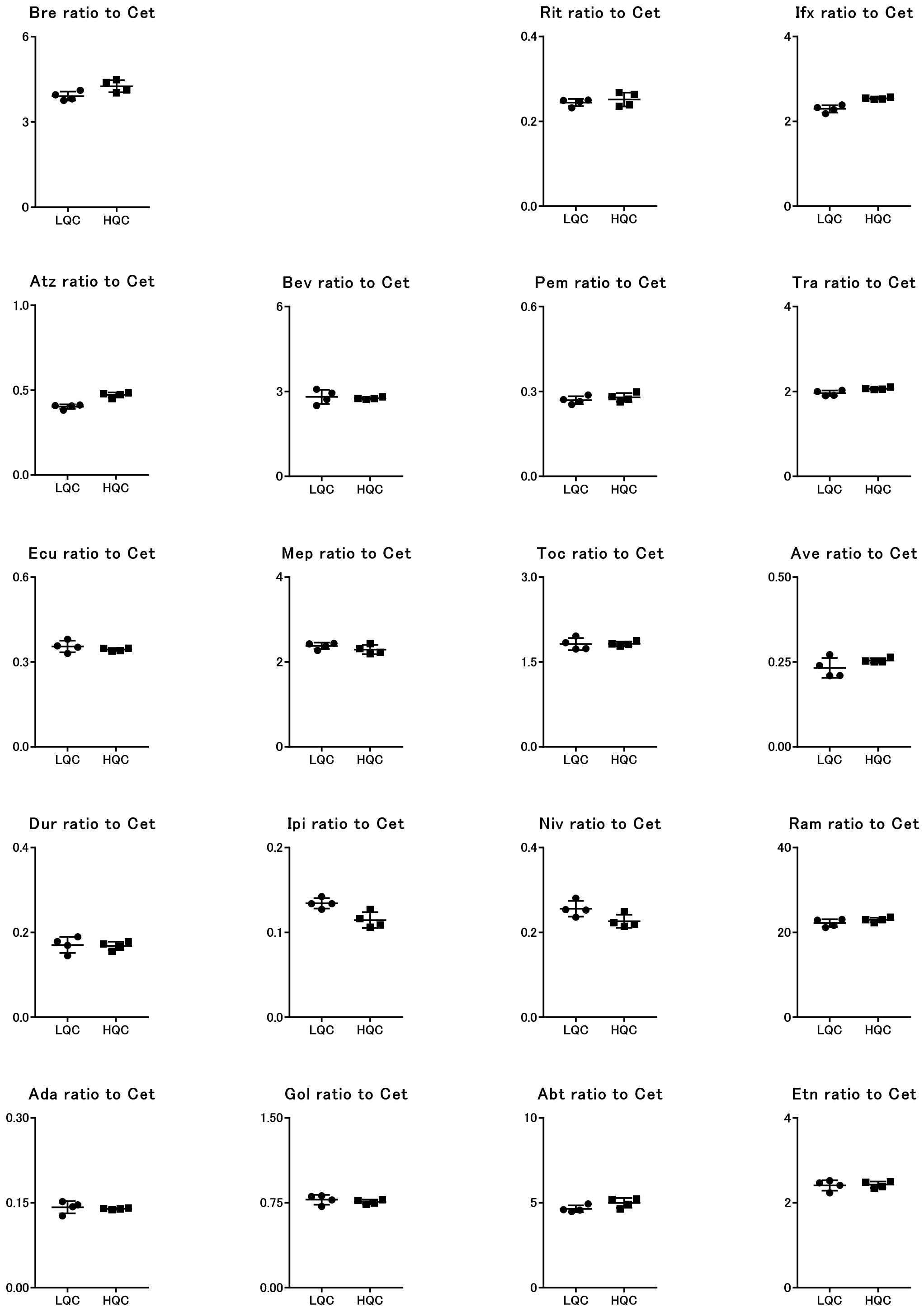

**C)** The ratio to rituximab

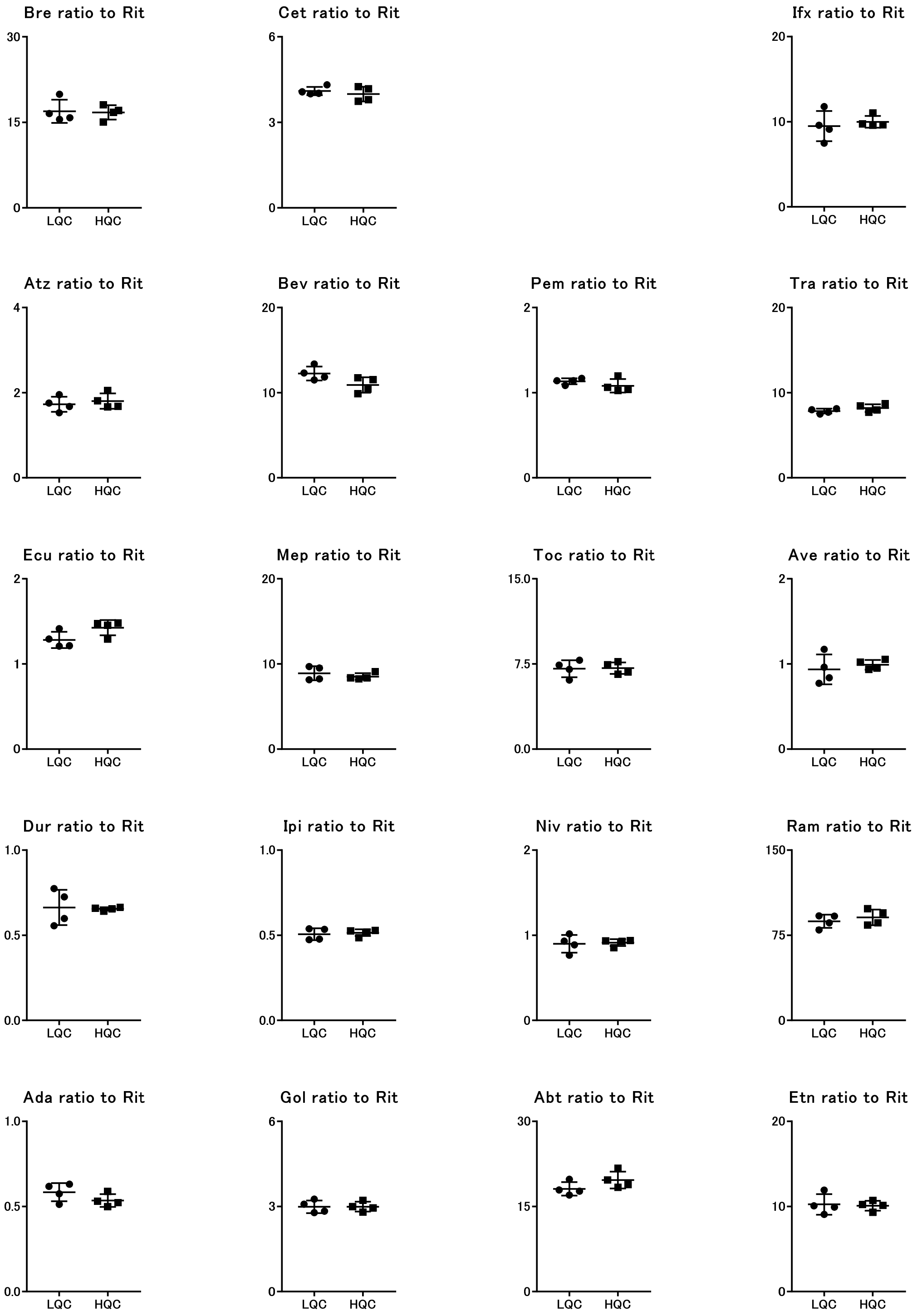

**D)** The ratio to infliximab

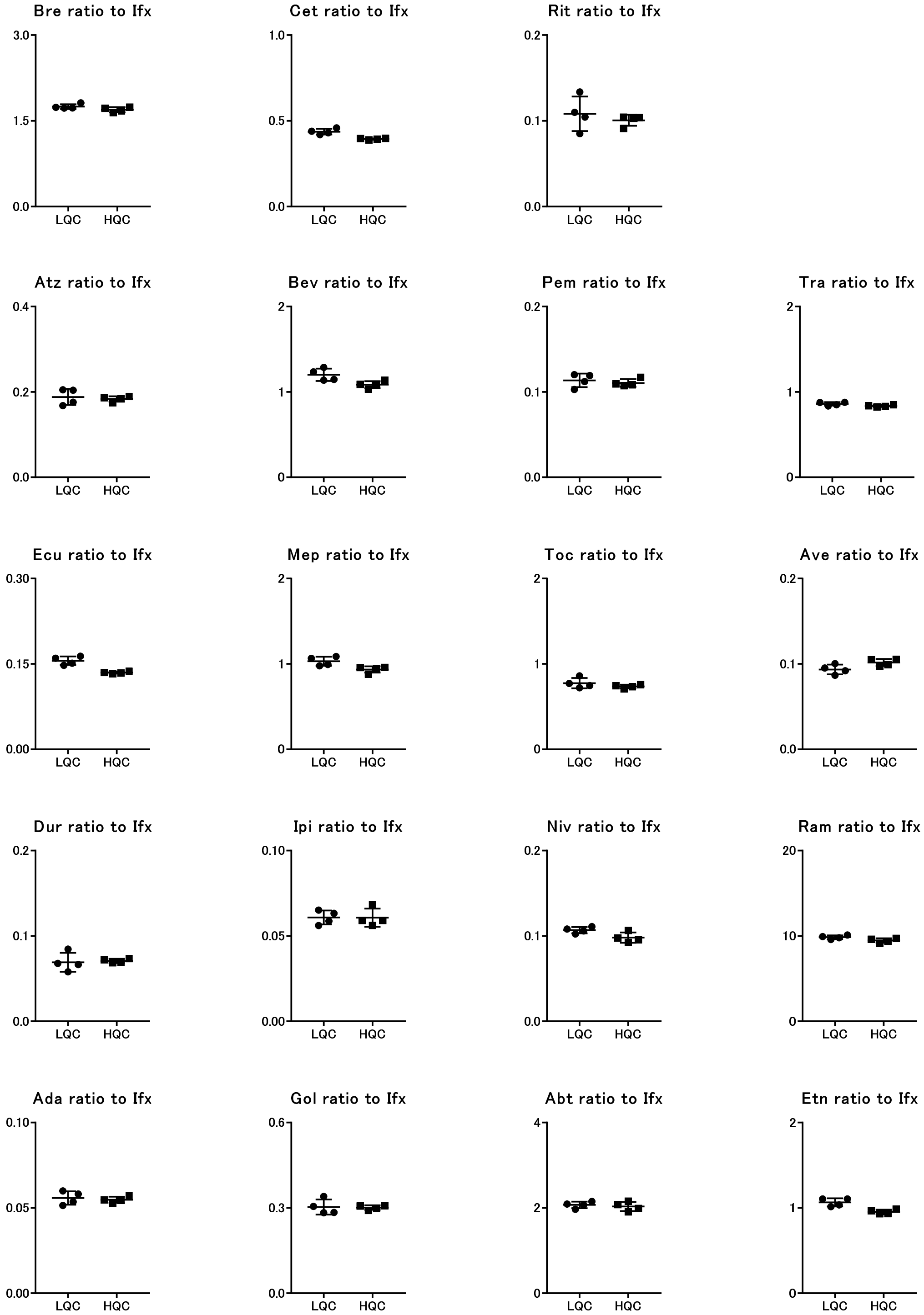

**E)** The ratio to atezolizumab

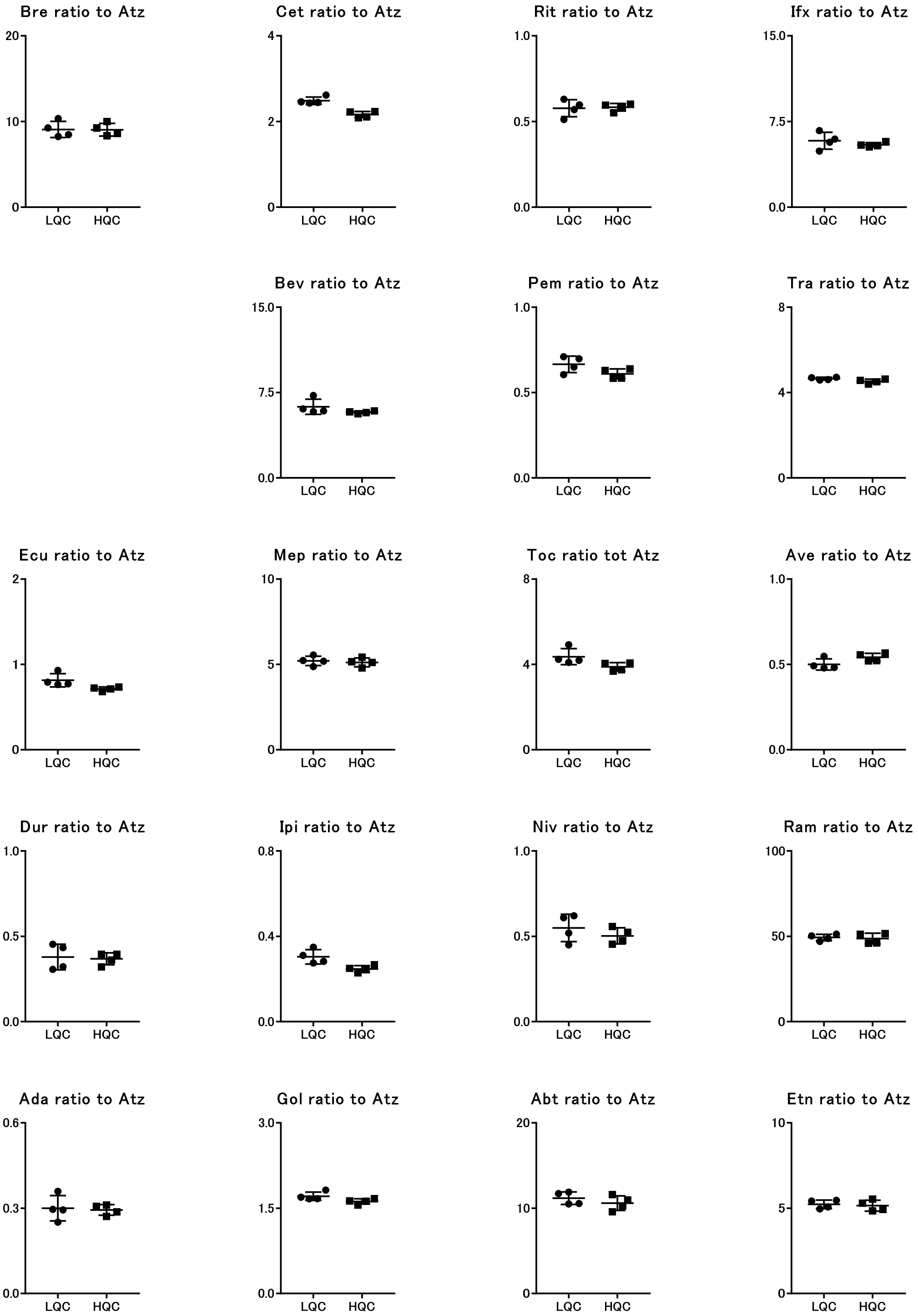

**F)** The ratio to bevacizumab

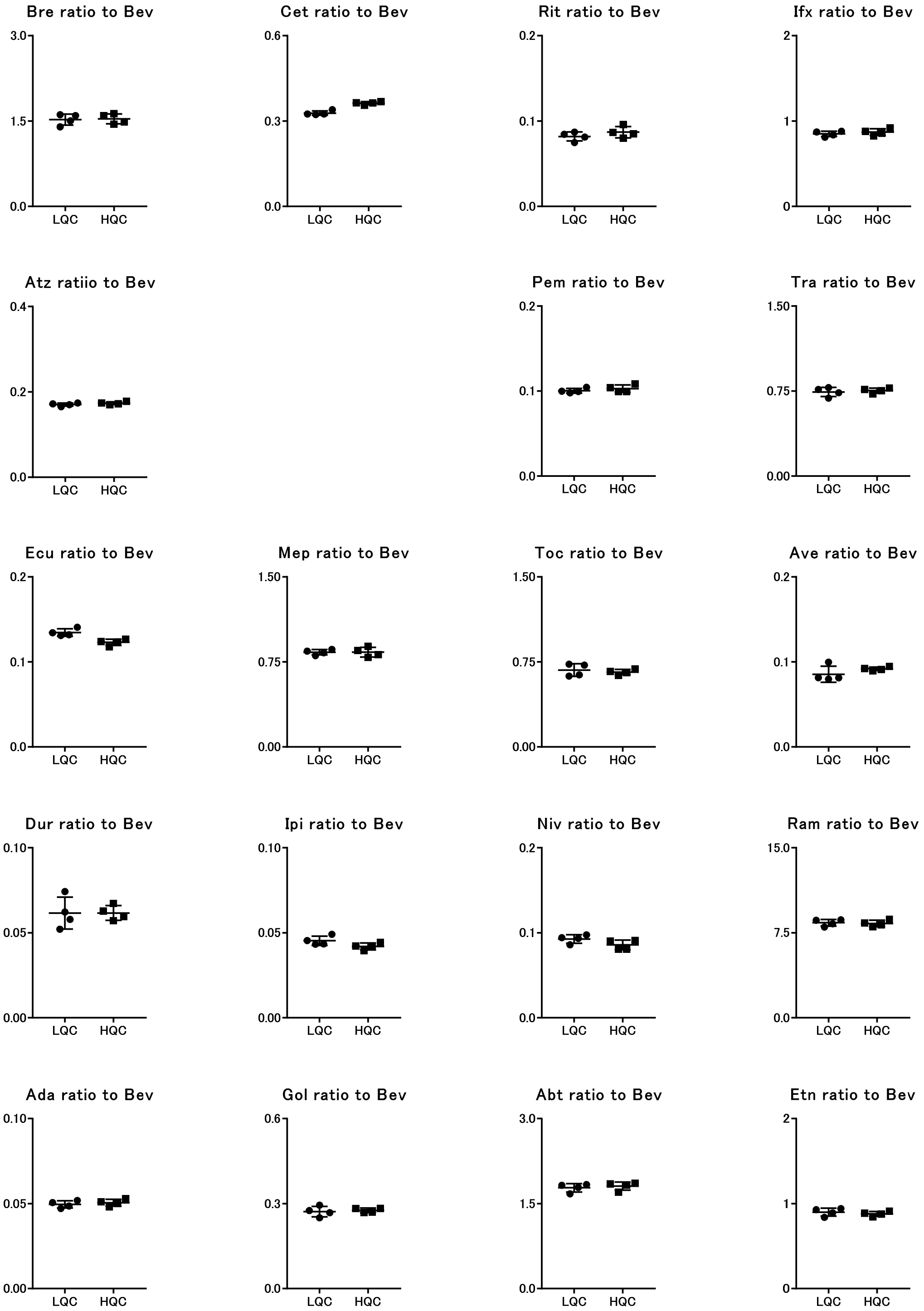

**G)** The ratio to pembrolizumab

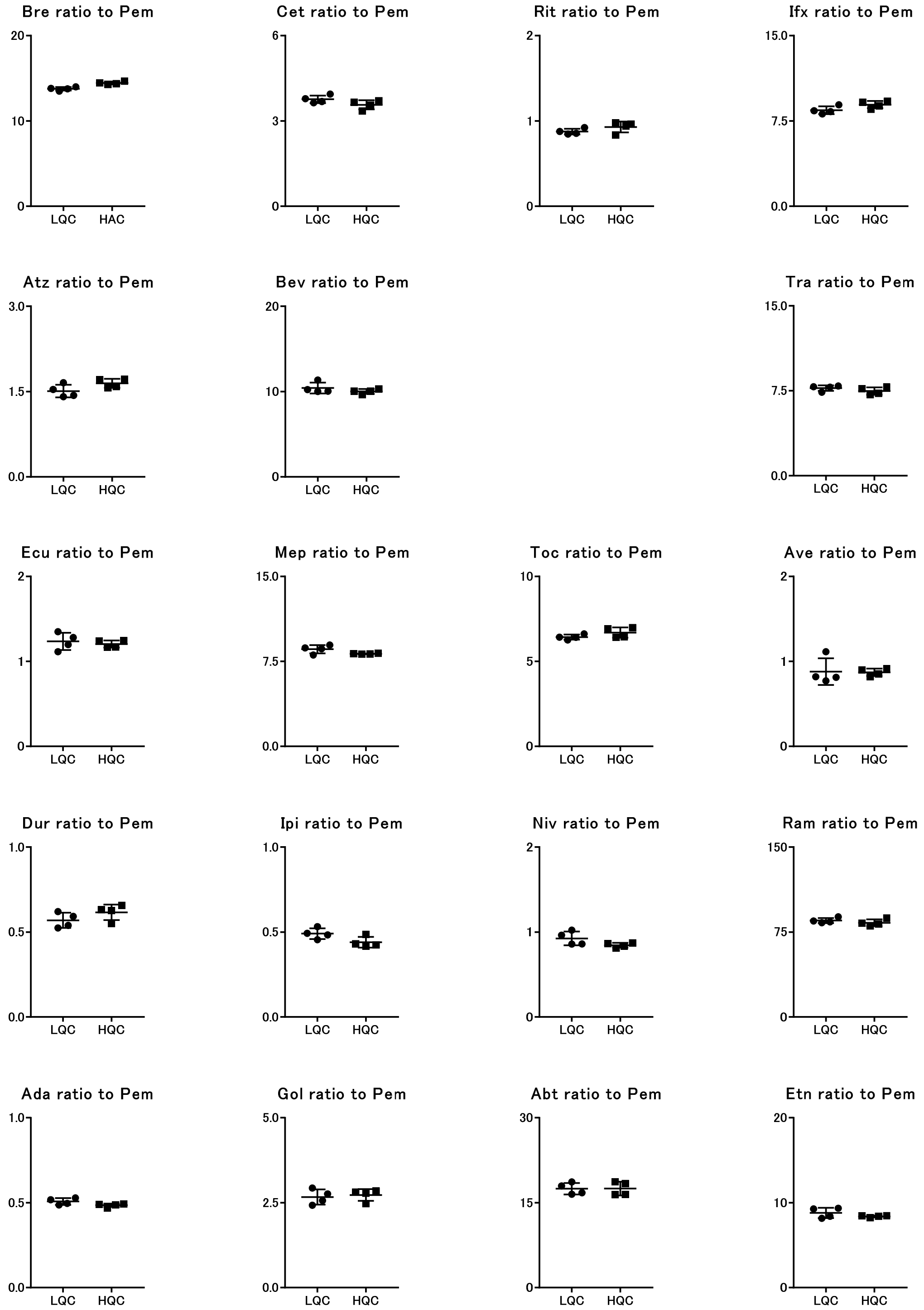

**H)** The ratio to eculizumab

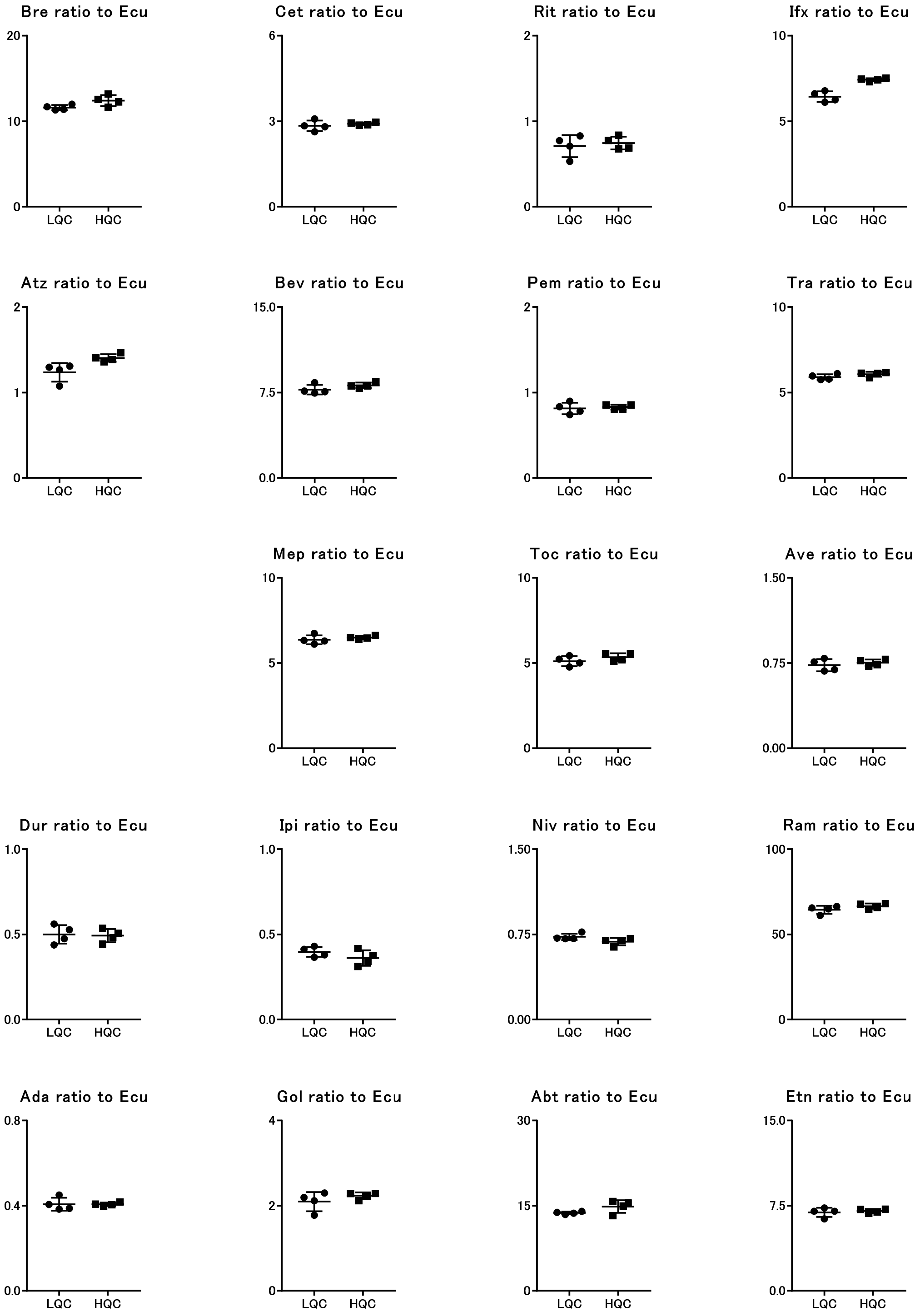

**I)** The ratio to mepolizumab

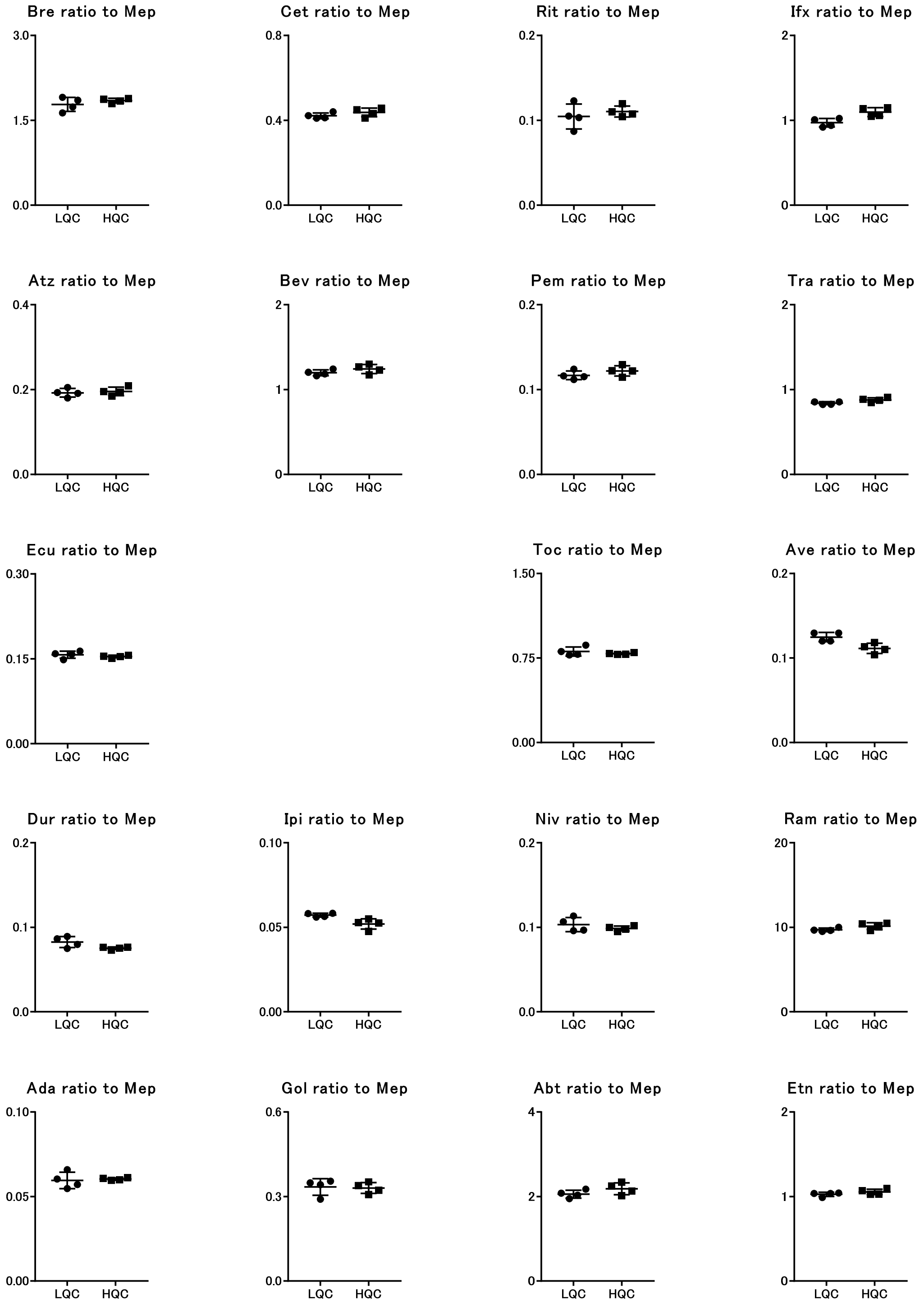

**J)** The ratio to tocilizumab

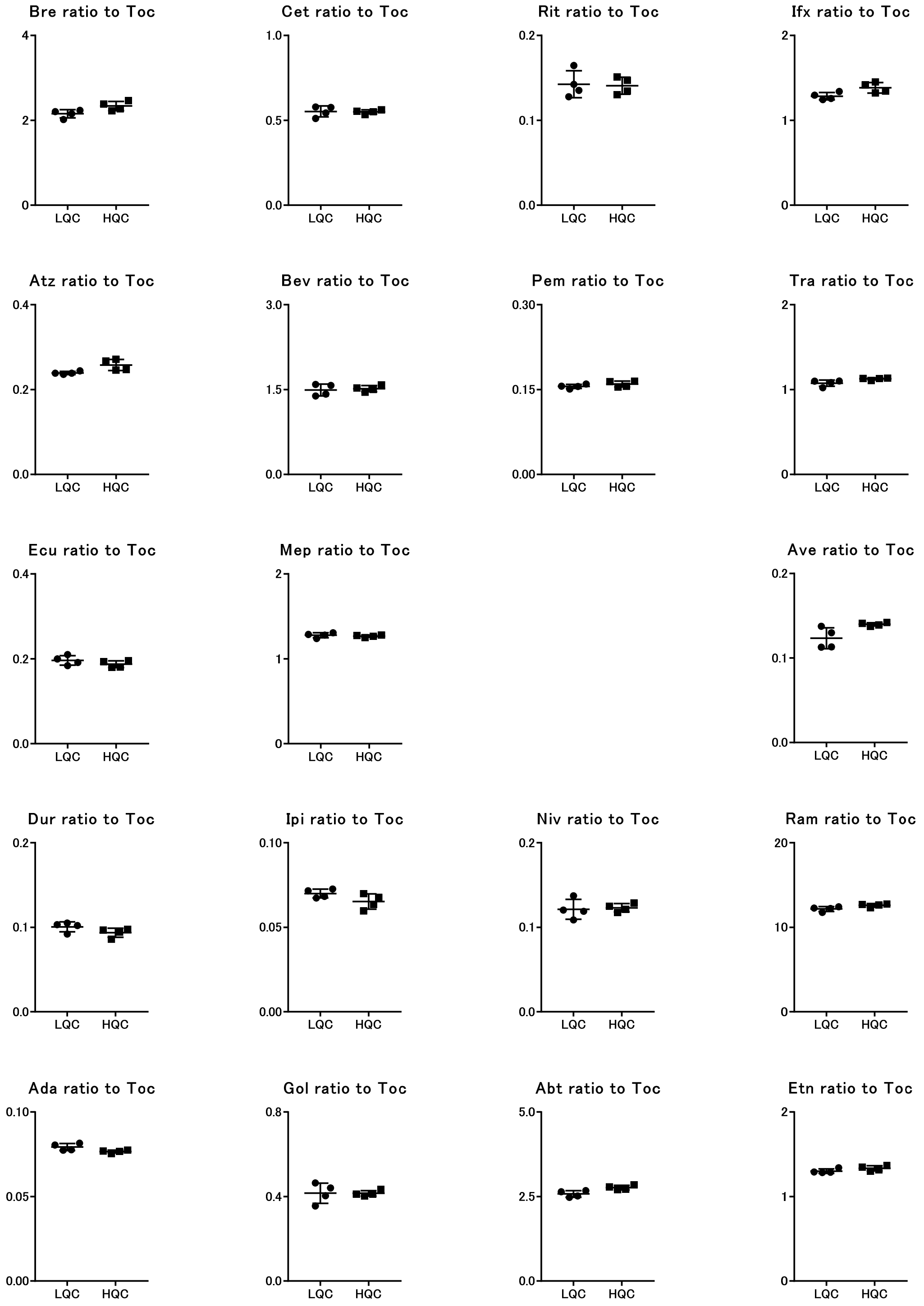

**K)** The ratio to avelumab

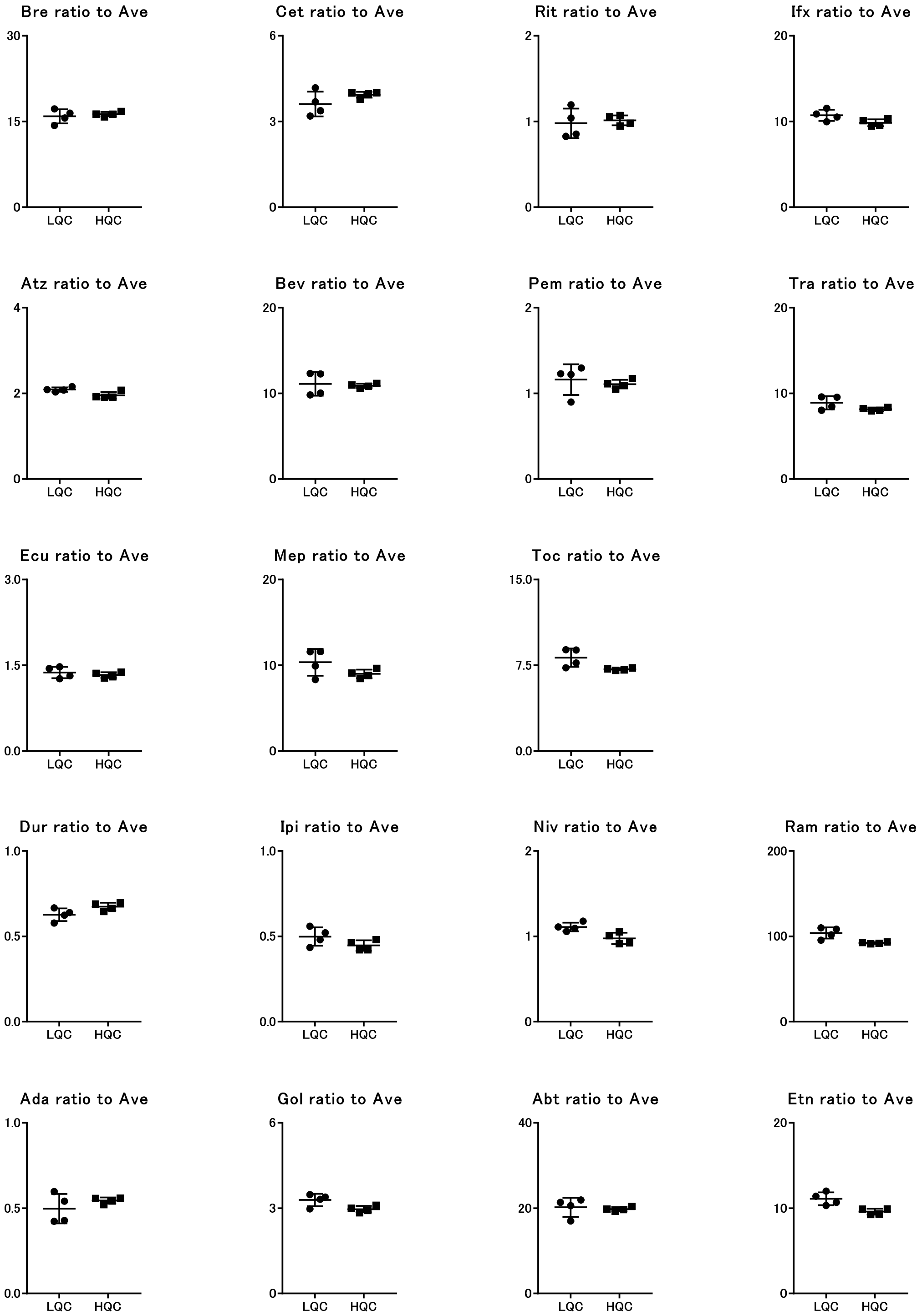

**L)** The ratio to durvalumab

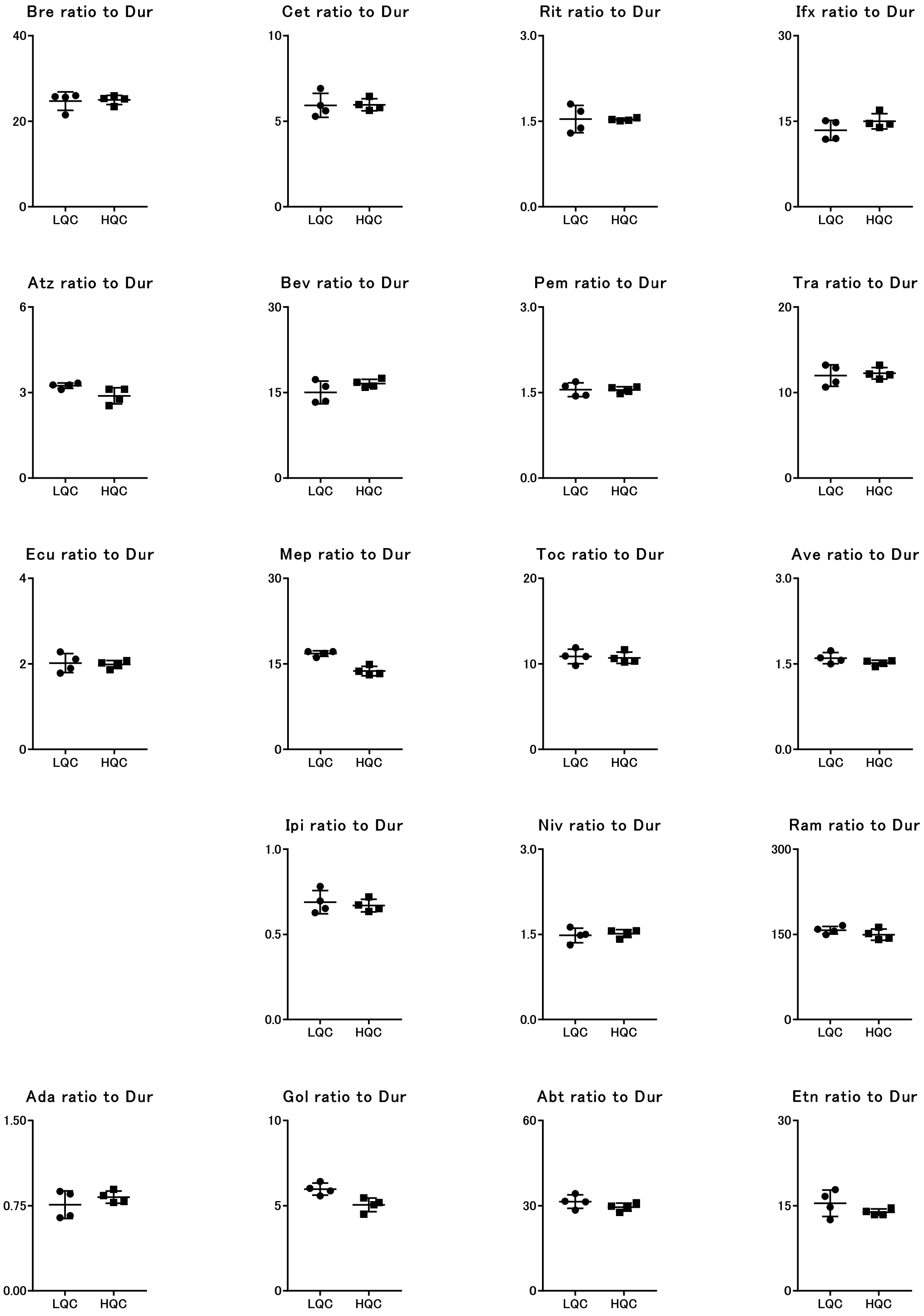

**M)** The ratio to ipilimumab

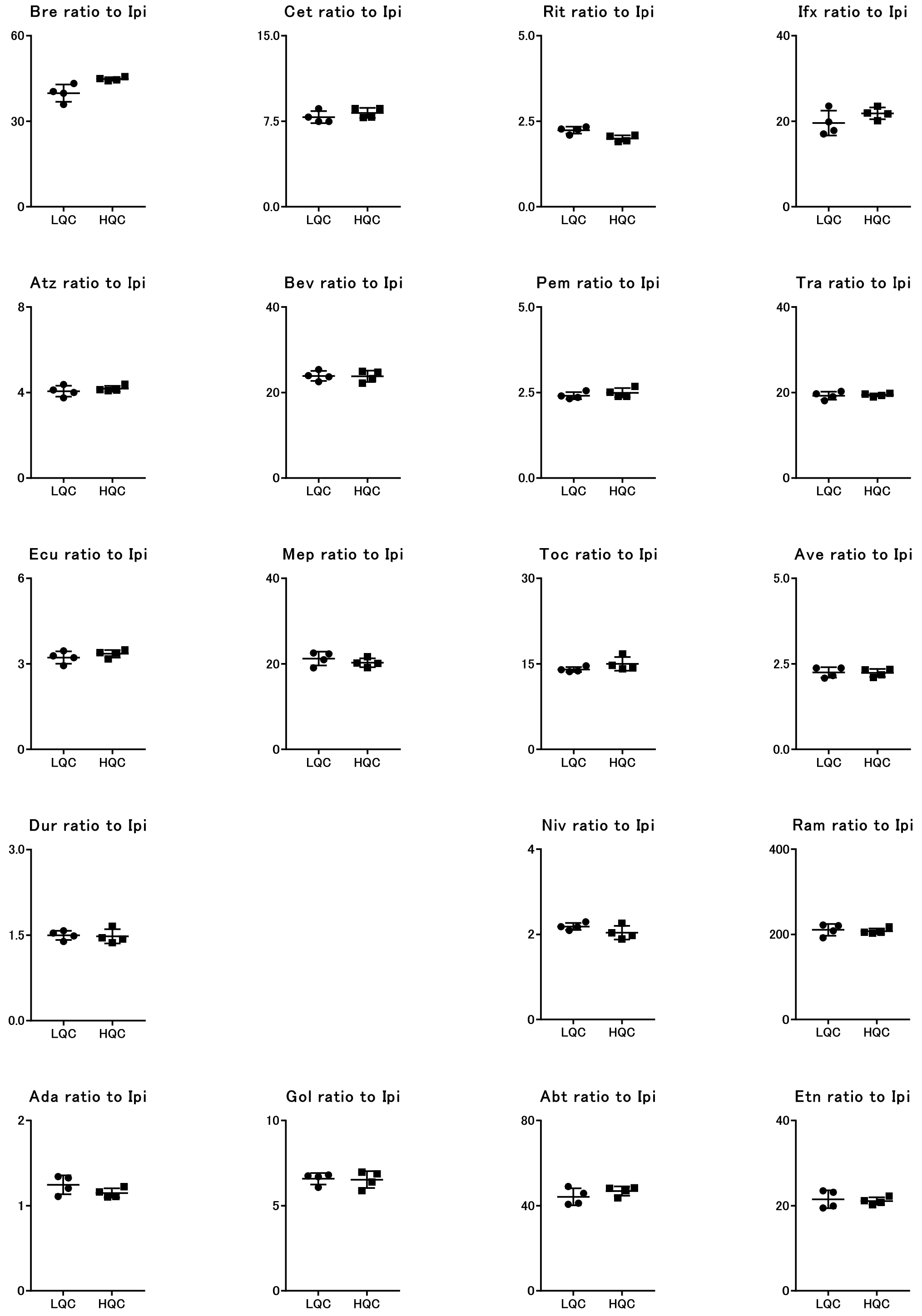

**N)** The ratio to nivolumab

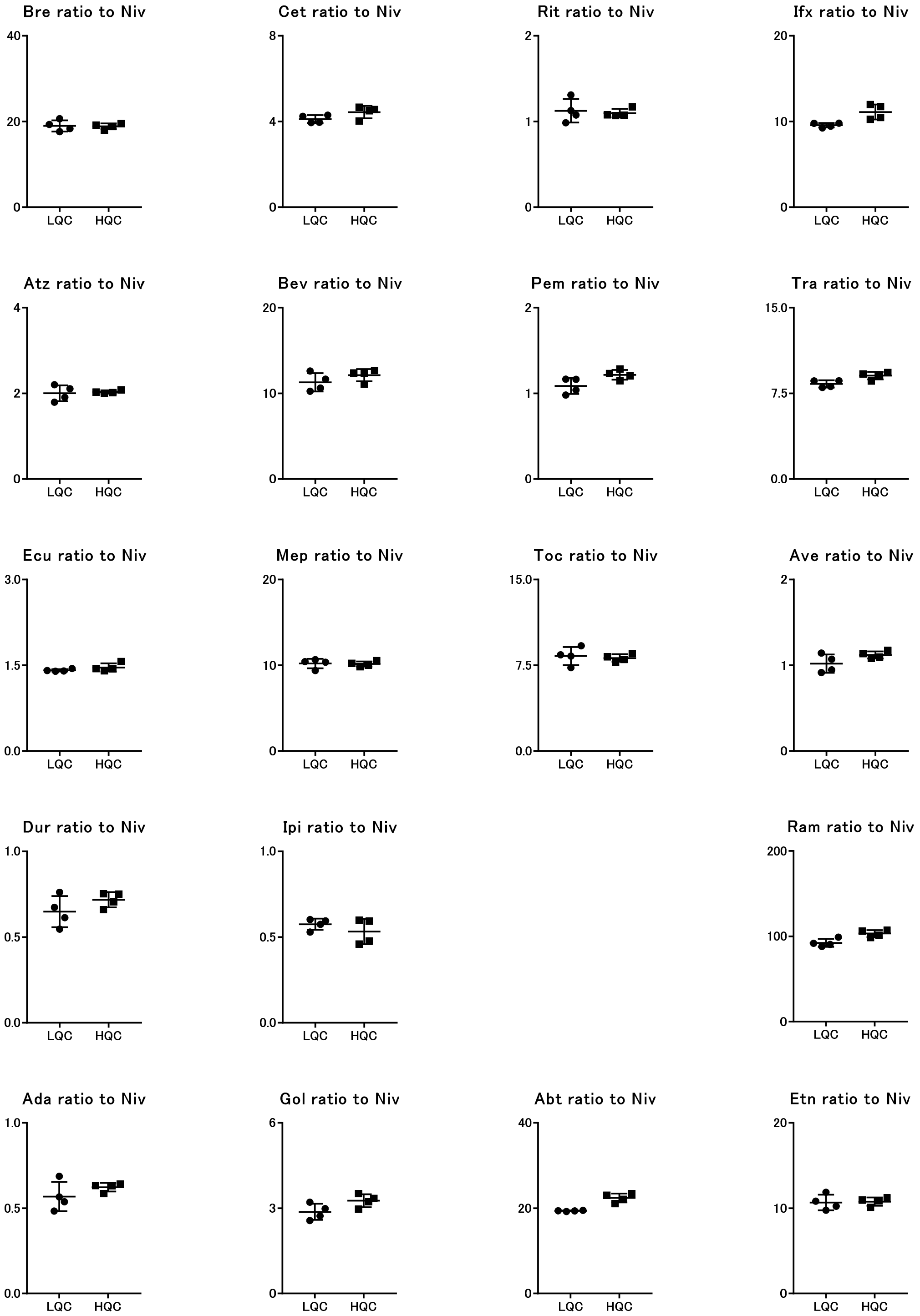

**O)** The ratio to ramucilumab

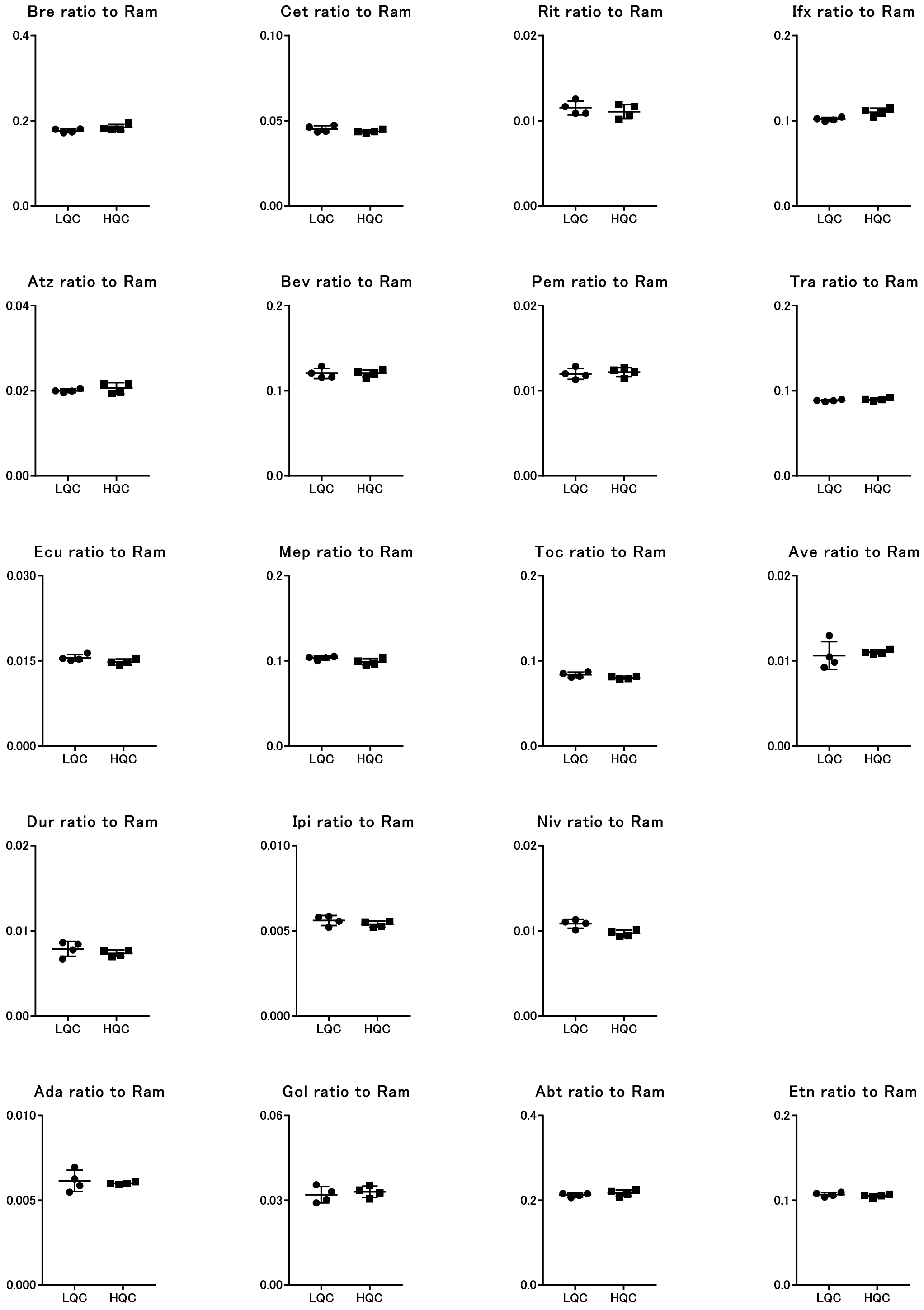

**P)** The ratio to adalimumab

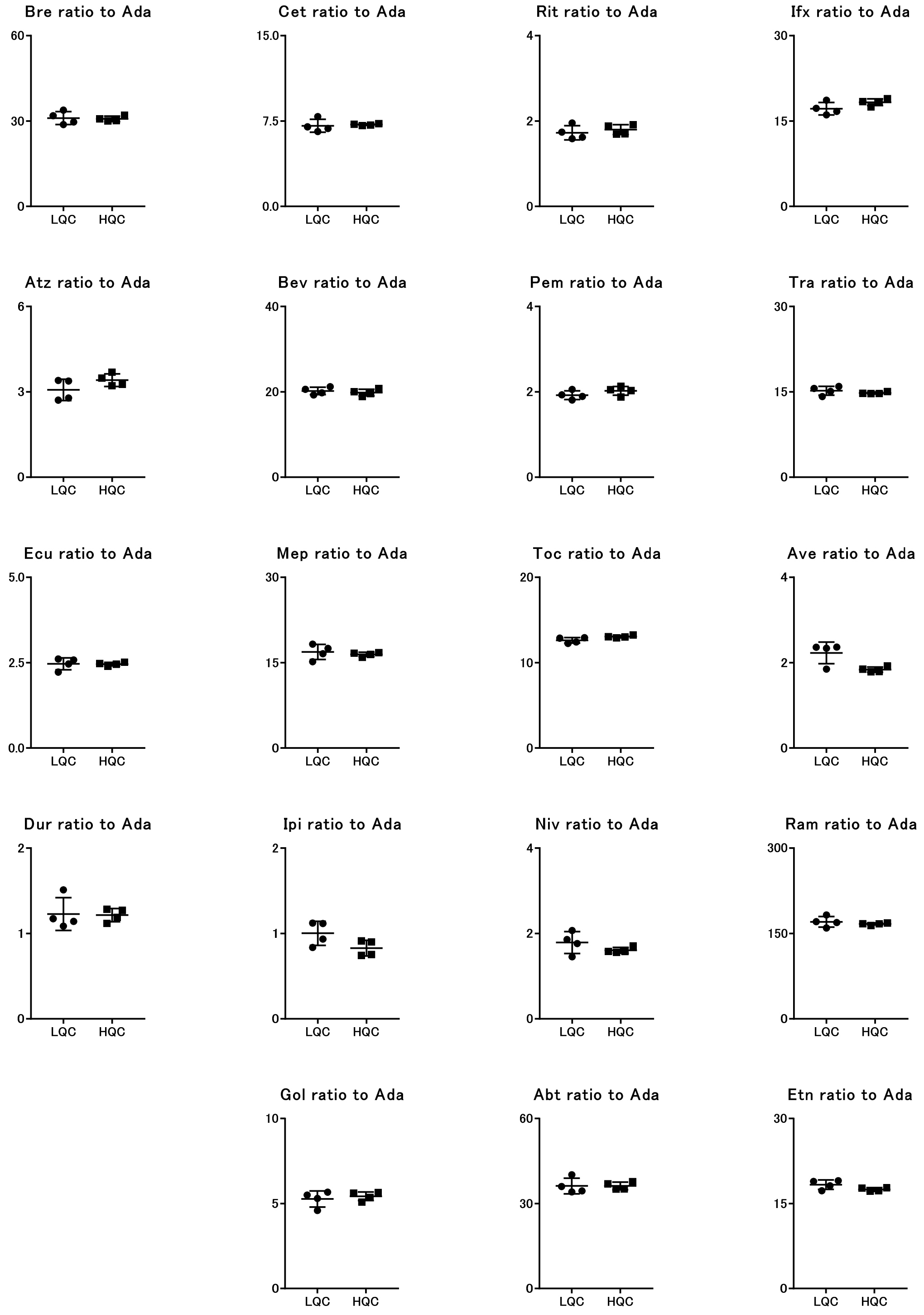

**Q)** The ratio to golimumab

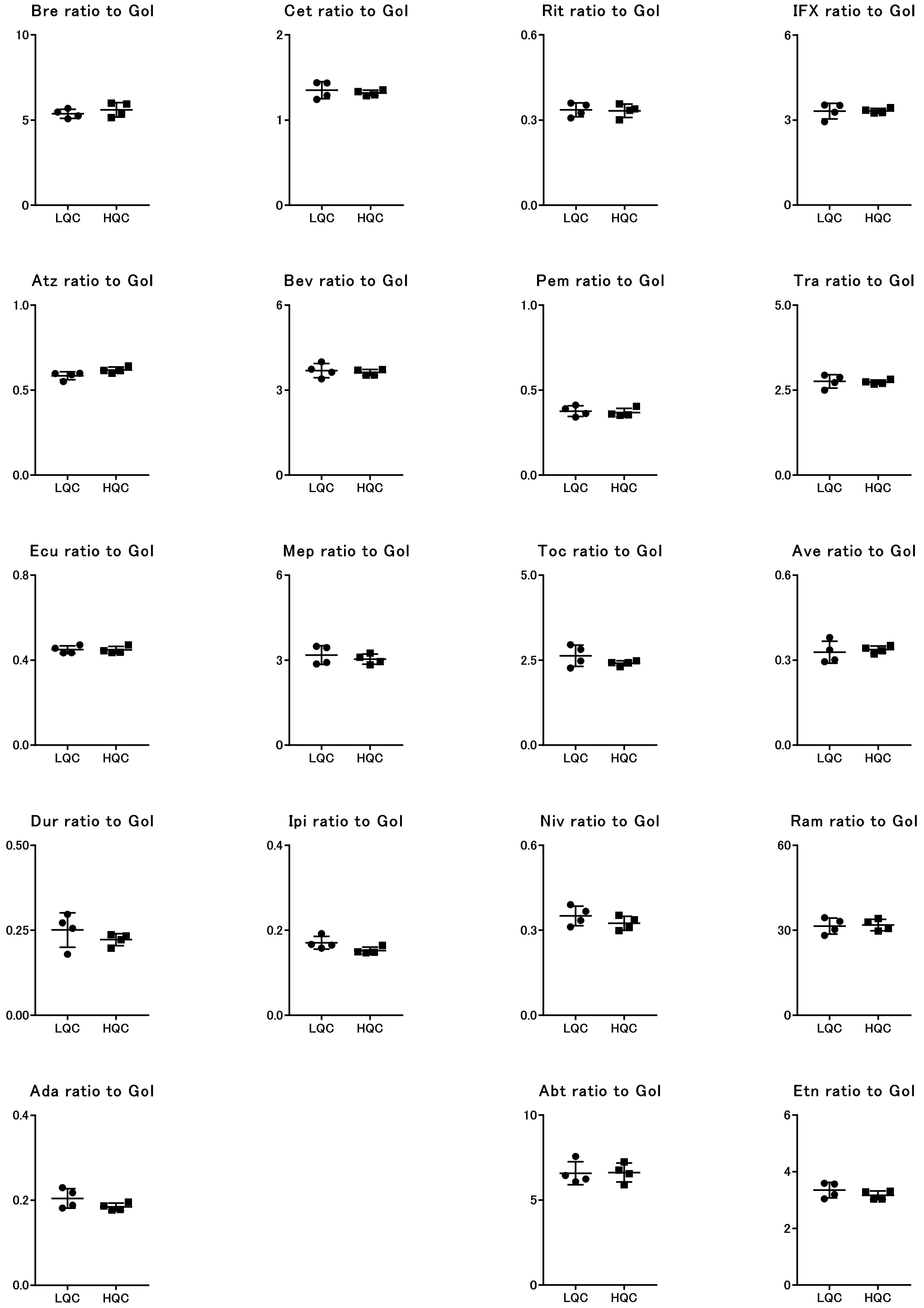

**R)** The ratio to abatacept

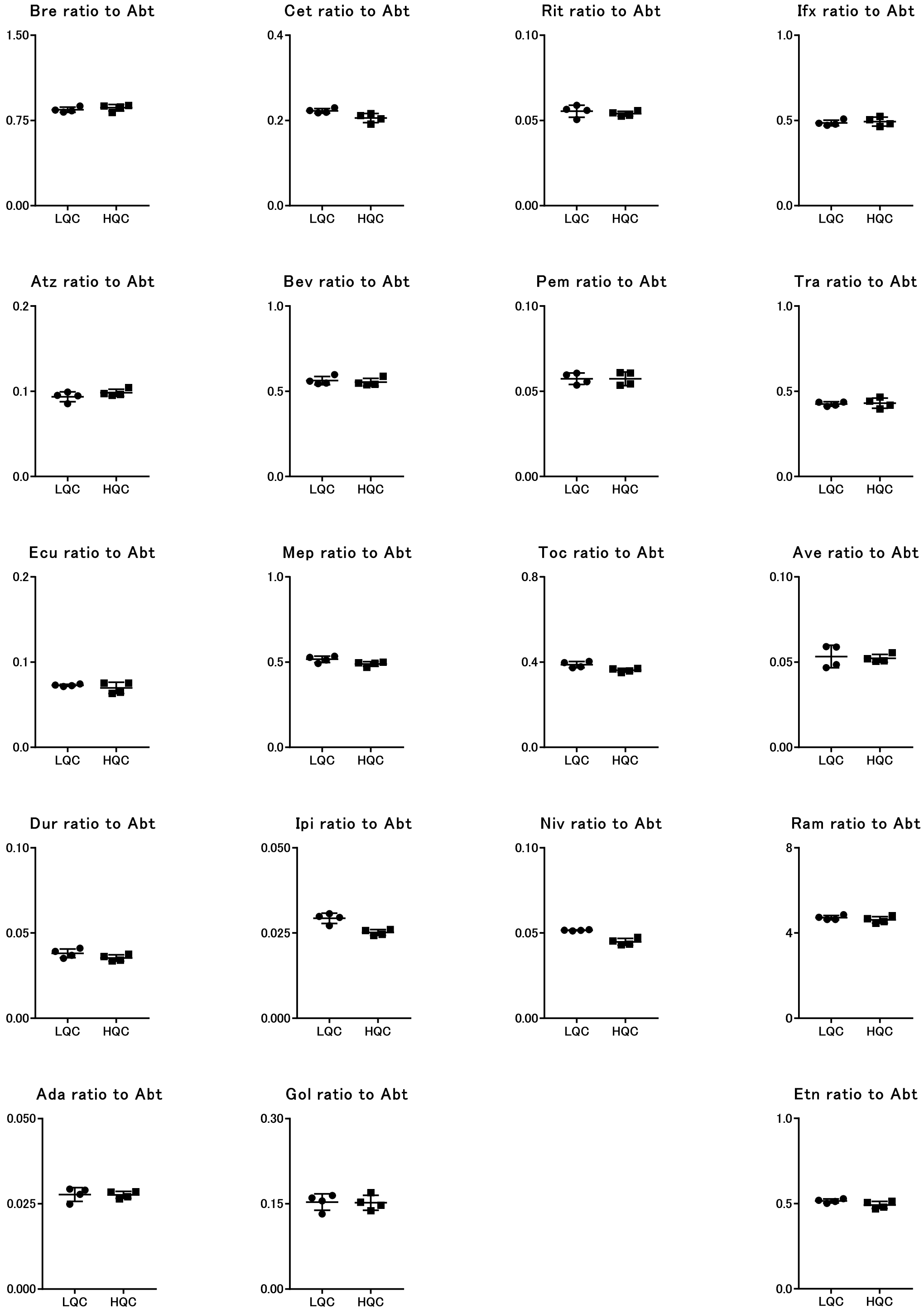

**S)** The ratio to etanercept

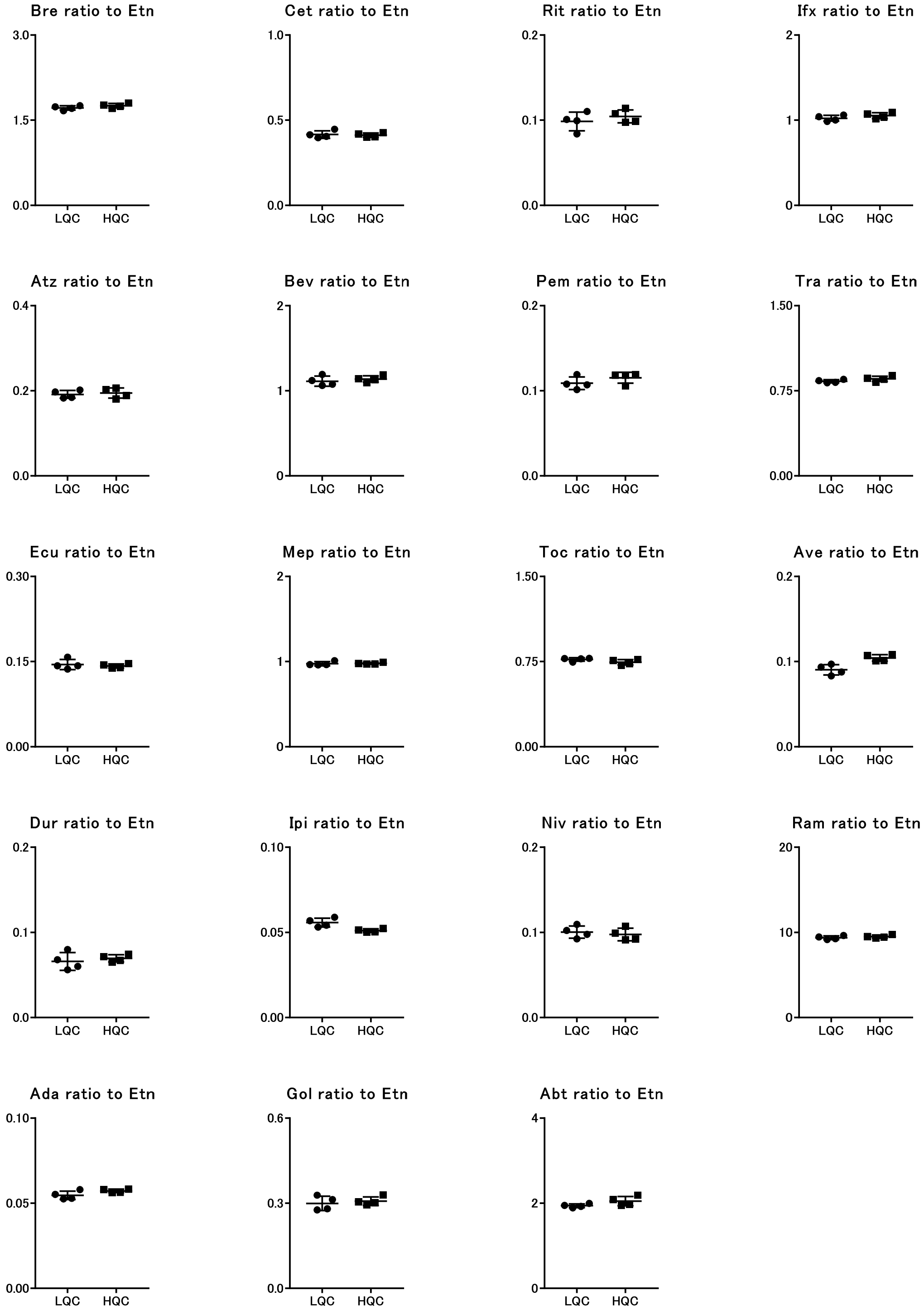

**Figure S4. The CPS ratio between each signature peptide of mAbs at low QC and high QC concentration points.** The CPS value ratio of each monoclonal antibody are shown (n = 4) for the ratio to a) Bre, b) Cet, c) Rit, d) Ifx, e) Atz, f) Bev, g) Pem, h) Ecu, i) Mep, j) Toc, k) Ave, l) Dur, m) Ipi, n) Niv, o) Ram, p) Ada, q) Gol, r) Abt, and s) Etn as a reference Ab. The y axis shows the CPS ratio (each Ab/reference Ab).

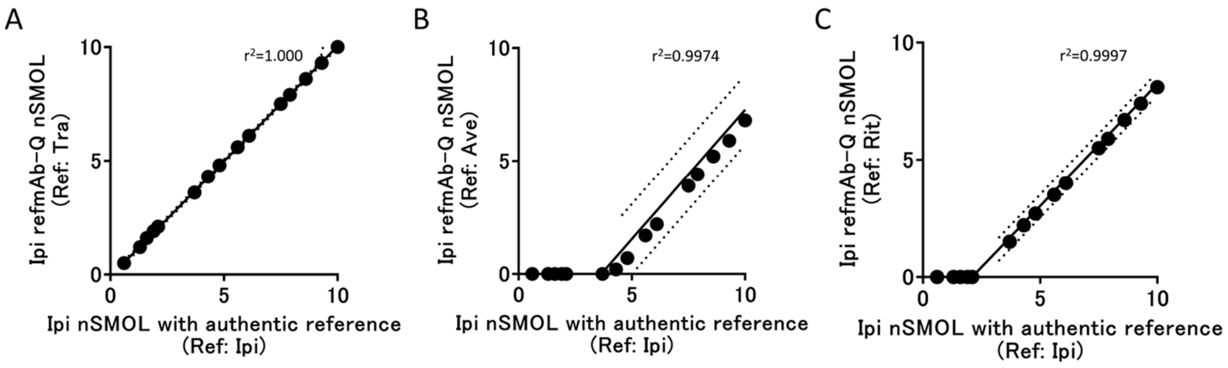


**Figure S5. Concordance between the conventional nSMOL and refmAb-Q nSMOL using preferential choice and low-sensitivity references.** Concordance between the conventional nSMOL assay (x-axis) and refmAb-Q nSMOL assay (y-axis) with A) Tra, B) Ave, or C) Rit as a reference mAb for the quantitation of ipilimumab. Tra was classified as a preferential choice mAb whereas Ave and Rit were designated as low sensitivity mAbs in Figure 3.

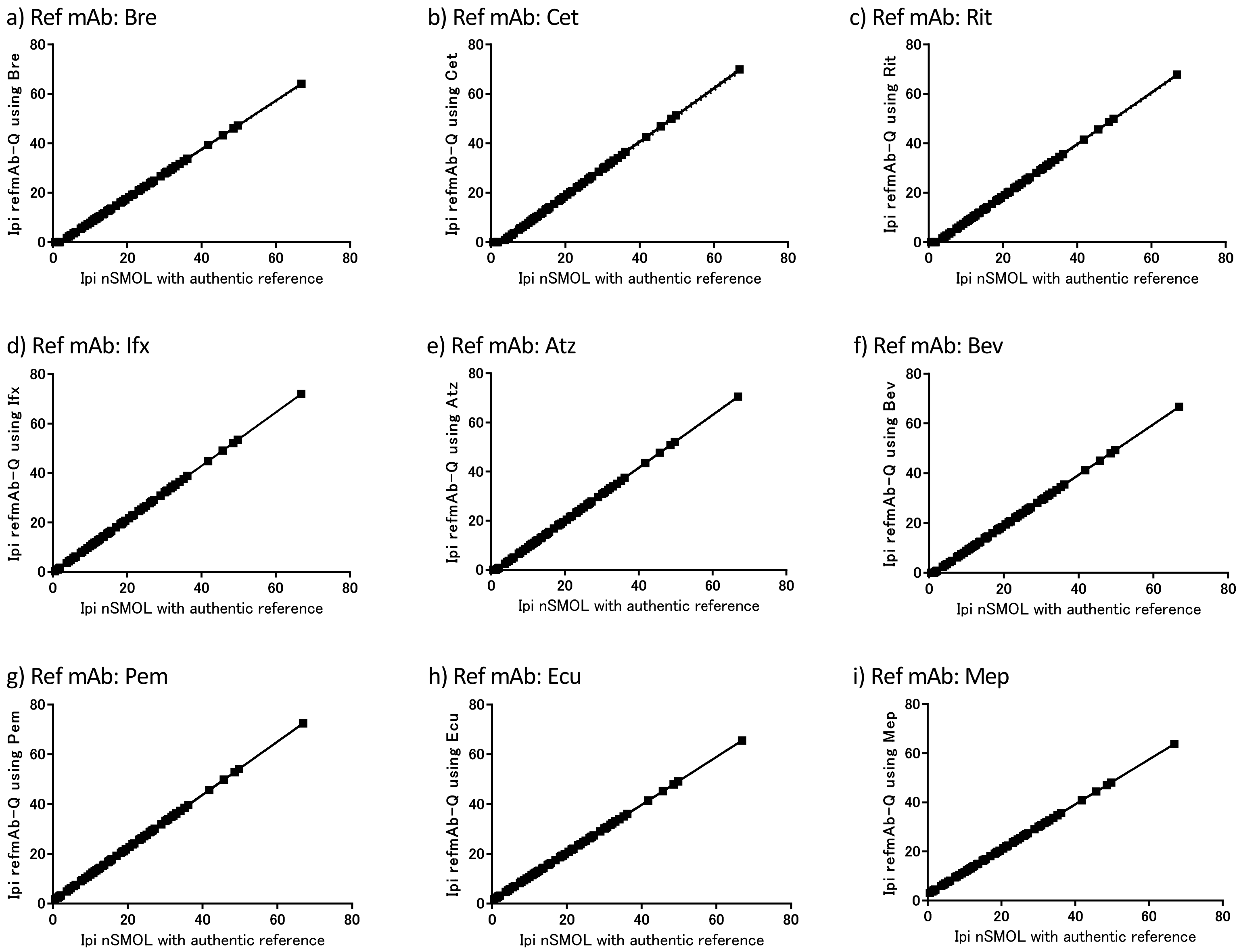
a) Ref mAb: Bre
Ipi refmAb-Q using Bre
Ipi nSMOL with authentic reference
b) Ref mAb: Cet
Ipi refmAb-Q using Cet
Ipi nSMOL with authentic reference
c) Ref mAb: Rit
Ipi refmAb-Q using Rit
Ipi nSMOL with authentic reference
d) Ref mAb: Ifx
Ipi refmAb-Q using Ifx
Ipi nSMOL with authentic reference
e) Ref mAb: Atz
Ipi refmAb-Q using Atz
Ipi nSMOL with authentic reference
f) Ref mAb: Bev
Ipi refmAb-Q using Bev
Ipi nSMOL with authentic reference
g) Ref mAb: Pem
Ipi refmAb-Q using Pem
Ipi nSMOL with authentic reference
h) Ref mAb: Ecu
Ipi refmAb-Q using Ecu
Ipi nSMOL with authentic reference
i) Ref mAb: Mep
Ipi refmAb-Q using Mep
Ipi nSMOL with authentic reference

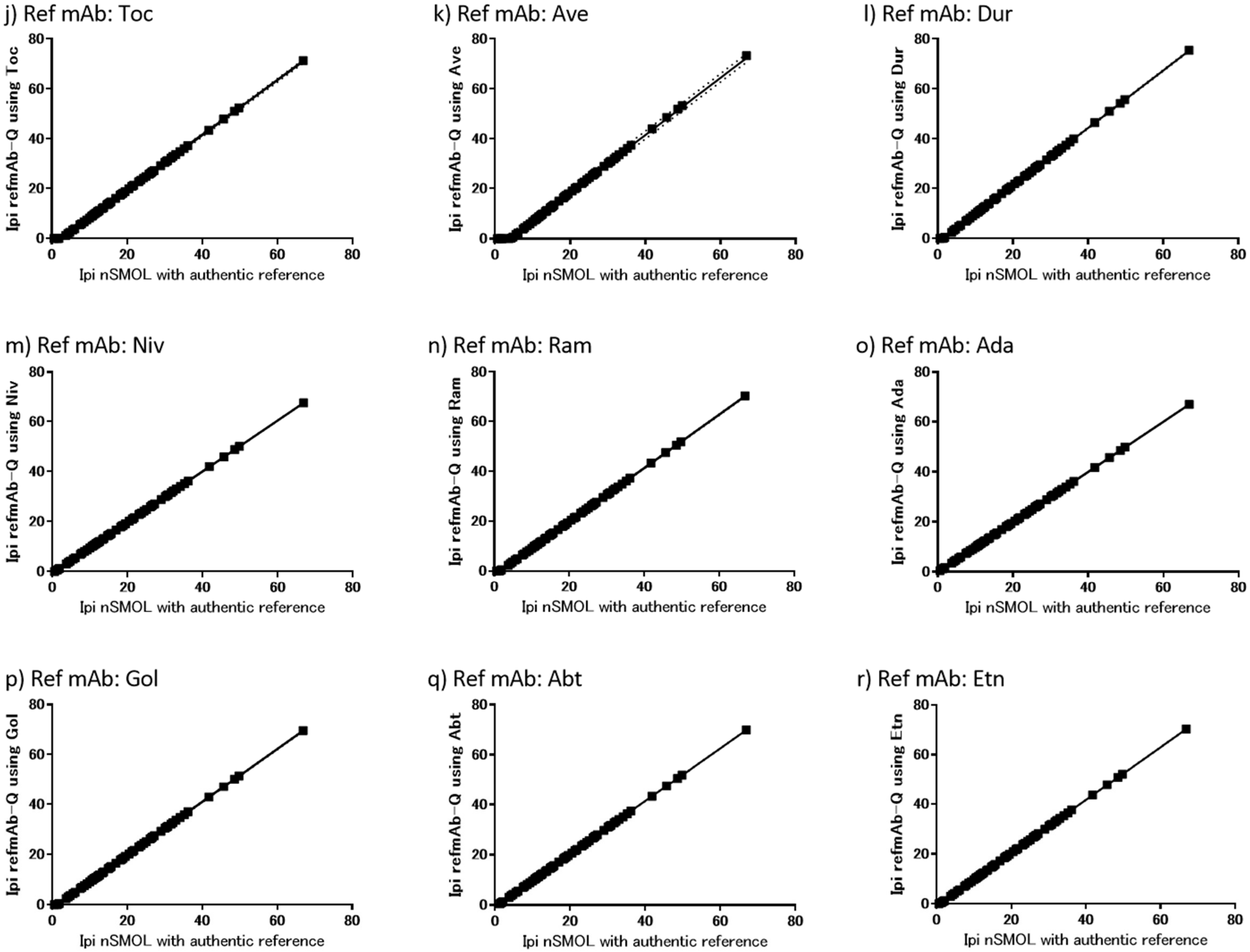
j) Ref mAb: Toc
Ipi refmAb-Q using Toc
Ipi nSMOL with authentic reference
k) Ref mAb: Ave
Ipi refmAb-Q using Ave
Ipi nSMOL with authentic reference
l) Ref mAb: Dur
Ipi refmAb-Q using Dur
Ipi nSMOL with authentic reference
m) Ref mAb: Niv
Ipi refmAb-Q using Niv
Ipi nSMOL with authentic reference
n) Ref mAb: Ram
Ipi refmAb-Q using Ram
Ipi nSMOL with authentic reference
o) Ref mAb: Ada
Ipi refmAb-Q using Ada
Ipi nSMOL with authentic reference
p) Ref mAb: Gol
Ipi refmAb-Q using Gol
Ipi nSMOL with authentic reference
q) Ref mAb: Abt
Ipi refmAb-Q using Abt
Ipi nSMOL with authentic reference
r) Ref mAb: Etn
Ipi refmAb-Q using Etn
Ipi nSMOL with authentic reference

**Figure S6. Pearson correlation analyses between nSMOL assay data with the authentic reference and refmAb-Q nSMOL assay data with the indicated reference antibody for clinical serum samples from patients treated with ipilimumab (n=71).** The reference antibody used was a) Bre, b) Cet, c) Rit, d) Ifx, e) Atz, f) Bev, g) Pem, h) Ecu, i) Mep, j) Toc, k) Ave, l) Dur, m) Niv, n) Ram, o) Ada, p) Gol, q) Abt, or r) Etn, respectively. The dashed line shows the 95% confidence interval. The coefficients are summarized in Table S4a.

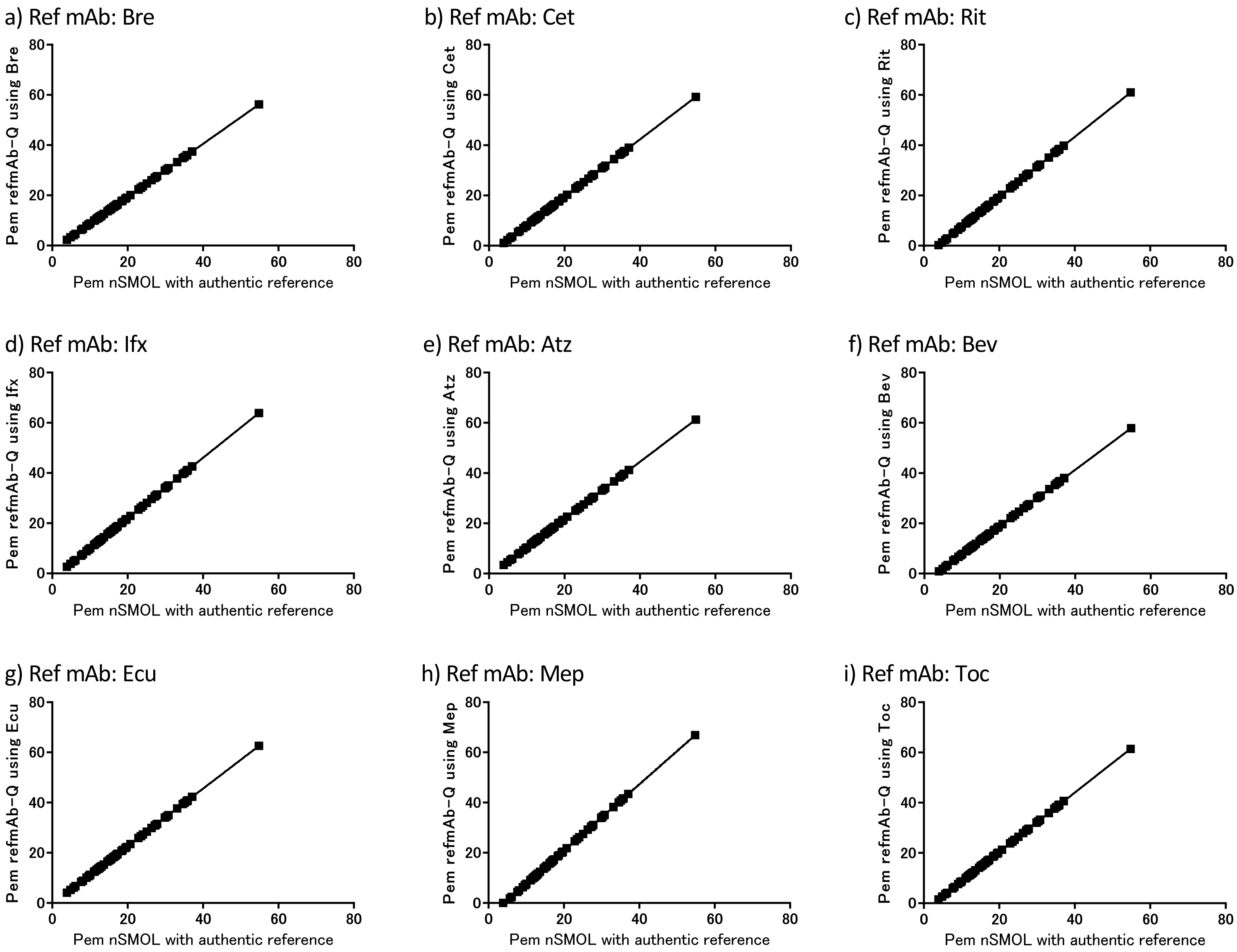
a) Ref mAb: Bre
Pem refmAb-Q using Bre
Pem nSMOL with authentic reference
b) Ref mAb: Cet
Pem refmAb-Q using Cet
Pem nSMOL with authentic reference
c) Ref mAb: Rit
Pem refmAb-Q using Rit
Pem nSMOL with authentic reference
d) Ref mAb: Ifx
Pem refmAb-Q using Ifx
Pem nSMOL with authentic reference
e) Ref mAb: Atz
Pem refmAb-Q using Atz
Pem nSMOL with authentic reference
f) Ref mAb: Bev
Pem refmAb-Q using Bev
Pem nSMOL with authentic reference
g) Ref mAb: Ecu
Pem refmAb-Q using Ecu
Pem nSMOL with authentic reference
h) Ref mAb: Mep
Pem refmAb-Q using Mep
Pem nSMOL with authentic reference
i) Ref mAb: Toc
Pem refmAb-Q using Toc
Pem nSMOL with authentic reference
0
20
40
60
80

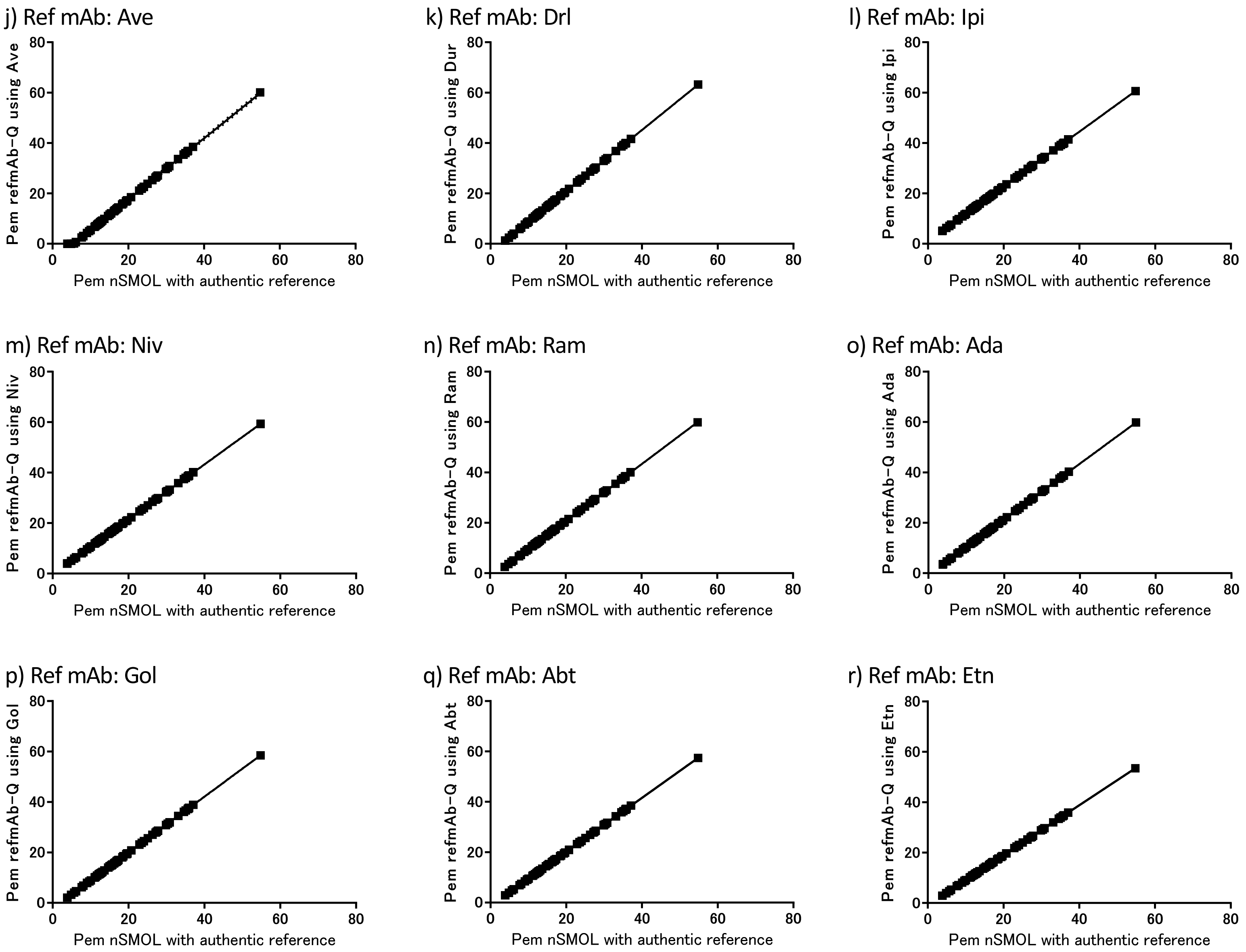
j) Ref mAb: Ave
Pem refmAb-Q using Ave
Pem nSMOL with authentic reference
k) Ref mAb: Drl
Pem refmAb-Q using Dur
Pem nSMOL with authentic reference
l) Ref mAb: Ipi
Pem refmAb-Q using Ipi
Pem nSMOL with authentic reference
m) Ref mAb: Niv
Pem refmAb-Q using Niv
Pem nSMOL with authentic reference
n) Ref mAb: Ram
Pem refmAb-Q using Ram
Pem nSMOL with authentic reference
o) Ref mAb: Ada
Pem refmAb-Q using Ada
Pem nSMOL with authentic reference
p) Ref mAb: Gol
Pem refmAb-Q using Gol
Pem nSMOL with authentic reference
q) Ref mAb: Abt
Pem refmAb-Q using Abt
Pem nSMOL with authentic reference
r) Ref mAb: Etn
Pem refmAb-Q using Etn
Pem nSMOL with authentic reference

**Figure S7. Pearson correlation analyses between nSMOL assay data with the authentic reference and refmAb-Q nSMOL assay data with the indicated reference antibody for clinical serum samples from patients treated with pembrolizumab (n=74).** The reference antibody used was a) Bre, b) Cet, c) Rit, d) Ifx, e) Atz, f) Bev, g) Ecu, h) Mep, i) Toc, j) Ave, k) Dur, l) Ipi, m) Niv, n) Ram, o) Ada, p) Gol, q) Abt, or r) Etn, respectively. The dashed line shows the 95% confidence interval. The coefficients are summarized in Table S4b.

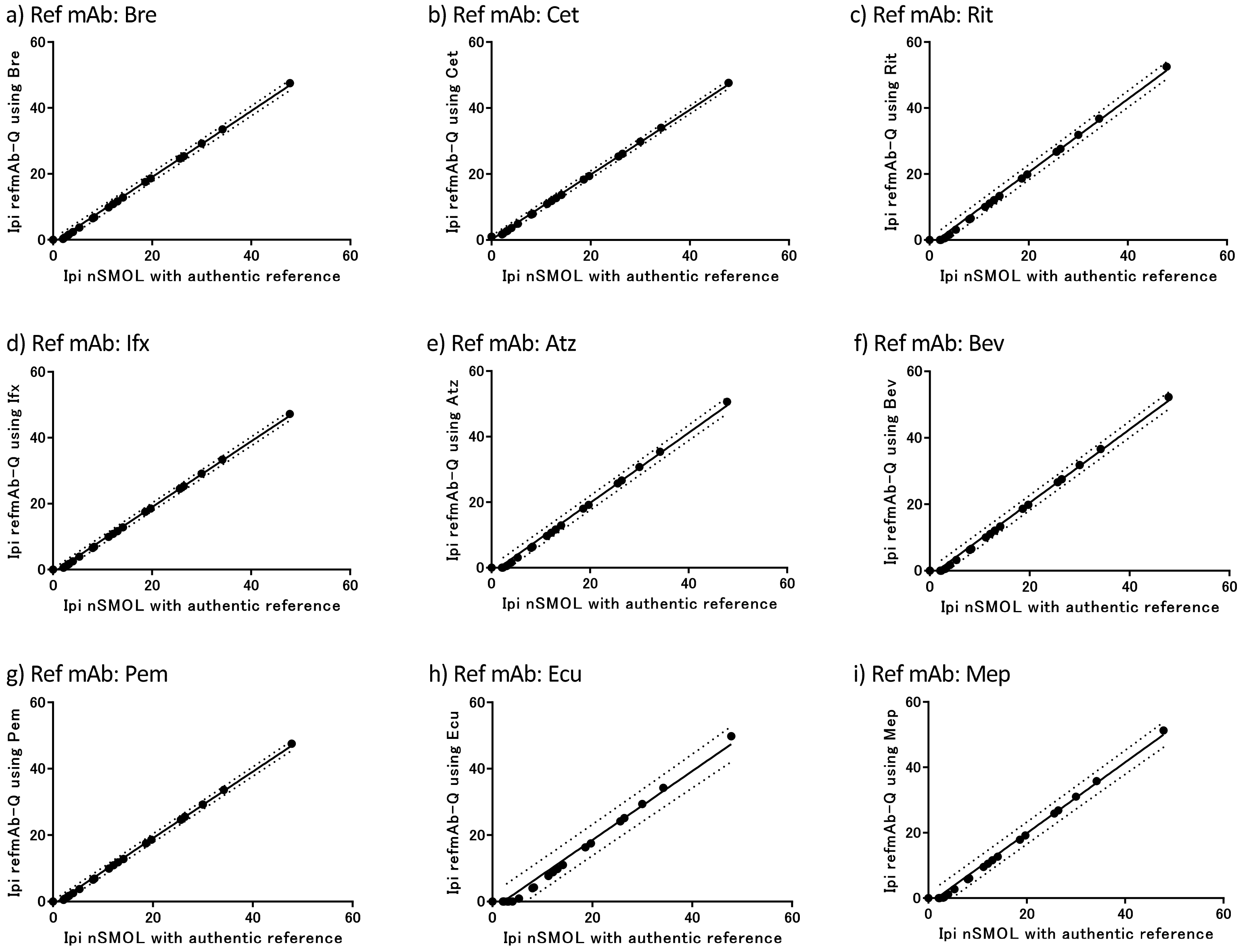
a) Ref mAb: Bre
Ipi refmAb-Q using Bre
Ipi nSMOL with authentic reference
b) Ref mAb: Cet
Ipi refmAb-Q using Cet
Ipi nSMOL with authentic reference
c) Ref mAb: Rit
Ipi refmAb-Q using Rit
Ipi nSMOL with authentic reference
d) Ref mAb: Ifx
Ipi refmAb-Q using Ifx
Ipi nSMOL with authentic reference
e) Ref mAb: Atz
Ipi refmAb-Q using Atz
Ipi nSMOL with authentic reference
f) Ref mAb: Bev
Ipi refmAb-Q using Bev
Ipi nSMOL with authentic reference
g) Ref mAb: Pem
Ipi refmAb-Q using Pem
Ipi nSMOL with authentic reference
h) Ref mAb: Ecu
Ipi refmAb-Q using Ecu
Ipi nSMOL with authentic reference
i) Ref mAb: Mep
Ipi refmAb-Q using Mep
Ipi nSMOL with authentic reference

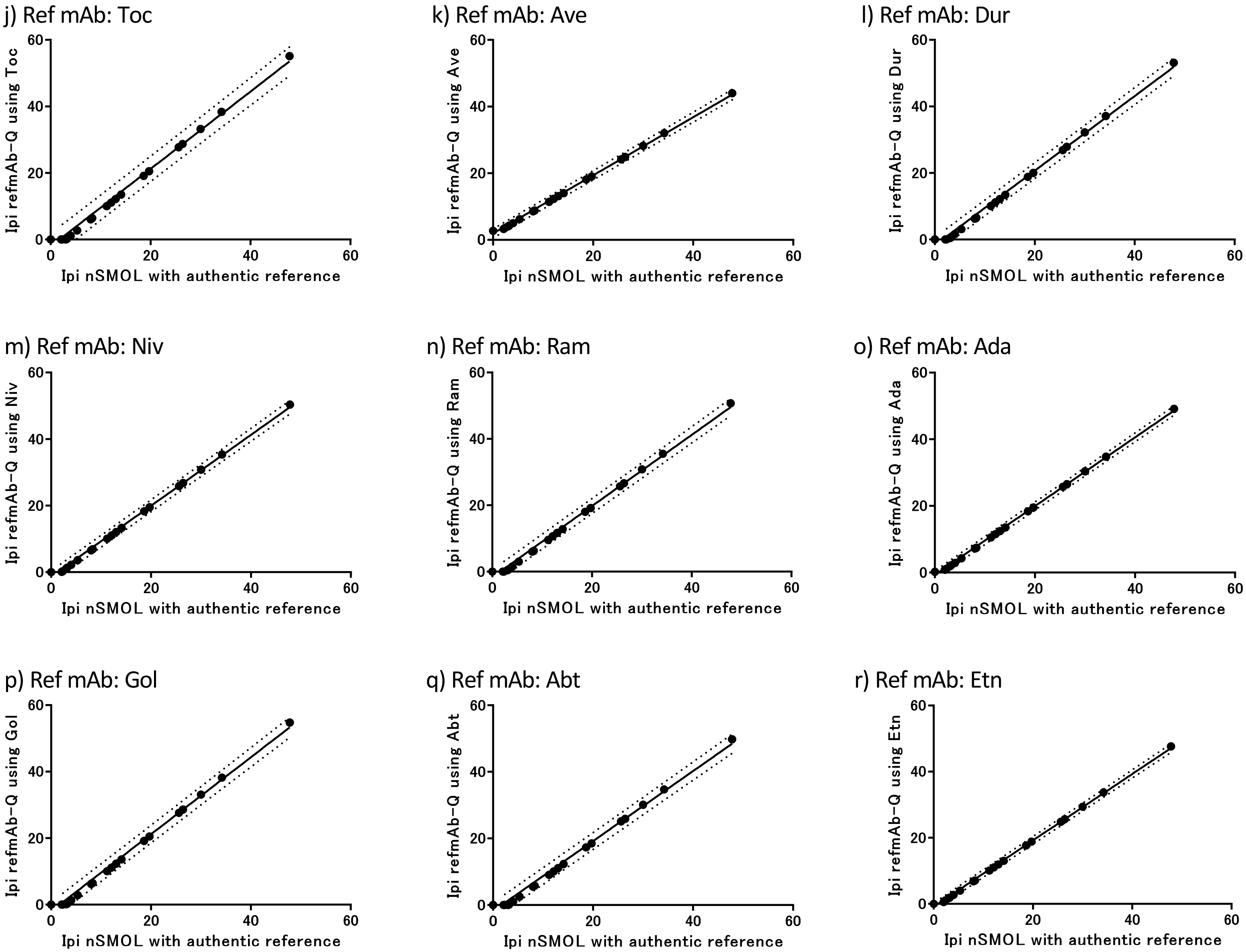
j) Ref mAb: Toc
Ipi refmAb-Q using Toc
Ipi nSMOL with authentic reference
k) Ref mAb: Ave
Ipi refmAb-Q using Ave
Ipi nSMOL with authentic reference
l) Ref mAb: Dur
Ipi refmAb-Q using Dur
Ipi nSMOL with authentic reference
m) Ref mAb: Niv
Ipi refmAb-Q using Niv
Ipi nSMOL with authentic reference
n) Ref mAb: Ram
Ipi refmAb-Q using Ram
Ipi nSMOL with authentic reference
o) Ref mAb: Ada
Ipi refmAb-Q using Ada
Ipi nSMOL with authentic reference
p) Ref mAb: Gol
Ipi refmAb-Q using Gol
Ipi nSMOL with authentic reference
q) Ref mAb: Abt
Ipi refmAb-Q using Abt
Ipi nSMOL with authentic reference
r) Ref mAb: Etn
Ipi refmAb-Q using Etn
Ipi nSMOL with authentic reference
0
20
40
60

**Figure S8. Pearson correlation analyses of ipilimumab data between nSMOL assay data with the authentic reference and refmAb-Q nSMOL assay data with the indicated reference antibody for clinical serum samples from patients treated with ipilimumab plus nivolumab combination therapy (n =27).** The reference antibody used was a) Bre, b) Cet, c) Rit, d) Ifx, e) Atz, f) Bev, g) Pem, h) Ecu, i) Mep, j) Toc, k) Ave, l) Dur, m) Niv, n) Ram, o) Ada, p) Gol, q) Abt, or r) Etn, respectively. The dashed line shows the 95% confidence interval. The coefficients are summarized in Table S4c.

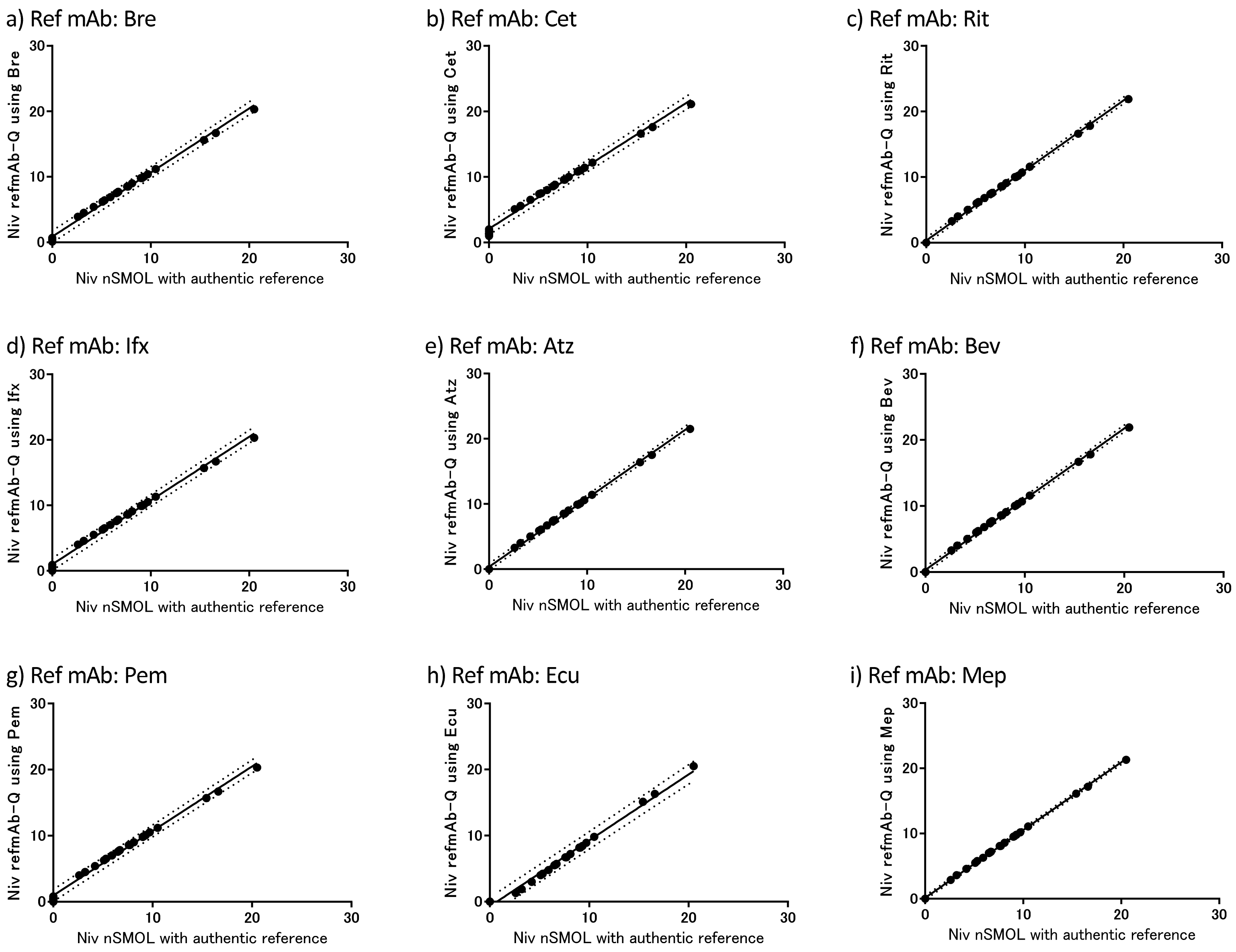
a) Ref mAb: Bre
Niv refmAb-Q using Bre
Niv nSMOL with authentic reference
b) Ref mAb: Cet
Niv refmAb-Q using Cet
Niv nSMOL with authentic reference
c) Ref mAb: Rit
Niv refmAb-Q using Rit
Niv nSMOL with authentic reference
d) Ref mAb: Ifx
Niv refmAb-Q using Ifx
Niv nSMOL with authentic reference
e) Ref mAb: Atz
Niv refmAb-Q using Atz
Niv nSMOL with authentic reference
f) Ref mAb: Bev
Niv refmAb-Q using Bev
Niv nSMOL with authentic reference
g) Ref mAb: Pem
Niv refmAb-Q using Pem
Niv nSMOL with authentic reference
h) Ref mAb: Ecu
Niv refmAb-Q using Ecu
Niv nSMOL with authentic reference
i) Ref mAb: Mep
Niv refmAb-Q using Mep
Niv nSMOL with authentic reference

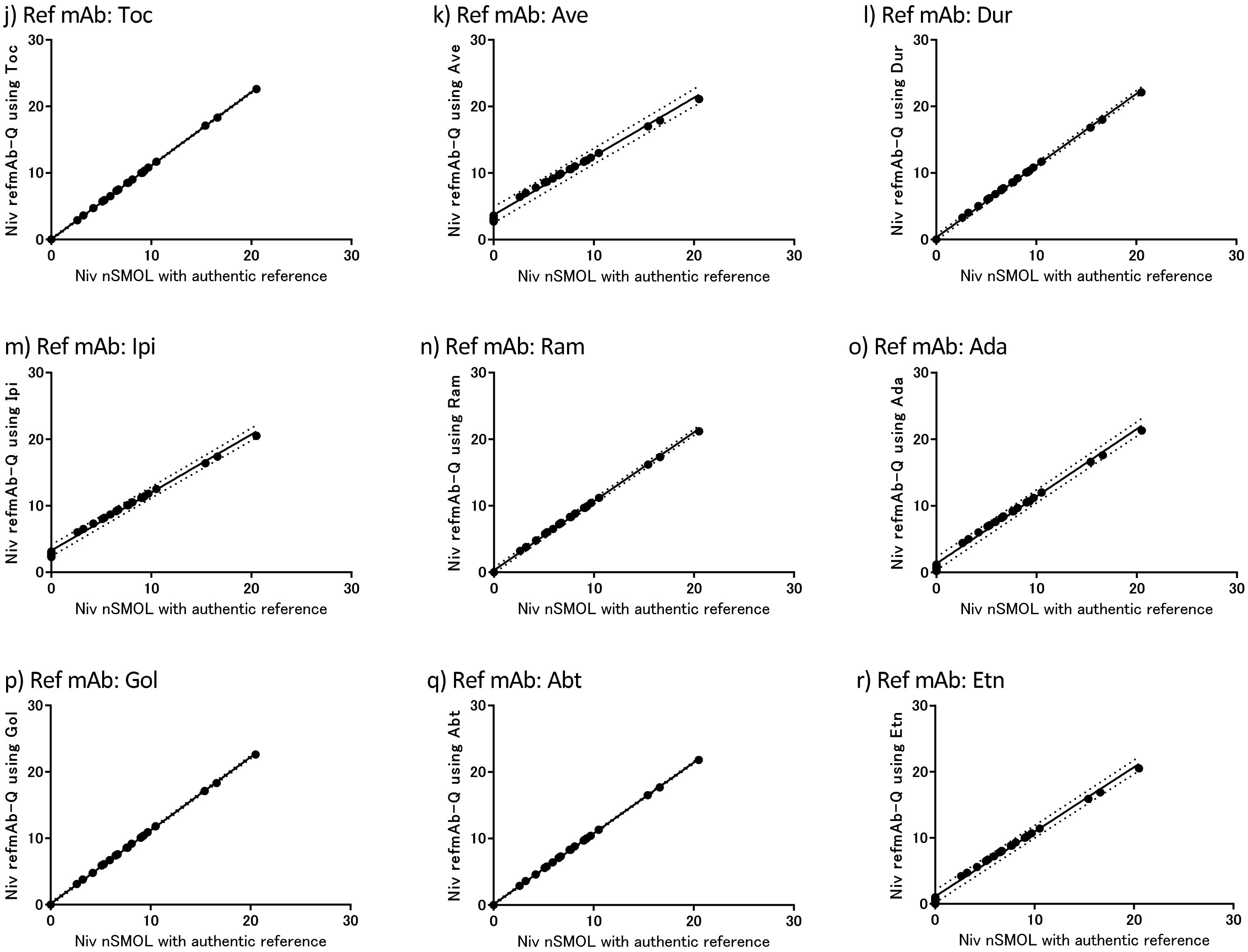
j) Ref mAb: Toc
Niv refmAb-Q using Toc
k) Ref mAb: Ave
Niv refmAb-Q using Ave
l) Ref mAb: Dur
Niv refmAb-Q using Dur
m) Ref mAb: Ipi
Niv refmAb-Q using Ipi
n) Ref mAb: Ram
Niv refmAb-Q using Ram
o) Ref mAb: Ada
Niv refmAb-Q using Ada
p) Ref mAb: Gol
Niv refmAb-Q using Gol
q) Ref mAb: Abt
Niv refmAb-Q using Abt
r) Ref mAb: Etn
Niv refmAb-Q using Etn
Niv nSMOL with authentic reference
0
10
20
30

**Figure S9. Pearson correlation analyses of nivolumab data between nSMOL assay data with the authentic reference and refmAb-Q nSMOL assay data with the indicated reference antibody for clinical serum samples from patients treated with ipilimumab plus nivolumab combination therapy (n =27).** The reference antibody used was a) Bre, b) Cet, c) Rit, d) Ifx, e) Atz, f) Bev, g) Pem, h) Ecu, i) Mep, j) Toc, k) Ave, l) Dur, m) Ipi, n) Ram, o) Ada, p) Gol, q) Abt, or r) Etn, respectively. The dashed line shows the 95% confidence interval. The coefficients are summarized in Table S4d.

**Table S1. MRM transition of each signature peptide of mAbs and Fc-fusion proteins.**

| mAb/Fc-fusion protein* | Signature peptide | MRM transition | Valency of parent ion | Ion series of fragment ion |
|---|---|---|---|---|
| Brentuximab vedotin [o)] | VLIYAASNLESGIPAR | 837.55>343.10 | +2 | y3+ |
| Cetuximab [a)] | SQVFFK | 378.20>540.30 | +2 | y4+ |
| Rituximab [b)] | ASGYTFTSYNMHWVK | 598.05>817.45 | +3 | y13++ |
| Infliximab [c, d)] | SINSATHYAESVK | 469.45>603.90 | +3 | y11++ |
| Atezolizumab [o)] | RHWPGGFDYWGQGTLVTVS SASTK | 660.10>780.30 | +4 | y8+ |
| Bevacizumab [e, f, g)] | FTFSLDTSK | 523.40>797.20 | +2 | y7+ |
| Pembrolizumab [h)] | LLIYLASYLESGVPAR | 882.60>343.20 | +2 | y3+ |
| Trastuzumab [f, i, j)] | IYPTNGYTR | 542.80>404.70 | +2 | y7++ |
| Eculizumab [d)] | LLIYGATNLADGVPSR | 830.45>515.10 | +2 | y5+ |
| Mepolizumab [d)] | DPPSSLLR | 442.70>672.40 | +2 | y6+ |
| Tocilizumab [d, k)] | VTMLR | 310.10>520.20 | +2 | y4+ |
| Avelumab [o)] | LGTVTTVDYWGQGTLVTVSS ASTK | 824.75>780.30 | +3 | y8+ |
| Durvalumab [o)] | VSSSYLAWYQQKPGQAPR | 689.35>625.25 | +3 | y6+ |
| Ipilimumab [p)] | TGWLGPFDYWGQGTLVTVSS ASTK | 853.50>780.40 | +3 | y8+ |
| Nivolumab [f, l, m)] | ASGITFSNSGMHWVR | 550.75>661.50 | +3 | y11++ |
| Ramucirumab [o)] | AFPPTFGGGTK | 540.25>431.30 | +2 | y9++ |
| Adalimumab [d)] | APYTFGQGTK | 535.30>901.40 | +2 | y8+ |
| Golimumab [d)] | SNWPPFTFGPGTK | 718.35>524.80 | +2 | y10++ |
| Abatacept [d, n)] | MHVAQPAVVLASSR | 489.25>420.20 | +3 | y4+ |
| Etanercept [d, n)] | VFCTK | 299.15>498.05 | +2 | y4+ |
| P14R IS | PPPPPPPPPPPPPPR | 512.10>292.30 | +3 | b3+ |

* Assay methods and conditions of each mAbs are available in the following publications. Only DOI numbers are listed below.

a) 10.4155/bio-2016-0018; b) 10.1248/bpb.b16-00230; c) 10.2174/1389201019666180703093517; d) 10.1016/j.jim.2019.06.014; e) 10.1016/j.dmpk.2015.11.004; f) 10.1016/j.ab.2017.11.002; g) 10.1208/s12248-019-0369-z; h) 10.1136/jitc-2021-002371; i) 10.1039/C5AY01588J; j) 10.1016/j.jpba.2017.06.032; k) 10.1016/j.jpba.2018.11.019; l) 10.1016/j.jchromb.2016.04.038; m) 10.1016/j.jchromb.2020.122489; n) 10.1002/prp2.422; o) 10.1007/978-1-0716-1450-1_11; p) 10.1136/jitc-2021-002663

**Table S2. Comparison between ELISA and refmAb-Q nSMOL for detection of mAbs.**

| | ELISA | refmAb-Q nSMOL platform |
|---|---|---|
| Detection principle | Absorbance, fluorescence, chemiluminescence, radiation | Triple quadrupole LC-MS/MS |
| Capture/detection Ab | Direct ELISA requires an idiotypic Ab for capture and anti-human Ab for detection, indirect ELISA uses target molecule with potential cross reactivity or masking by anti-drug Ab | No |
| Reagents | Sandwich, detection Ab,<br>Buffers and reagents | Protein A resin,<br>Nanoparticle trypsin beads<br>Buffers, column, and solvents |
| Sample volume | 50-100 μL | 5 μL |
| Diluent | Requires pH optimization and protein additive for blocking | PBS |
| Cross reactivity | Dependent on Abs, biological samples, and animals | No |
| Reaction selectivity and specificity | Cross-reactivity can be an issue due to indirect detection | By signature peptide structure (physicochemical),<br>Direct structure determination |
| Assay interference from anti-drug antibodies | Interfered | No* |
| Assay development time | Several months | Within a week, can be shorter if antibody sequence is available |
| Internal standard | Individual Ab | Universal one peptide (P14R) |
| Calibrant | Unique authentic | One reference protein |
| Linearity | Narrow, sigmoid | Wide, linear |
| Intra-assay error | Typically within 20-25% | Within 5-15% |
| Multiplexed | Requires special arrangement (Luminex-type assay) | Compatible |
| Operational expertise | Basic wet lab skills | HPLC and MS expertise |
| Preparation time | 3-6 hours | 5-7 hours |
| Automation | Compatible | Compatible |

* Iwamoto N et.al., Antibody drug quantitation in coexistence with anti-drug antibodies on nSMOL bioanalysis. DOI: 10.1016/j.ab.2017.11.002

**Table S3. Signature peptide sequences identified by 18-plex monoclonal antibody analysis and Peaks DB search scores using refmAb-Q nSMOL coupled with Q-TOF-MS analysis.** Peptide and protein score is -10*Log(P), where P is the probability that the observed match is a random event. Peptide score P means the peptide spectrum match (PSM) and greater than 25 are within the threshold of 1% false discovery rate (FDR) in this DB search. Protein scores greater than 20 are of relatively high confidence.

| Antibody | Signature peptide | Region | Peptide score | HC protein score | LC protein score |
|---|---|---|---|---|---|
| Brentuximab vedotin | VLIYAASNLESGIPAR | L-CDR2 | 77.4 | 288 | 273 |
| Cetuximab | SQVFFK | H-CDR2 | 95.0 | 280 | 267 |
| Rituximab | ASGYTFTSYNMHWVK | H-CDR1 | 62.7 | 286 | 267 |
| Infliximab | SINSATHYAESVK | H-CDR2 | 72.4 | 312 | 268 |
| Atezolizumab | RHWPGGFDYWGQGTLVT VSSASTK | H-CDR3 | 76.5 | 304 | 267 |
| Bevacizumab | FTFSLDTSK | H-CDR2 | 42.9 | 296 | 188 |
| Pembrolizumab | LLIYLASYLESGVPAR | L-CDR2 | 73.9 | 295 | 276 |
| Trastuzumab | IYPTNGYTR | H-CDR2 | 45.7 | 295 | 269 |
| Eculizumab | LLIYGATNLADGVPSR | L-CDR2 | 70.0 | 272 | 261 |
| Mepolizumab | DPPSSLLR | H-CDR3 | 41.2 | 274 | 261 |
| Tocilizumab | VTMLR | H-CDR2 | 71.0 | 276 | 260 |
| Avelumab | LGTVTTVDYWGQGTLVT VSSASTK | H-CDR3 | 73.0 | 286 | 147 |
| Durvalumab | VSSSYLAWYQQKPGQAPR | L-CDR1 | 50.0 | 297 | 280 |
| Ipilimumab | TGWLGPFDYWGQGTLVT VSSASTK | H-CDR3 | 85.8 | 305 | 280 |
| Nivolumab | ASGITFSNSGMHWVR | H-CDR1 | 54.9 | 277 | 267 |
| Ramucirumab | AFPPTFGGGTK | L-CDR3 | 54.4 | 302 | 269 |
| Adalimumab | APYTFGQGTK | L-CDR3 | 44.5 | 294 | 270 |
| Golimumab | SNWPPFTFGPGTK | L-CDR3 | 53.0 | 295 | 272 |

**Table S4.** A) Pearson correlation analyses for clinical samples of ipilimumab (n = 71) between refmAb-Q nSMOL with the indicated antibody as the universal reference standard and original data obtained by the conventional nSMOL assay with the authentic reference standard.

| Reference mAb | Goodness of fit $r^2$ | Best fit slope |
|---|---|---|
| Brenzuximab vedotin | 0.998 | 0.983 ± 0.001655 |
| Cetuximab | 0.9993 | 1.08 ± 0.003505 |
| Rituximab | 0.9997 | 1.042 ± 0.002249 |
| Infliximab | 1 | 1.081 ± 0.0004027 |
| Atezolizumab | 0.9999 | 1.074 ± 0.0009812 |
| Bevacizumab | 0.9999 | 1.016 ± 0.0009297 |
| Pembrolizumab | 1 | 1.066 ± 0.0004455 |
| Trastuzumab | 1 | 1.013 ± 0.0004117 |
| Eculizumab | 1 | 0.9631 ± 0.0004318 |
| Mepolizumab | 1 | 0.9152 ± 0.0003888 |
| Tocilizumab | 0.9995 | 1.099 ± 0.00301 |
| Avelumab | 0.9974 | 1.144 ± 0.007094 |
| Durvalumab | 0.9999 | 1.15 ± 0.001206 |
| Nivolumab | 1 | 1.02 ± 0.0004209 |
| Ramucirumab | 0.9999 | 1.071 ± 0.00112 |
| Adalimumab | 1 | 1.004 ± 0.0003909 |
| Golimumab | 0.999 | 1.061 ± 0.001055 |
| Abatacept | 1 | 1.058 ± 0.0003952 |
| Etanercept | 1 | 1.064 ± 0.0005331 |

**Table S4.** B) Pearson correlation analyses for clinical samples of pembrolizumab (n = 74) between refmAb-Q nSMOL with the indicated antibody as the universal reference standard and data obtained by the conventional nSMOL assay with the authentic reference standard.

| Reference mAb | Goodness of fit $r^2$ | Best fit slope |
|---|---|---|
| Brenzuximab vedotin | 1 | 1.059 ± 0.0003607 |
| Cetuximab | 1 | 1.142 ± 0.0003428 |
| Rituximab | 1 | 1.194 ± 0.0003085 |
| Infliximab | 1 | 0.9536 ± 0.0003484 |
| Atezolizumab | 1 | 1.077 ± 0.0003537 |
| Bevacizumab | 1 | 1.12 ± 0.0003395 |
| Trastuzumab | 1 | 1.077 ± 0.0005767 |
| Eculizumab | 1 | 1.021 ± 0.000366 |
| Mepolizumab | 0.9999 | 1.322 ± 0.001245 |
| Tocilizumab | 1 | 1.177 ± 0.0003459 |
| Avelumab | 0.9991 | 1.209 ± 0.004242 |
| Durvalumab | 1 | 1.195 ± 0.0003061 |
| Ipilimumab | 1 | 1.089 ± 0.0003753 |
| Nivolumab | 1 | 1.079 ± 0.0003388 |
| Ramucirumab | 1 | 1.129 ± 0.0003513 |
| Adalimumab | 1 | 1.103 ± 0.0003304 |
| Golimumab | 1 | 1.108 ± 0.000402 |
| Abatacept | 1 | 1.071 ± 0.0003572 |
| Etanercept | 1 | 0.9932 ± 0.0003678 |

**Table S4.** C) Pearson correlation analysis for clinical samples of ipilimumab (n = 27) in the combination therapy treated with ipilimumab and nivolumab between refmAb-Q nSMOL with the indicated antibody as the universal reference standard and original data obtained by the conventional nSMOL assay with the authentic reference standard.

| Reference mAb | Goodness of fit $r^2$ | Best fit slope |
|---|---|---|
| Brenzuximab vedotin | 0.9974 | 1.003 ± 0.01025 |
| Cetuximab | 0.9982 | 0.9829 ± 0.008363 |
| Rituximab | 0.9942 | 1.11 ± 0.01692 |
| Infliximab | 0.9979 | 0.9955 ± 0.00924 |
| Atezolizumab | 0.9944 | 1.069 ± 0.01603 |
| Bevacizumab | 0.9944 | 1.105 ± 0.0166 |
| Pembrolizumab | 0.9977 | 1.002 ± 0.009586 |
| Trastuzumab | 0.9955 | 1.085 ± 0.01454 |
| Eculizumab | 0.9851 | 1.038 ± 0.02556 |
| Mepolizumab | 0.9929 | 1.083 ± 0.01836 |
| Tocilizumab | 0.9921 | 1.162 ± 0.02073 |
| Avelumab | 0.9982 | 0.8713 ± 0.00747 |
| Durvalumab | 0.994 | 1.121 ± 0.01739 |
| Nivolumab | 0.9962 | 1.065 ± 0.01316 |
| Ramucirumab | 0.994 | 1.071 ± 0.01658 |
| Adalimumab | 0.9981 | 1.031 ± 0.008942 |
| Golimumab | 0.9928 | 1.156 ± 0.01964 |
| Abatacept | 0.9923 | 1.051 ± 0.01857 |
| Etanercept | 0.9982 | 1.003 ± 0.008425 |

**Table S4.** D) Pearson correlation analysis for clinical samples of nivolumab (n = 27) in the combination therapy treated with ipilimumab and nivolumab between refmAb-Q nSMOL with the indicated antibody as the universal reference standard and original data obtained by the conventional nSMOL assay with the authentic reference standard.

| Reference antibody | Goodness of fit $r^2$ | Best fit slope |
|---|---|---|
| Brenzuximab vedotin | 0.9932 | 0.9796 ± 0.01622 |
| Cetuximab | 0.9928 | 0.9595 ± 0.01637 |
| Rituximab | 0.9985 | 1.068 ± 0.008222 |
| Infliximab | 0.9929 | 0.9741 ± 0.01649 |
| Atezolizumab | 0.9982 | 1.051 ± 0.00897 |
| Bevacizumab | 0.9984 | 1.07 ± 0.008463 |
| Pembrolizumab | 0.9931 | 0.9759 ± 0.01628 |
| Trastuzumab | 0.9966 | 1.055 ± 0.01228 |
| Eculizumab | 0.9921 | 0.9981 ± 0.01783 |
| Mepolizumab | 0.9996 | 1.038 ± 0.004004 |
| Tocilizumab | 0.9999 | 1.105 ± 0.002026 |
| Avelumab | 0.9919 | 0.88 ± 0.01586 |
| Durvalumab | 0.9987 | 1.08 ± 0.007665 |
| Ipilimumab | 0.9923 | 0.873 ± 0.01542 |
| Ramucirumab | 0.9987 | 1.037 ± 0.007558 |
| Adalimumab | 0.9924 | 1.011 ± 0.01768 |
| Golimumab | 0.9997 | 1.104 ± 0.003991 |
| Abatacept | 0.9998 | 1.065 ± 0.002897 |
| Etanercept | 0.9922 | 0.9792 ± 0.01732 |
